# CSF1R-dependent macrophages control B cell development and function in the chicken immune system

**DOI:** 10.64898/2026.03.16.712250

**Authors:** Zhiguang Wu, Rakhi Harne, Alewo Idoko-Akoh, Francesca Foschi, Simone L. Meddle, Joni Macdonald, Barbara Shih, Mike Mcgrew, David A. Hume, Adam Balic

**Author notes:** The authors contributed equally (Zhiguang Wu, Rakhi Harne). Joint senior authors (David A. Hume, Adam Balic).

## Abstract

Acquired immunity in mammals depends upon capture and presentation of antigens by specialised macrophage populations in splenic marginal zone and lymph node sinuses and follicular dendritic cells (FDC) within germinal centres. Cells referred to as FDC in chickens express CSF1R, the receptor for macrophage colony-stimulating factor (CSF1) and IL34. We utilised single cell RNA-seq on CSF1R^+^ cells from chicken spleen to identify monocytes and two distinct populations of macrophages. TIMD4/C1Q/MAFB^+^ macrophages were enriched for expression of genes involved in iron metabolism. A MARCO/VSIG4^+^ population expressed SPIC, a transcription factor associated with red pulp macrophages in mammals but also expressed receptors (CR2) and trophic factors (TNFSF13, CXCL13) associated with mammalian FDC. SPIC^+^ cells were located within follicles in spleen, caecal tonsil and bursa.

We generated a *CSF1R* knockout in the chicken germ line. Mutant birds lack macrophages in the embryo. They were indistinguishable from wild type at hatch and behaved and fed normally but from day 5-6 post hatch they failed to thrive. Loss of CSF1R function in hatchlings led to monocytopenia and granulocytosis and the loss of macrophage subpopulations in lymphoid organs. Consistent with their expression of B cell trophic factors, the loss of follicular macrophages in the bursa was associated with involution and severe B cell deficiency in the circulation and spleen.

In summary, lymphoid tissues of chickens contain specialised macrophage populations with distinct expression profiles. The details of regulation by CSF1R, specialised functions and underlying transcriptional regulation are quite different between birds and mammals.

## Introduction

In the mammalian innate immune system, two hematopoietic growth factors, macrophage colony-stimulating factor (CSF1) and interleukin 34 (IL34), control the proliferation, differentiation and development of cells of the mononuclear phagocyte lineage acting through a common receptor, CSF1R (1, 2). The receptor, both of its ligands and a macrophage-specific super-enhancer within the *CSF1R* locus (3–6) are conserved in birds. The extracellular domain of CSF1R and one its ligands, CSF1, has apparently been subjected to positive selection during avian evolution suggesting pathogen selection (5). This may be related to the unique expression of CSF1R on antigen-capturing cells in follicle-associated epithelium of bursa and lymphoid follicles (7). We have previously generated a series of resources to study the mononuclear phagocyte system of chickens including *CSF1R* reporter transgenes (8, 9), recombinant CSF1-Fc (10), CSF2 and XCL1 (11) and antibodies against CSF1R (4), TIMD4 (8, 9), CSF1 (10) and FLT3 (11). These resources enabled the identification of subpopulations of antigen-presenting cells with similarities and differences to the cDC1 subset of mammalian dendritic cells (9, 12).

Mammalian secondary lymphoid organs contain multiple specialised macrophage populations distinguished by surface markers and transcription regulators. In the spleen and lymph nodes macrophages associated with the marginal zone and subcapsular and medullary sinuses respectively are adapted to capture and present particulate antigens (13, 14). In both mice and rats these populations are uniquely-dependent upon local production of CSF1 (15, 16). By contrast, macrophages of splenic red pulp, which function in erythrophagocytosis, are relatively CSF1/CSF1R-independent (15, 17). Birds do not have lymph nodes. The development and antigen-mediated affinity evolution of B lymphocytes in birds occurs primarily in the bursa of Fabricius and in germinal centres in the cecal tonsil and spleen (18). These tissues contain abundant resident populations expressing CSF1R (4, 7, 8), variously referred to as bursal secretory dendritic cells (BSDC) and follicular dendritic cells (FDC) based upon their morphology (19, 20). CSF1 is highly-expressed in the splenic ellipsoid, the main site of phagocytic clearance of blood-borne particles (10). In the bursa, CSF1 expression is concentrated around the corticomedullary junction which also contains TIMD4^+^ macrophages (10). Administration of a neutralising anti-CSF1 antibody to hatchling birds led to complete depletion of CSF1R^+^ macrophages in the liver and partial depletion in the bursa and caecal tonsil but had no detectable effect in the spleen (10). The function of CSF1R^+^ macrophages in avian immune tissues and their relationship to counterparts in mammalian systems has not yet been determined. Here we apply single cell RNA sequencing and in situ localisation of mRNA markers to identify functional subpopulations of chicken lymphoid tissue macrophages. The results highlight the distinction between mammalian FDC which are of mesenchymal origin(21–23) and avian FDC which are CSF1R^+^ and macrophage-like.

The original descriptions of phagocyte development in the yolk sac were carried out in the chick. Using a *CSF1R* reporter transgene we described the appearance and function of yolk sac derived phagocytes and their ability to proliferate and migrate to populate the embryo (3, 8). Unlike rodents, which are underdeveloped at birth compared to human infants, chickens at hatch are precocial. Recent advances in genome editing via primordial germ cells (24, 25) have enabled the generation of targeted mutations in chickens (12, 26–28). To determine the function of CSF1R signaling in avian macrophage development we describe the generation and functional characterisation of a null mutation in the chicken *CSF1R* gene.

## Materials and Methods

All birds were obtained from the National Avian Research Facility (NARF) at The Roslin Institute, The University of Edinburgh. Chicks were hatched and housed in premises licensed under a UK Home Office Establishment License in full compliance with the Animals (Scientific Procedures) Act 1986 and the Code of Practice for Housing and Care of Animals Bred, Supplied or Used for Scientific Purposes. Chicken experiments were conducted under UK Home Office licence PP9565661 and PCD70CB48 and PP3522089 and approved by the Roslin Institute Animal Welfare and Ethical Review Board Committee.

### Generation of a CSF1R knockout in the chick

To generate null mutations in the chick *CSF1R* locus we used a CRISPR-Cas9-based targeting approach in primordial germ cells (PGC) and sterile surrogate hosts as described in detail by Ioanidis *et al*. (24). The process is illustrated in **Figure 1A**. We simultaneously delivered a high fidelity CRISPR/Cas9 vector and guide RNAs into *in vitro* propagated male PGC (25). Guide RNAs were designed to delete the exon encoding the transmembrane domain of CSF1R. To prepare gRNAs for delivery into PGCs culture, the two gRNAs were cloned separately into the SpCas9 nuclease encoding vector, pSpCas9(BB)-2A-Puro or (PX459) V2.0 as described previously (29) and co-transfected into PGC using lipofectamine. Genotyping of selected pools confirmed homozygous deletion of the targeted sequence. Single cell cloning generated PGC with the expected homozygous 77bp deletion leading to both removal of the transmembrane domain and a frame shift as shown in **Figure 1B**. Targeted PGCs were injected into transgenic surrogate host chicken embryos with induced deletion of the DDX4 gene (30) so that the only gametes that develop are derived from donor PGCs. Female G0 founder birds were maintained to reach sexual maturity (approximately six months). All the male G0 progeny were culled. G0 hens were bred to homozygous *CSF1R*-EGFP reporter transgenic males (8) via artificial insemination to generate G1 *CSF1R* mutant heterozygotes also containing the *CSF1R*-EGFP transgene. The G1 progeny were then grown to sexual maturity and interbred to produce homozygous mutants. There was a formal possibility that the mutant allele could generate a truncated CSF1R protein with dominant inhibitory activity. However, heterozygous mutant birds grew normally and were fertile, and there was no detectable effect on expression of the *CSF1R*-EGFP transgene.

**Figure 1.**
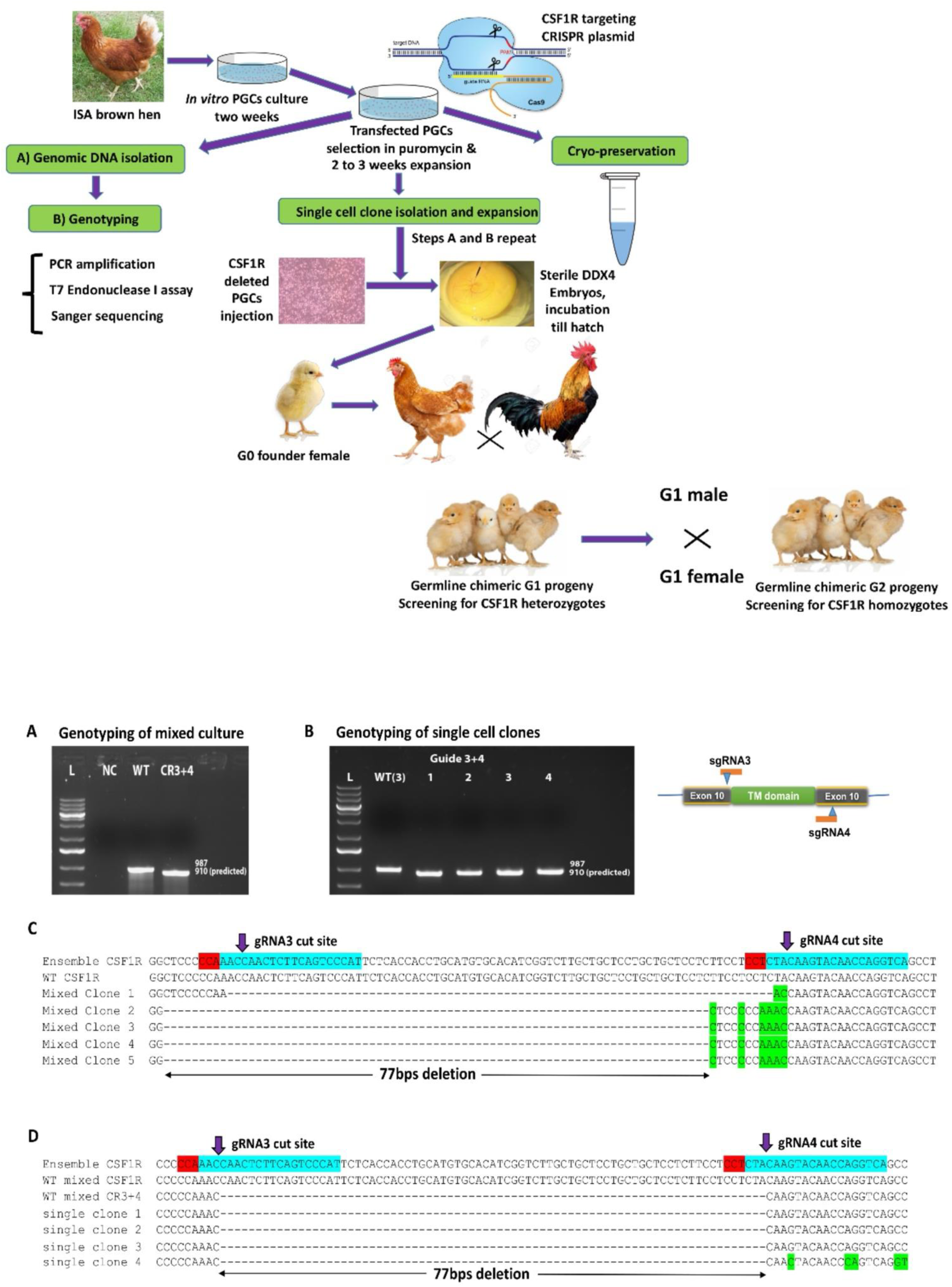
Generation of the *CSF1R* knockout via primordial germ cells. The upper panel shows the overall strategy for the generation of the null *CSF1R* allele using CRISPR-CAS9 in primordial germ cells (PGC). The lower panels show the confirmation of the deletion in PGC by PCR and Sanger sequencing of DNA from mixed and cloned PGC co-transfected with the gRNA pair indicated. Arrows indicate the cleavage sites, CRISPR guides are highlighted in blue and PAM sequence in red.

### Flow cytometry

Blood leukocytes, bone marrow cells and splenocytes were isolated from chicks as described previously (12). Single-cell suspensions from tissues or blood were prepared and resuspended in cold FACS buffer (PBS, 1.0% BSA (w/v) and 0.05% sodium azide (w/v); Sigma-Aldrich) and placed on ice for 10 min. Cells were then incubated with primary antibody in FACS buffer for 30 min on ice in the dark. If required, cells were washed and incubated with secondary antibodies for 20 min on ice in the dark. Cells were then washed three times, resuspended in cold FACS buffer and stained with SYTOX® Blue Dead Cell Stain (Invitrogen; 1.0mM stock, 1/4000 dilution) prior to analysis with a LRSFortessa flow cytometer (BD Biosciences). Data were analysed using FlowJo V10 software. Dead cells were excluded by SYTOX® Blue staining and doublets were then discriminated based on signal processing (FSC-A/W). Fluorescence minus one (FMO) controls were used to confirm gating strategies. As described previously, mouse monoclonal antibodies against CD45, BU.1 (CD45R), CD3, KUL01, MHCII were obtained from Southern Biotech, anti-GFP from ThermoFisher Scientific, UK and anti-FLT3 (ROSAV184) was produced in house (12).

### Immunofluorescent staining

Tissue samples were trimmed into 1.0 cm^2^ blocks and fixed overnight at 4°C in 4% paraformaldehyde (PFA; Sigma-Aldrich)/phosphate buffered saline (PBS; Sigma-Aldrich). Samples were removed from PFA/PBS and placed in 10% sucrose (w/v; Sigma-Aldrich)/PBS at 4°C overnight. Samples were then placed in 15/20/25/30% sucrose/PBS (w/v) for 24 h at 4°C for each sucrose concentration. Tissue samples were embedded in Cellpath™ OCT embedding matrix (Fisher Scientific UK Ltd, Loughborough, UK) and snap-frozen at −80°C for two hours. 10µm sections were cut onto Superfrost Plus slides (Menzel-Gläser, Braunschweig, Germany) and air-dried for 1h at RT. Primary antibodies were rabbit anti-lysozyme (ab391, Abcam), anti-CD11 (8F2) ((31) a gift from Dr. S. Hartle), anti-TIMD4 (clone JH9, (9)), anti-BU-1 (AV20, Southern Biotech), anti-MAFB (HPA005653, Atlas antibodies) and anti-GFP Rabbit Polyclonal Antibody, Alexa Fluor™ 488 (ThermoFisher). All slides were blocked for one hour in 10% normal horse serum (Sigma-Aldrich), 0.1% Triton X-100 (Sigma-Aldrich) in PBS (HST-PBS). All primary antibodies were diluted in blocking reagent (above) and incubated at 4°C overnight, washed for 20 min in PBS, followed by incubation with secondary antibodies for two hours (donkey anti-mouse IgG Alexa Fluor 594, donkey anti-mouse IgG1 Alexa Fluor 594, donkey anti-mouse IgG2a Alexa Fluor 647; Thermo Fisher Scientific) used at 1/300 dilution and mounted in ProLong® Gold Antifade Mountant (Thermo Fisher Scientific). Where appropriate, sections were counterstained with 1 μg/ml 4′, 6′-diamidino-2-phenylindole (DAPI; Sigma-Aldrich) in the final incubation step. Samples were imaged using an inverted confocal microscope (Zeiss LSM710) and images were analysed using Zeiss ZEN 3.1 software.

### In situ hybridization

RNAscope Fluorescent Multiplex V2 Assay (ACD Biosciences) was performed according to the manufacturer’s protocol. *SPIC*^+^ cells were identified using negative-sense RNA probe (1031301-C1 (Gg-SPIC;Bio-Techne, USA) derived from the chicken *SPIC* mRNA sequence (XM_025155792). Negative control tissues were probed with the 310043- RNAscope® Negative Control Probe (Bio-Techne, USA) derived from the Bacillus subtilis strain SMY methylglyoxal synthase (mgsA) gene (EF1915.1). Hybridization was done on 6 mm formalin fixed paraffin sections and detected using Opal Dyes (Akoya Biosciences). After *in situ* hybridization, samples were washed in TBST and immunofluorescence was performed as above. Samples were counterstained with DAPI, mounted in Prolong Gold and imaged using a Zeiss LSM 880 confocal microscope.

### Single cell RNA-seq

Splenocytes were separately prepared from two 20-week-old female *CSF1R*-EGFP transgenic birds. *CSF1R*-EGFP^+^ cells were sorted using a BD FACS Aria IIIu sorter with the target of 5000 cells per sample. Libraries were prepared using a Chromium™ Single Cell 3’ Library & Gel Bead Kit v3 using the 10X Chromium Single Cell RNA Sequencing Platform. The single-cell libraries were sequenced at a depth of 50,000 reads per cell on an Illumina NovaSeq machine at Edinburgh Genomics, The University of Edinburgh. Cell Ranger (version 3.1.0) was used to generate transcriptomic reference index from Chicken reference genome (*Gallus gallus* GRCg6a, Ensembl version 101) through *cellranger mkref*, and gene expression matrix through *cellranger count*. Downstream analyses were performed on R Seurat (version 3.2.2) (32). Cells with low number of features (gene; <200) and outlier counts (unique molecule identifier; <300 or >12,000), or high mitochondrial count (>40% counts from mitochondrial genes) were removed from further analysis. Principal component analysis was performed on the cells using 50 components, and the top 30 components were used for generating a network graph using Graphia (version 2.0) (33) retaining nodes and edges fulfilling the following parameters: r ≥ 0.75, knn = 10, node degree > 5, and component size > 5. The resultant cells remained in the Graphia network graph were retained in the R Seurat analysis, which was subjected to further quality control using DoubletDecon to remove doublets (34). Following quality control, cells from two samples were integrated using *SelectintegrationFeatures, FindIntegrationAnchors* and *IntegrateData* functions in Seurat.

Cell cycle states were labelled with *CellCycleScoring* using gene list organised by Seurat based on Tirosh *et al.* (35). From the splenic CSF1R-EGFP^+^ cells scRNAseq data, macrophages and monocytes (*MAFB^+^ XCR1^-^ JCHAIN^-^* cells) were selected for re-clustering to exclude *XCR1^+^* cDCs and pDCs (12). To avoid clustering artifacts, proliferating cells (clusters high in *TOP2A, PCNA, MCM6, MKI67* and *STMN1*) were also removed from this analysis. Uniform Manifold Approximation and Projection (UMAP) dimension reduction was performed across all cell types and Potential of Heat-diffusion for Affinity-based Transition Embedding (PHATE) (36) was used to further interrogate the populations. The scRNAseq data for the study is available on https://www.ncbi.nlm.nih.gov/ (BioProject : PRJNA996296)

## Results

### Characterisation of chicken lymphoid tissue macrophage populations

The adult chicken spleen contains a population of cDC1-like classical dendritic cells expressing XCR1, FLT3 and other DC markers. Unlike mammalian cDC1, chicken cDC1 also express CSF1R and the *CSF1R-*EGFP transgene (9, 11, 12). The proportion of these cells amongst the *CSF1R-*EGFP splenic cell population increases from around 3% at hatch to 40% in adults (12). Single cell RNA-seq analysis of isolated *CSF1R-*EGFP^+^ adult spleen cells, with DC and proliferating cells excluded, revealed separate clusters as shown in the UMAP projection in **Figure 2A**. Phate trajectory analysis (**Figure 2B**) highlights the clear segregation of three macrophage populations based upon expression of *TIMD4, CSF3R* and *SPIC.* The myeloid transcription factor, *MAFB*, was detected in all of the clusters (**Figure S1A**). The *CSF3R* expressing cells were previously identified as monocytes (12). **Table S1** and **Figure 2C** show the set of transcripts selectively expressed in *TIMD4*^+^, *SPIC*^+^ and *CSF3R^+^*populations. In mammals, *SPIC* is expressed by splenic red pulp macrophages and is essential for the function in erythrophagocytosis and iron homeostasis (37). By contrast, *SLC40A1* (encoding ferriportin), *HMOX1* and *FTH* in the chick are enriched in the *TIMD4* subset. *TIMD4*^+^ cells are also enriched for expression of class II MHC (*BLB1/2, CD74*), *CTSS, C1QA/B/C, HAVCR1 (TIM1), MERTK* and *STAB1* suggesting a function in antigen capture and processing akin to marginal zone macrophages in rodent spleen and sinus macrophages in mammalian lymph nodes. *MMR1L4* (KUL01) was detectable in a subset of TIMD4^+^ cells in the scRNA-seq data (**Figure S1A**) but previous analysis showed the colocation of TIMD4 and KUL01 in the peri-ellipsoid compartment in spleen (9). The *SPIC^+^* population on the other hand expresses genes associated in mammals with follicular dendritic cells and B cell development including *CR2, CXCL13* and *TNFSF13B* (B cell activating factor, BAFF) (22, 38) (**Figure 2D**) as well as *MARCO, VSIG4* and the transcription factor *NR1H3* (Lxr). *Nr1h3* is required for development of marginal zone macrophages in mice (39). In mammals, follicular dendritic cells (FDC) derive from non-hematopoietic vascular mural cells (21, 40). We examined the location of key markers in splenic germinal centres (**Figure 2E, Figure S1B**). The colocalization of IgY immune complexes, CSF1R protein, *CSF1R-*EGFP, *SPIC* mRNA and *MAFB* in germinal centres confirms that a specialised population of CSF1R^+^, SPIC^+^ macrophages in birds likely differentiates to perform the function of FDC.

**Figure 2.**
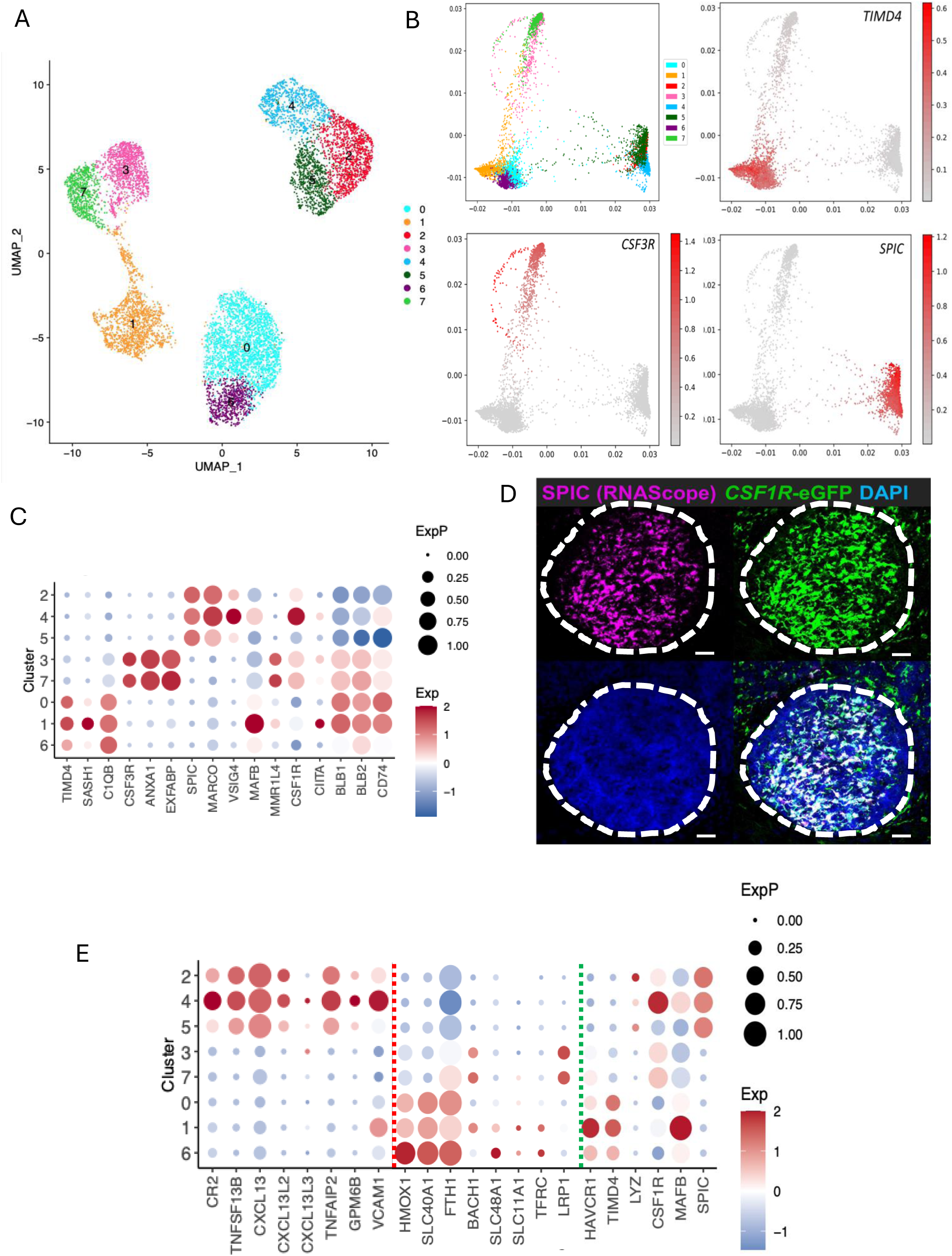
Identification of populations of macrophages in avian spleen using single cell RNA-seq. Unbiased clustering of RNA-sequence data from 16,994 *CSF1R*-EGFP transgene expressing cells derived from the spleens of two 20 week old hens. B) PHATE map of macrophage clusters with feature plots of markers for *TIMD4, CSF3R* and *SPIC*. C) Dotplot of selected macrophage/monocyte associated genes transcript abundance in cell clusters. D) Representative image of combined immunofluorescence staining of EGFP and RNAScope *in situ* hybridisation detection of *SPIC* expression in adult chicken splenic germinal centre. Germinal centre outlined in white dashed line. Scale bars = 50 µm. E) Dotplot of selected genes in SPIC^+^ splenic macrophages (Cluster 2,4,5). Selected genes in left panel are associated with FDC function and in central panel with red pulp macrophage erythrophagocytosis in mammals.

The gene encoding lysozyme, *LYZ*, is highly expressed in chicken spleen and bursa and in isolated macrophages (41) but was detected in only a small subset of *SPIC*^+^ cells in scRNA-seq (**Figure 2C**). Based upon evidence from analysis of mammalian splenic macrophages (42), we speculated that LYZ^+^ macrophages might be selectively lost in the process of tissue disaggregation. To test this proposal, we colocalised *CSF1R-*EGFP, *LYZ*, *SPIC* and *TIMD4* in spleen and extended the analysis of these markers to bursa and cecal tonsil (**Figure 3, Figure S2A/B**). In the spleen, a subset of *SPIC*^+^ cells within the ellipsoid expressed LYZ. A distinct population of LYZ^+^, EGFP^+^ cells form a ring around the ellipsoid endothelial cells, interior to the layer of TIMD4^+^ macrophages described previously (9) (**Figure 3A/B**), reminiscent of the arrangement of separate CD169^+^ and CD209B^+^ macrophages in rodent splenic marginal zone (15). Mammalian germinal centres contain a specialised phagocyte population, tingible body macrophages, that engulf apoptotic B cells. In mice, these cells lack detectable CSF1R expression (43). These cells are also identified as a separate CSF1R-EGFP^low^, TIMD4^+^ population in chicken splenic follicles (**Figure S2C).**

**Figure 3.**
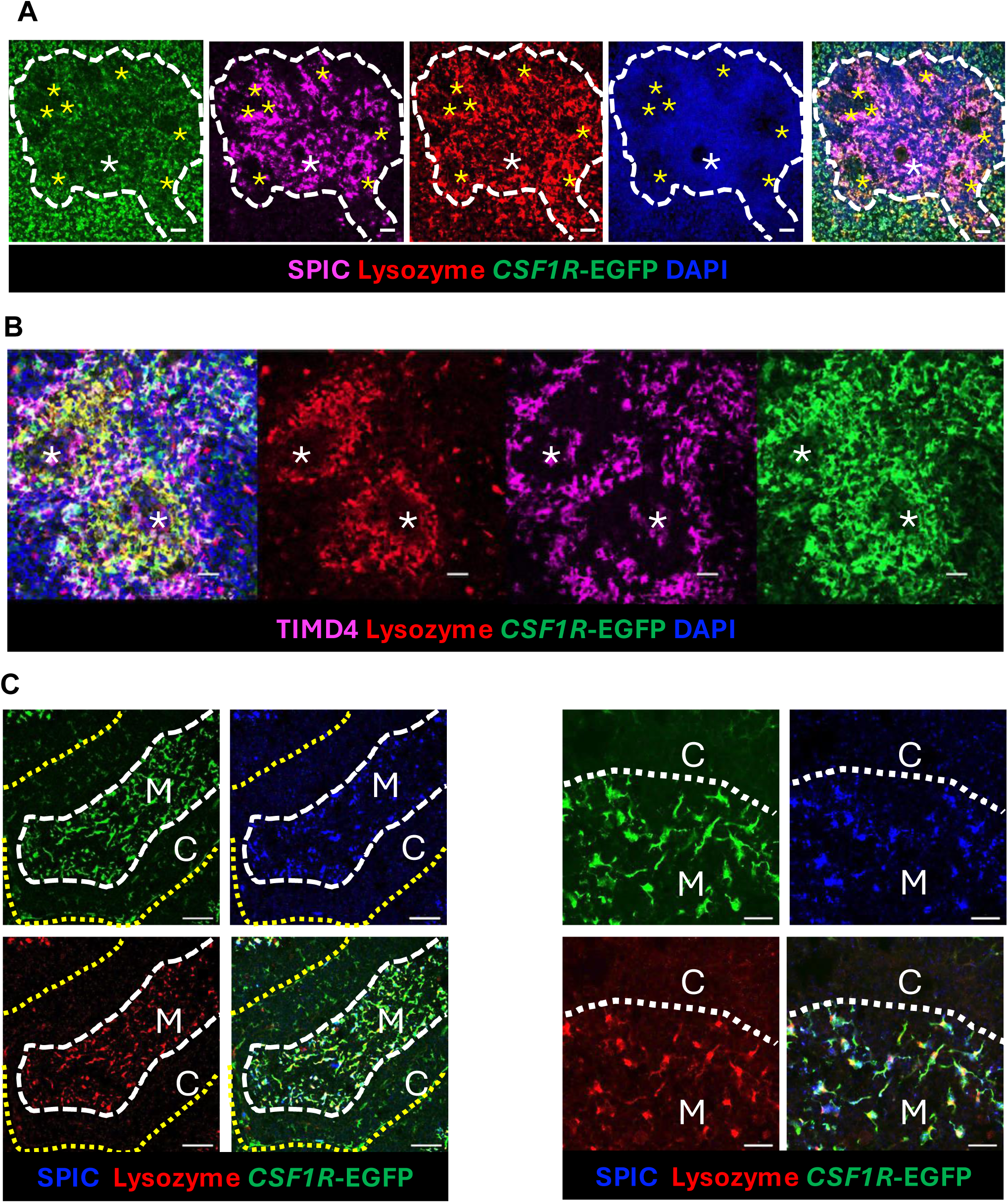
Identification of subpopulations of macrophages in chicken spleen. Representative images of combined detection of EGFP and lysozyme by immunofluorescence and *SPIC*by RNAScope *in situ* hybridisation within the adult splenic ellipsoid (demarcated by the white dashed line). The position of the ellipsoid central artery and penicillary capillaries are show by white and yellow asterix respectively. Scale bars = 20 µm A) Representative images of combined detection of EGFP, Lysozyme and TIMD4 by immunofluorescence within the adult splenic ellipsoid. Note that Lysozyme expressing *CSF1R*-EGFP^+^ cells form a ring around the ellipsoid endothelial cells (white*), surrounded by a more distal layer of TIMD4+ macrophages. Scale bars = 20 µm. C) Representative images of combined detection of EGFP and Lysozyme by immunofluorescence and *SPIC* by RNAScope *in situ* hybridisation within adult bursa of Fabricius. An individual B-cell follicle is outlined by a yellow dotted line. The border between the outer cortex (C) region and the inner medulla (M) is marked by a white dashed line. Scale bars = 50 µm. Panel at right shows a higher magnification. The border between the outer cortex (C) region and the inner medulla (M) is marked by a white dotted line. Scale bars = 10 µm

In the bursa, TIMD4 was previously located in macrophages at the cortico-medullary junction of individual follicles (9). By contrast, stellate EGFP^+^ macrophages in the medulla were SPIC^+^/LYZ^+^ (**Figure 3C**). In the caecal tonsil EGFP^+^ cells in the germinal centre are also SPIC^+^/LYZ^+^. Within the lamina propria LYZ is not detected but a subset of GFP^+^ macrophages express SPIC (**Figure S2B**).

In summary, like mammalian spleen and lymph nodes, chicken lymphoid tissues contain multiple CSF1R^+^ resident macrophage populations defined by markers and location. The unique features of the avian system include the apparent coopting of macrophages to perform the function of FDC and the distinctive expression of the transcription factor, SPIC.

### Analysis of CSF1R-deficient chickens

In order to dissect the functions of CSF1R and CSF1R-dependent macrophages in the chicken immune system, we created a null mutation in the germ line by CRISPR-Cas9 editing (**Figure 1**). To enable analysis of the impact on myeloid populations, the deletion was generated in the *CSF1R*-EGFP reporter line. Macrophages expressing *CSF1R* mRNA or *CSF1R*-reporter transgenes are detected first in the yolk sac at stage HH13 and confined to the lumen of primitive blood vessels (8). In later stages of development, ramified macrophages expressing *CSF1R*-transgene reporter are found throughout the embryo (3, 8). Initial genotyping of *CSF1R*^-/-^ embryos at day 8 of incubation indicated that homozygous mutants were present at the expected Mendelian frequency and were not morphologically distinct (not shown). However, there were no *CSF1R-*EGFP expressing macrophages detectable in *CSF1R*^-/-^(*CSF1RKO*) embryos, including in areas such as the footpad, where they are abundant in *CSF1R*^+/+^ (WT) birds and actively engaged in phagocytosis of dying cells (**Figure S3**). Notably, as reported previously in macrophage-deficient mice (44, 45), there is no apparent defect in development of the digits, indicating that non-macrophage mesenchymal cells can take over the task of apoptotic cell removal. *CSF1RKO* embryos remained macrophage-deficient at day 17 of incubation. Microglia expressing *CSF1R-*EGFP or surface markers CD44 and CD45 were undetectable in brain parenchyma, and TIMD4^+^ macrophages in the meninges were also greatly reduced (**Figure S4A**). Microglia remained undetectable at 3 days post hatch (**Figure S4B**).

Upon heterozygous mating, *CSF1RKO*, heterozygous and WT chicks hatched at expected 1:2:1 Mendelian frequencies and *CSF1RKO* chicks were initially physically indistinguishable from their siblings. Upon histological analysis of the brains, the absence of microglia was not associated with macroscopic anatomical defects (hydrocephalus, callosal dysgenesis, olfactory bulb involution) reported in juvenile *Csf1rko* rodents (46, 47). We showed previously that post-hatch treatment of chicks with a neutralising anti-CSF1 antibody led to complete osteoclast depletion and a mild increase in trabecular bone (10). Unlike *Csf1r* knockout mice and rats, where the growth plate is compromised (48), the long bones were not foreshortened and there was no impact of the mutation on stature. In this respect, the *CSF1RKO* chicken phenotype resembles osteosclerosis in human infants with homozygous recessive *CSF1R* mutation (49, 50). The precocial phenotype in the chicken hatchling is supported by the yolk sac which is internalised prior to hatch and provides a nutrient reserve during the first 3 to 5 days post hatch (51). The body weights of WT and *CSF1RKO* chicks were not significantly different at hatch or the first 6 days post hatch. However, after day 6 post hatch, body weight gain in *CSF1RKO* chicks abruptly ceased and their condition declined rapidly, requiring euthanasia by day 9.

### Selective depletion of monocytes, macrophage and B cells in CSF1RKO chicks

Figure 4 shows FACS analysis of *CSF1R-*EGFP and various immune cell markers in blood of WT, *CSF1R*^+/-^ (HET) and *CSF1RKO* chickens. An 80-90% reduction in *CSF1R-EGFP* monocyte count was confirmed by staining for the monocyte marker, KUL01 (MRC1L-b, gene name *MMR1L4*) (9) (Figure 4A**,B**). Amongst the non-myeloid populations, the relative abundance of thrombocytes (CD41^+^, CD45^+^) and erythrocytes was unaffected. Amongst lymphoid populations, CD3^+^ T cells were similar to WT (not shown). By contrast, in *CSF1RKO* birds, staining for BU.1 or MHCII revealed the almost complete absence of B cells, also confirming the reduced numbers of *CSF1R*-EGFP^+^ monocytes which share these markers with B cells (Figure 4C**,D**).

**Figure 4.**
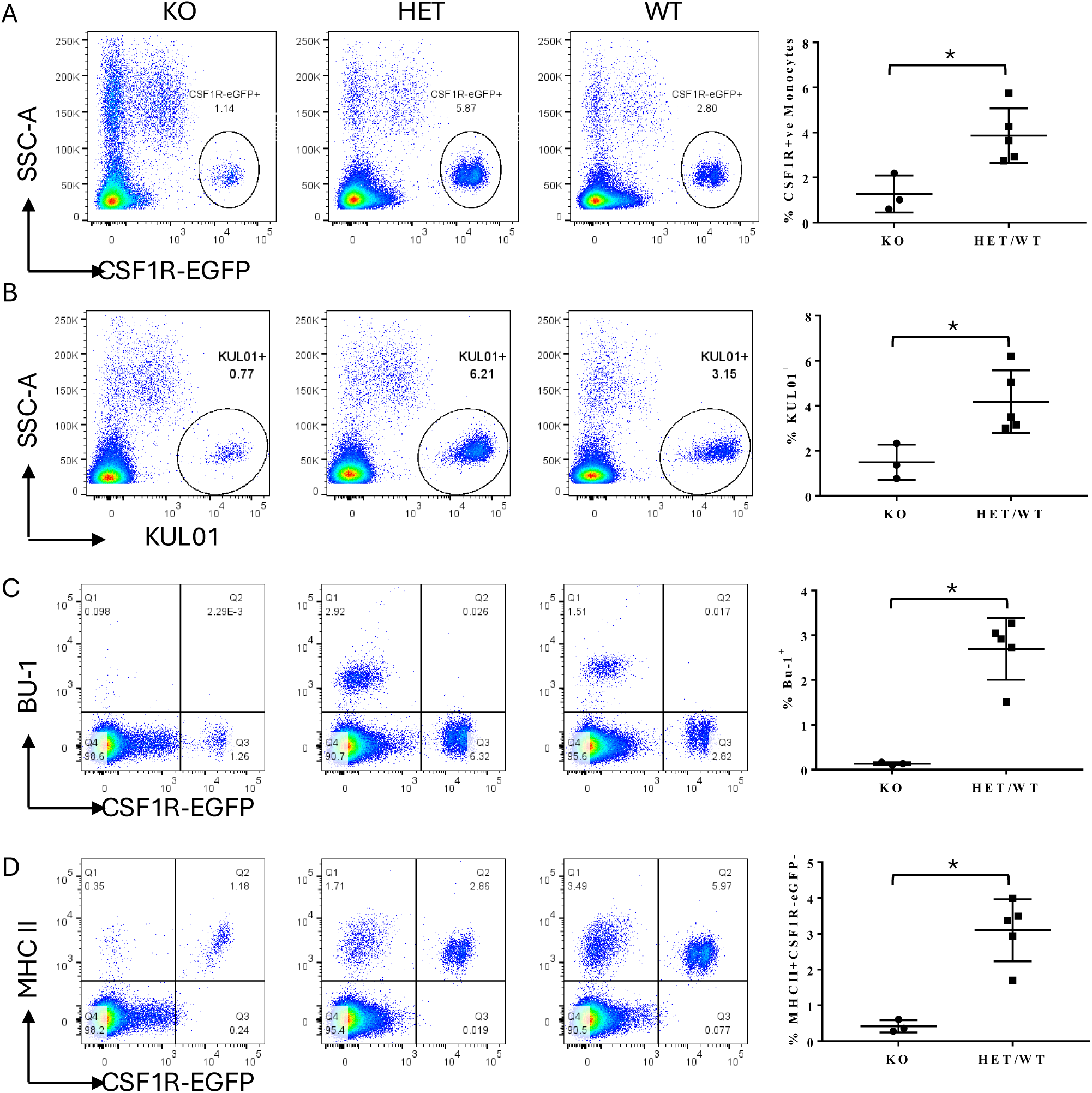
Analysis of blood leukocytes in *CSF1RKO* chicken. Flow cytometric analysis of CSF1R-EGFP expressing monocytes and other immune cell populations in blood from a sibship of WT, *CSF1R*^+/-^ (HET) and *CSF1RKO* chickens at 6 days post hatch. Representative plots are shown for each genotype, n=3 (KO) and n=5 (HET/WT). The frequencies of gated populations were calculated and compared between the KO and HET/WT groups. Statistical analysis was conducted using unpaired non-parametric Mann-Whitney test. Statistical significance was defined as follows: ∗, p < 0·05; ∗∗, p < 0·01; and ∗∗∗, p < 0·001, ns = not significant. The selective loss of monocytes and B cells in *CSF1RKO* chickens was confirmed in an independent cohort.

We showed previously that the ligand, CSF1, is highly-expressed in the peri-ellipsoid region of spleen, coincident with the TIMD4/KULO1^+^ macrophage population, but these cells were resistant to depletion with anti-CSF1 treatment in hatchlings (10). Because of the early mortality of the *CSF1KO* chickens, analysis of the mutation was performed at a stage when WT birds have few cDC1 and germinal centre formation in the spleen is minimal. FACS analysis revealed the almost complete depletion of *CSF1R*-EGFP^+^ cells in *CSF1RKO* spleen, the large majority in the WT co-expressing KUL01 (Figure 5A**, B**). Consistent with the blood profiles, the *CSF1RKO* spleens were almost completely depleted of BU.1^+^ B cells, whereas CD3^+^ T cells were unaffected (Figure 5C-E). By contrast to mammalian macrophages the majority of EGFP^high^ cells in the WT spleen co-expressed FLT3 and FLT3^+^ macrophages were also depleted in the *CSF1RKO* spleens (Figure 5F).

**Figure 5.**
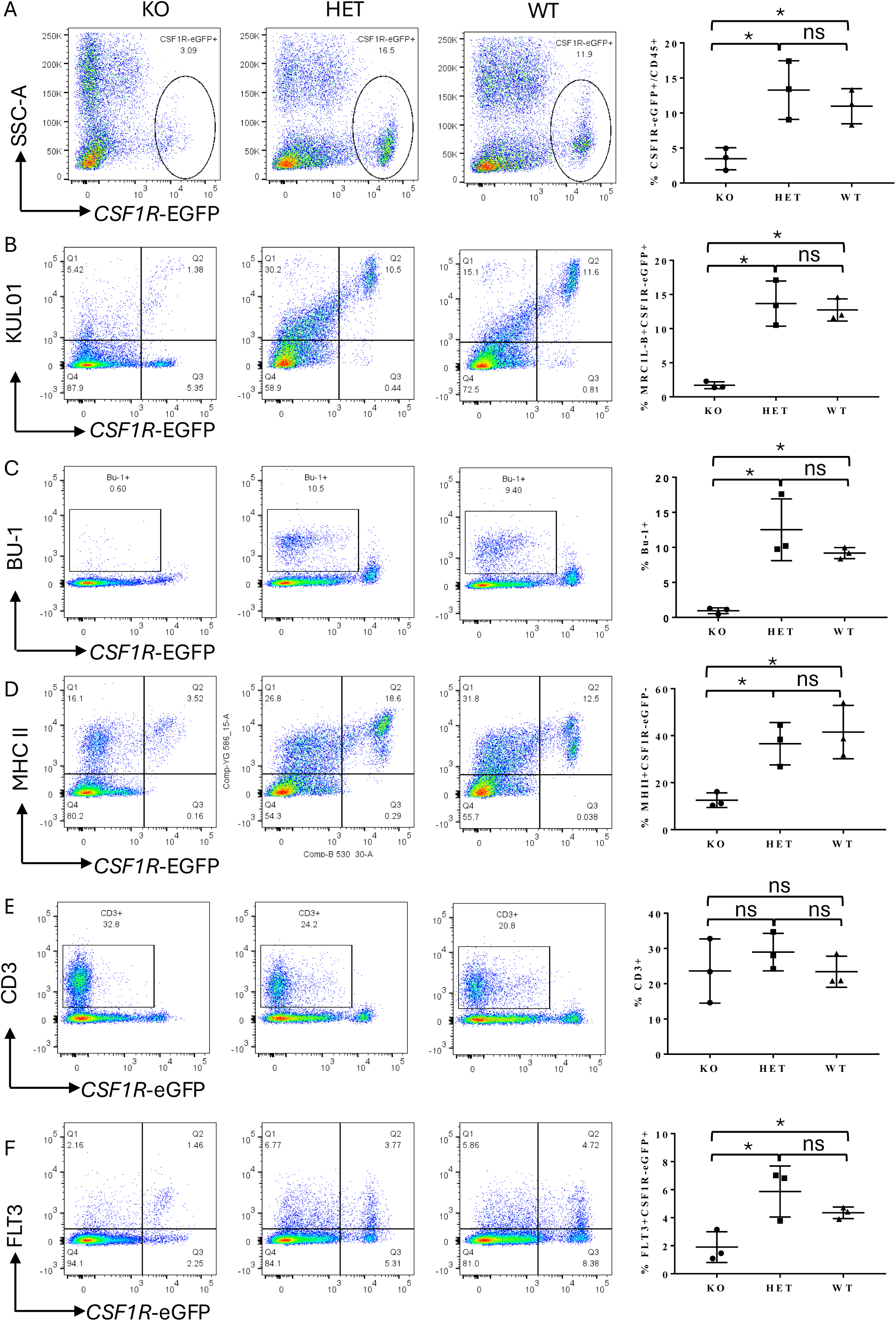
Analysis of splenic leukocyte populations in *CSF1RKO* chicken. Single-cell suspensions of splenocytes were isolated from a sibship of WT, *CSF1R*^+/-^ (HET) and *CSF1RKO* chickens at 3 days post hatch. Panels show representative profiles for the chosen markers with quantitation (N=3) for indicated quadrants. Statistical analysis was conducted using a two-tailed unpaired t-test with Welch’s correction. Statistical significance was defined as follows: ∗, p < 0.05; ∗∗, p < 0.01; ∗∗∗, p < 0.001. The selective loss of monocytes, macrophages and B cells defined by KUL01, BU.1, MHCII and FLT3 in *CSF1RKO* chickens was confirmed in an independent cohort.

The absence of B cells in blood and spleen obviously focusses attention on the bursa of Fabricius, the main site of B lymphopoiesis in birds. CSF1 protein is highly expressed by cells within the corticomedullary junction in the bursa (10). The bursa contains two abundant macrophage populations. Cortical macrophages expressing TIMD4 and KUL01, like Kupffer cells are CSF1R^+^ but do not express the *CSF1R*-EGFP transgene (7). GFP^+^, TIMD4^-^, KUL01^-^follicular macrophages, also called bursal secretory dendritic cells, form an extensive network throughout the medulla (7, 19). Unique to birds, both *CSF1R*-EGFP and CSF1R protein are also expressed in antigen-sampling cells associated with follicle-associated epithelium in the intestinal lymphoid tissues including the bursa (7). During development, bursal follicle bud formation is initiated by infiltration of a blood born inducer population. Subsequently separate B cell and *CSF1R-*EGFP+ precursors enter the bursal mesenchyme (52). Despite the apparent absence of macrophages in the embryo, the bursa appeared to develop normally. Like the mammalian thymus, the avian bursa undergoes involution in adults associated with reduced endocytic activity in the follicle-associated epithelium (53). At a gross level, the bursa of *CSF1RKO* chicks was already substantially involuted at 3 and 6 days post hatch. Figure 6 shows comparative images of WT and *CSF1RKO* bursa with the *CSF1R*-EGFP marker and staining of B cells (BU.1) and CD11b/c. Several important features are evident. Most obviously, the bursa in *CSF1RKO* chicks is grossly B cell deficient, with the majority of follicles having no detectable BU.1^+^ cells. *CSF1R*-EGFP^high^ stellate cells in both the follicle medulla and the cortex, both shown to express high levels of CSF1R by IF localisation (19) are absent in the knockout. By contrast, the knockout had no apparent effect on the EGFP^high^ M cell like cells within the FAE (Figure 6A). The so-called bursal secretory dendritic cells (BDSC) (19), the major myeloid population within the bursal follicles, express CD11b/c (19). Staining for this marker confirmed the complete loss of BDSC but also highlighted the presence of CD11b/c^+^ cells surrounding the follicles in the knockout. These cells are granulocytes, also detected as EGFP^low^ cells (Figure 6)

**Figure 6.**
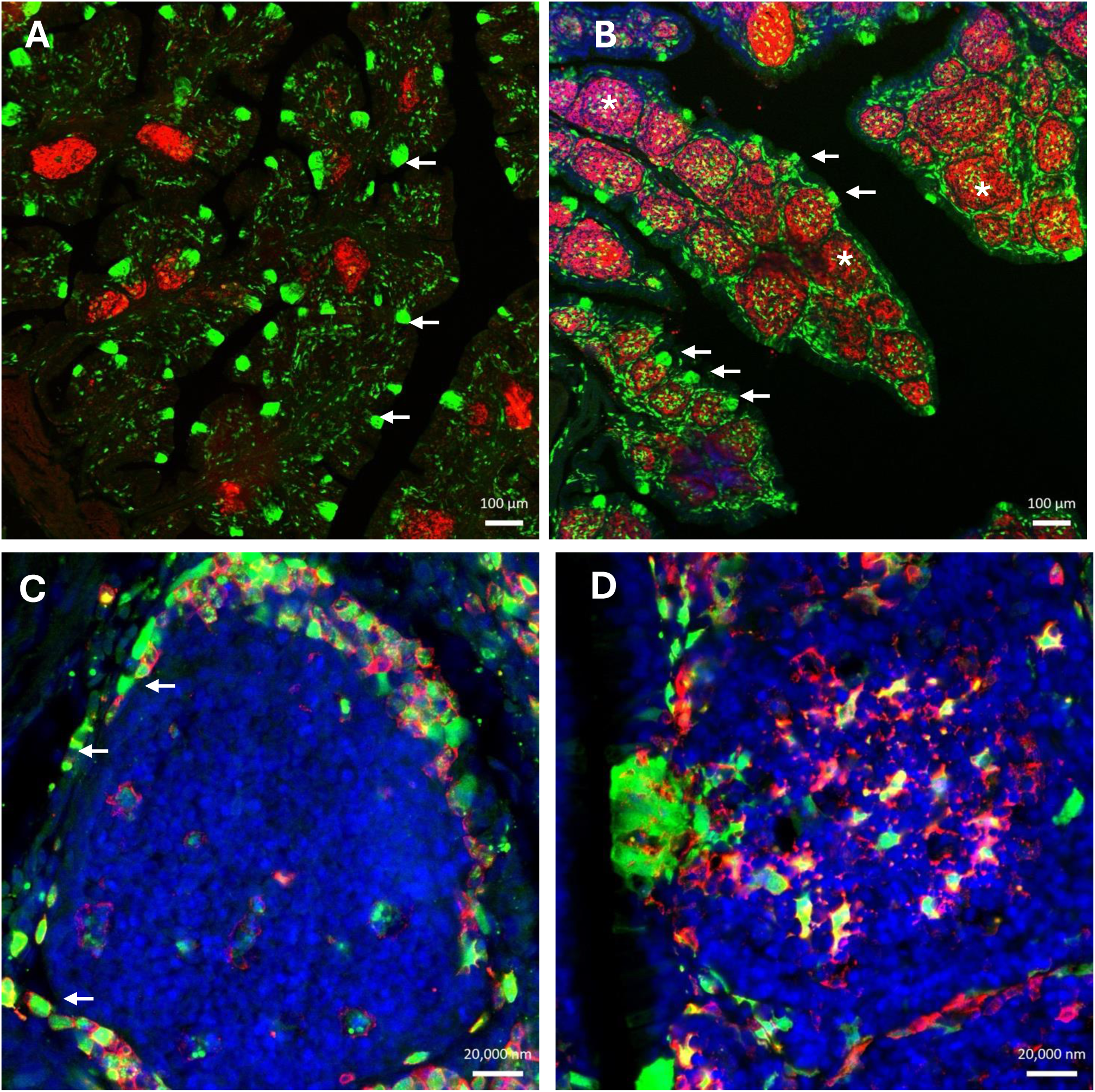
Involution of the bursa of Fabricius in *CSF1RKO* chicken. Panels show representative images of bursa of Fabricius from three-day old *CSF1RKO* (A,C) or WT (B,D) chicks stained for BU-1+ B-cells (Red) and the CSF1R-EGFP transgene (Green) (A,B) or CD11 (Red) and the CSF1R-EGFP transgene (Green) (C,D). Note in A, B that BU.1^+^ cells in intact B-cell follicles (asterix) are depleted in *CSF1RKO* bursa and intrafollicular EGFP+ macrophages are absent. By contrast, M cell-like EGFP^+^ cells within follicle-associated epithelium are retained (arrows). Higher magnification of individual follicles in panels C and D confirms the absence of CD11^+^/EGFP^+^ intrafollicular macrophages in the *CSF1RKO* bursa alongside perifollicular accumulation of CD11^+^/EGFP^+^ granulocytes (arrows). Scale bars = 100 µm (A, B) or 20 µm (C, D).

## Discussion

In this study we have characterised subpopulations of CSF1R-expressing macrophages in chicken immune organs generated and analysed the effect of a homozygous null mutation in the chicken *CSF1R* gene. The outcome of the *CSF1RKO* provides a formal confirmation of our previous work demonstrating that the fundamental biology of CSF1R and its two ligands, CSF1 and IL34, is conserved in birds (3–5, 8, 10–12, 54). Recent studies in zebrafish support the same conclusion in fish, albeit complicated by the duplication of receptor and ligand genes (55, 56). Given the abundance of macrophages in the chicken embryo and their active participation in phagocytic clearance of apoptotic cells (3, 8), the most surprising observation is that the *CSF1RKO* chickens develop and hatch entirely normally despite the gross macrophage deficiency. The same pattern is evident in *CSF1R*^-/-^ rats which are indistinguishable from littermates *in utero* (45) and at birth (48). The fundamental biology of microglia is conserved from birds to mammals (57). Microglia were entirely absent in the *CSF1RKO* chicks (**Figure S4**). The lack of any gross anatomical deficiencies in the brain or behavioural phenotypes supports recent evidence from rodent *Csf1r* mutations that microglia are not entirely essential for normal postnatal development of the brain (58). In both the chicken and the rat, the absence of erythroblastic island (EBI) macrophages in fetal liver, attributed essential functions in erythrocyte development (59), is not associated with any detectable red cell abnormality. Recent studies have suggested that EBI macrophages regulate the balance between erythropoiesis and granulopoiesis (60), which could explain the granulocytosis in *CSF1RKO* chicks that is common to *CSF1R*^-/-^ rats (48) and mice (17).

In *CSF1R*^-/-^ rats there is a severe postnatal decline in somatic growth that is associated with a delay in development of the liver and the loss of circulating insulin-like growth factor 1 (IGF-1). Restoration of wild-type tissue macrophage populations by bone marrow cell transfer at weaning rescues the phenotype (48, 61). In the chicken *CSF1RKO*, the sudden and rapid decline in somatic growth from day 6 post hatch occurs shortly after the resorption of the yolk sac is complete. Chicken yolk contains immunoglobulin and a complex mixture of proteins secreted by the liver in the laying hen and transferred via the oviduct (62). The depletion of this resource in the post hatch chick may in some way mirror the separation of the rodent embryo from trans-placental transfer of proteins produced by the maternal liver. It is possible that the gross immunodeficiency apparent in the *CSF1RKO* predisposes to infection but they are not granulocyte-deficient and the failure to thrive is very consistent and is a great deal more severe than the immunoglobulin knockout in chicken (63). Based upon our previous studies (3) it would likely be possible to rescue the tissue macrophage populations by *in ovo* transfer of wild-type marrow. By contrast to the lethality of the *CSF1RKO* in the chick, homozygous *Csf1r* mutant mice and rats can be relatively long-lived (17, 48, 64, 65). It appears birds lack alternative pathways to mitigate the impact of CSF1R deficiency. Whereas *CSF1RKO* birds were almost completely monocyte deficient (Figure 4), in rodents, CSF1R is not absolutely required for monocytopoiesis (48, 64, 66). Tissue macrophages in some major organs, notably lung, liver and intestine, are only partly CSF1R-dependent (17, 48, 64, 65). Interestingly, in spleen and lymph nodes of *Csf1rko* rats, the two major marginal zone macrophage populations defined by expression of SIGLEC1 (CD169) and CD209B (SIGNR1) are absent (15). Whereas TIMD4^+^/KULO1^+^ macrophages were absent in *CSF1RKO* spleen (Figure 5), they were resistant to depletion with anti-CSF1 treatment (10). This difference likely reflects redundancy of CSF1R ligands, since *IL34* is expressed at similar levels to *CSF1* in the spleen of chicken hatchlings (67).

The rapid decline in viability of *CSF1RKO* chickens from day 6 clearly compromises the utility of the model for functional studies of macrophage biology. Nevertheless, we highlighted a profound loss of B lymphocytes. The phenotype is more severe than the effect of immunoglobulin gene deletions, which deplete mature circulating B cells but retain B cells in the bursa (63, 68). The most obvious explanation is the likely loss of the key B cell survival factor, B cell Activating Factor of the TNF Family (BAFF, *TNFSF13B*), which is highly-expressed in the bursa (69, 70), by follicular macrophages in the spleen (12) and by bone marrow-derived chicken macrophages grown in CSF1 (67). The effect of the *CSF1R* mutation in the bursa was more penetrant than we observed following post hatch anti-CSF1 treatment (10). This may also reflect a partial redundancy*, IL34* is also highly-expressed in bursa (41, 67). We infer that bursal involution in the *CSF1RKO* is due to the complete loss of bursal macrophages as shown in Figure 6. Interestingly, *TNFSF13B* expression is greatly reduced in fetal liver and bone marrow of *Csf1rko* rats (15). CSF1R in mammals is not expressed by B lymphocytes, but in common with the chicken, mutations of either CSF1 or CSF1R lead to selective loss of B lymphocytes (48, 61). In the rat, this deficiency can be rescued by transfer of wild-type marrow at weaning. The donor cells do not generate B cells (48, 61) but instead re-establish the hematopoietic niche for B cell development associated with periosteal location in the bone marrow (71) allowing regeneration of recipient B cells. The bursal M cell-like antigen sampling cells, which express CSF1R, were retained in the *CSF1RKO.* However, we cannot eliminate the possibility that the function of these cells is compromised by the loss of CSF1R and contributes to involution.

Our analysis further highlights unique features of antigen-presenting cell (APC) populations in birds. CSF1R-dependent chicken blood monocytes express high levels of class II MHC. The liver contains a major population of sinusoidal APC, in excess and distinct from actively phagocytic Kupffer cells that express TIMD4 (9). Both of these liver myeloid populations were depleted by anti-CSF1 treatment (10). The CSF1R-dependent TIMD4^+^/KUL01^+^ macrophages of the splenic periellipsoid zone also express Class II MHC and appear to combine the functions of red pulp macrophages in iron homeostasis and marginal zone macrophages in antigen trapping. The UMAP analysis in Figure 2 suggests this population can be separated into two subpopulations. In the spleen, we have extended phenotypic analysis of follicular CSF1R^+^, MAFB^+^, SPIC^+^ macrophages that likely perform the trophic functions of mammalian FDC. These cells were clearly distinct from tingible body macrophages that express TIMD4 and likely contribute to the removal of apoptotic B cells (43). In the bursa we demonstrate that the functionally similar cells referred to as bursal secretory dendritic cells (BSDC) are also a SPIC^+^ macrophage population. On the basis of poor CD45 staining and morphological criteria, it was previously proposed that chicken FDC, like their mammalian namesakes, are not haemopoietic cells (72). In overview, our data demonstrate based upon gene expression and CSF1R-dependence that the cells referred to as bursal secretory dendritic cells (BSDC) and follicular dendritic cells (FDC) (19, 20) would be more appropriately called bursal and splenic follicular macrophages. The expression and function of the SPIC transcription factor has clearly diverged between mammalian and avian lineages. In mammalian spleen, SPIC is associated with the development of macrophages in the red pulp (37) related to their function in iron recycling and erythrophagocytosis. More generally, although B cell selection and affinity maturation are shared features of acquired immunity, it appears that *bona fide* germinal centers (GC) evolved independently in mammals and birds (73), likely reflected in the contrasting cell origins of their FDC as demonstrated herein.

In the adult spleen, cells that share many transcriptional characteristics with the cDC1 subset of mammalian DC are considerably more abundant than their counterparts in mammalian lymphoid tissues (12). Unlike cDC1 in mice, these cells in the chicken also express CSF1R and the *CSF1R* reporter transgenes and they are at least partly CSF1R-dependent. The biology of FLT3 may vary between mammalian species; by contrast to mouse a loss of function mutation in FLT3 ligand leads to monocyte deficiency (74). Conversely, administration of a CSF1-Fc protein to hatchling chickens led to expansion of cells expressing a *CSF1R*-mApple transgene in all tissues examined (3). Interestingly, *CSF1* and *CSF1R* genes both show extensive evidence of positive selection during avian evolution that is less evident in mammals (5). Taken together, our results indicate that the immunomodulatory functions of CSF1/CSF1R are unique to birds and worthy of further study in other bird species.

## Acknowledgments

This work was funded by Biotechnology and Biological Sciences Research Council (BBSRC) through the project grant BB/R003653/1. Roslin Scientists were funded by BBSRC Institute Strategic Programme grants (grant references BBS/E/RL/230001C and BBS/E/RL/230002A). We thank staff at the National Avian Research Facility of the Roslin Institute for care and maintenance of the chickens used in this study (BBSRC Core Capability Grant, BB/CCG2270/1). Ms Hazel Gilhooley provided assistance with genotyping. DAH is supported by NHMRC (Australia) Investigator Grant 2009750 and core support from The Mater Foundation.

## Author contributions

ZW, RH, AI, FF, JM, BS, AB performed experiments and contributed to data analysis. SLM, MM, DAH and AB contributed funding, planning and supervision. DAH wrote the manuscript and SLM, ZW and AB edited the draft.

---

**Figure S1.**
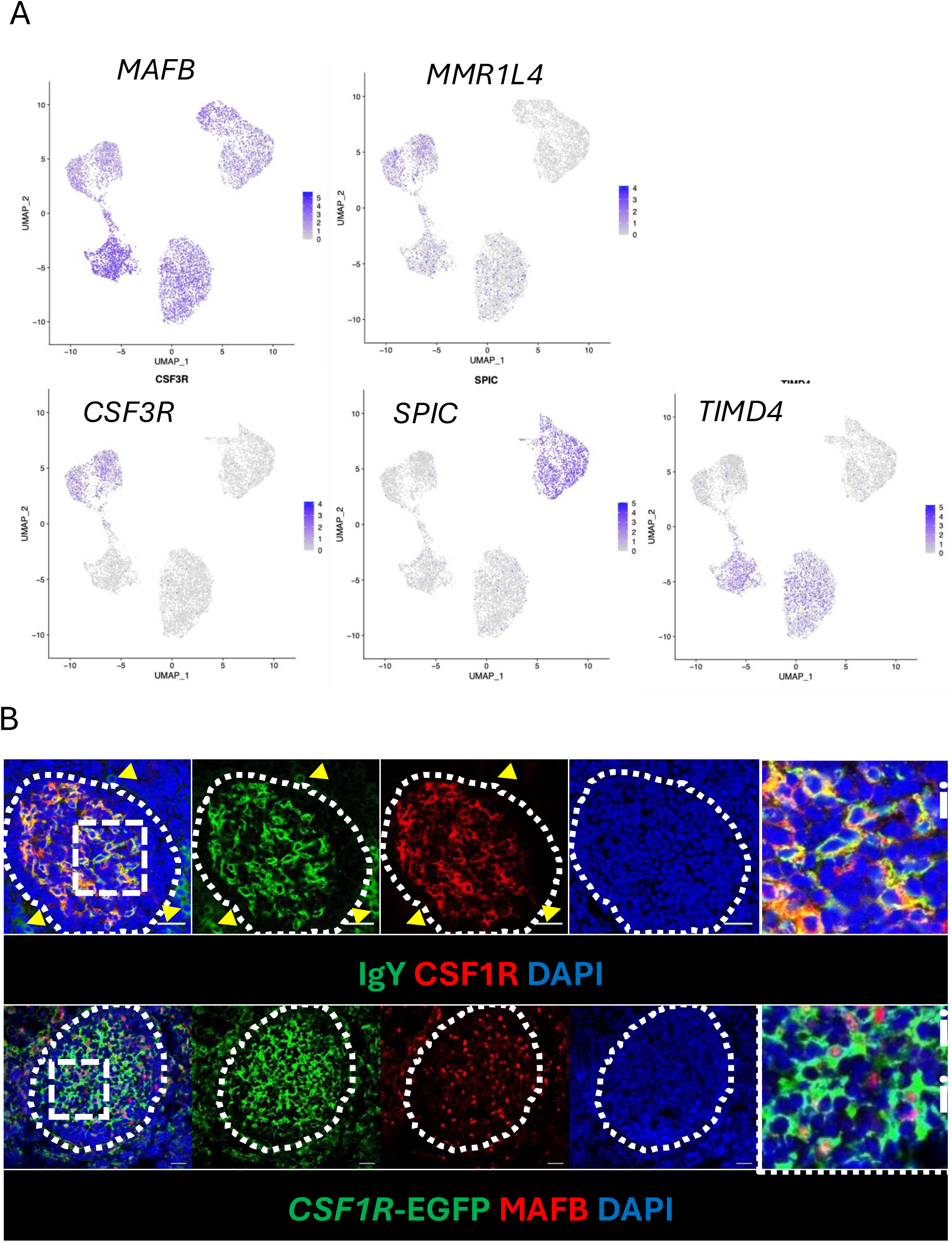
Analysis of chicken splenic macrophage subpopulations. A) Overlay of specific markers on clusters defined by single cell RNA-seq data in Figure 2A. MMR1L4 encodes the marker KUL01. B) Representative images of immunolocalisation of CSF1R protein and surface bound immune complex (identified by IgY staining) in germinal centre. IgY^+^ plasma cells (yellow arrows) lack detectable CSF1R. Detail of dashed box region shown in i. B) Chicken germinal centre FDCs express high levels of the CSF1R-eGFP transgene and show nuclear staining of MAFB. Detail of dashed box region shown in ii. Scale bars = 50 µm

**Figure S2.**
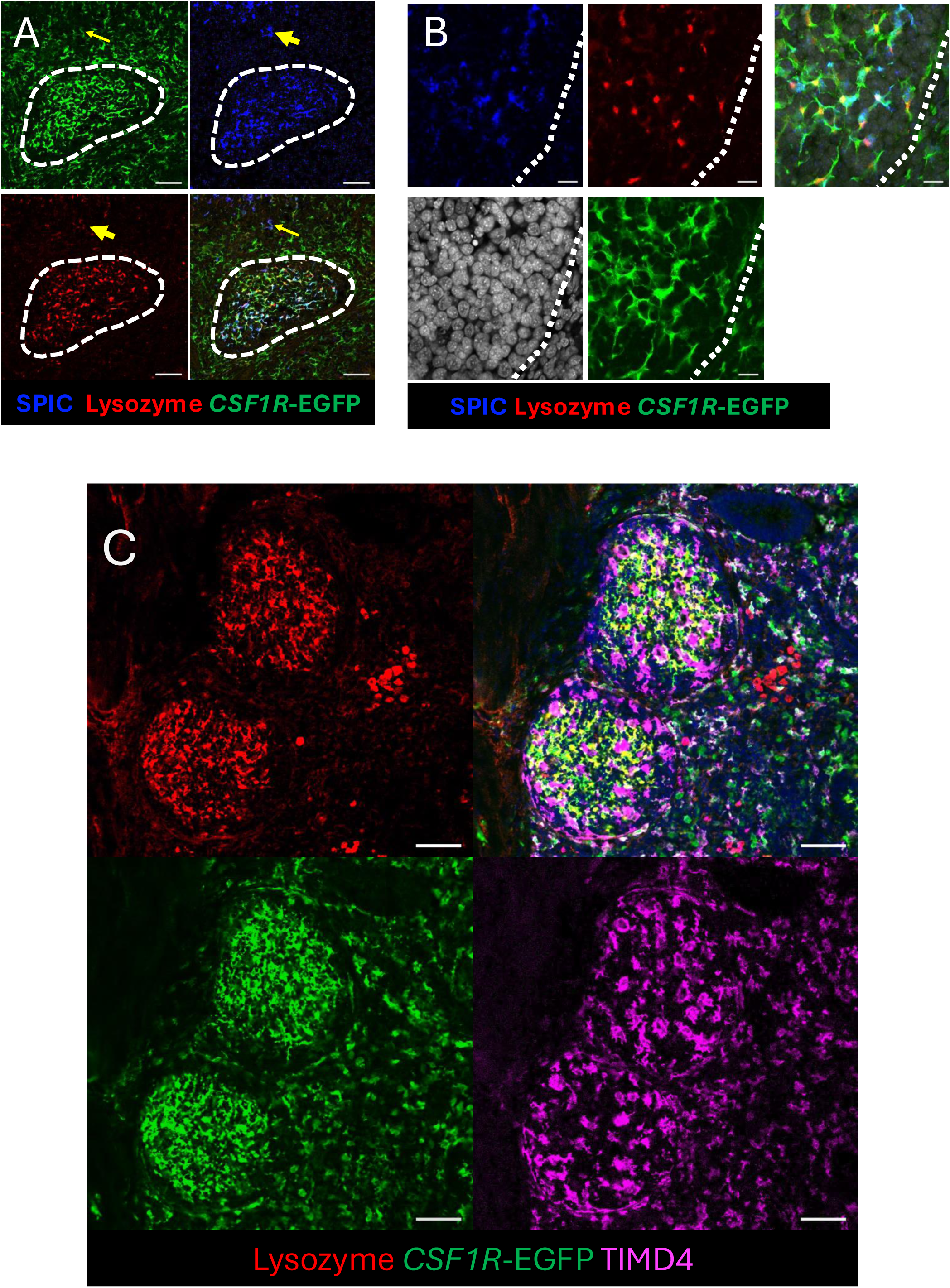
Macrophage subpopulations in the caecal tonsil and spleen. A. Representative images of combined detection of EGFP and lysozyme by immunofluorescence and *SPIC* by RNAScope *in situ* hybridisation within the adult caecal tonsil. Note that macrophages sin the germinal centre (dashed white line) express both SPIC and lysozyme. GFP^+^/SPIC^+^/ lysozyme negative (yellow arrow) are also located in the lamina propria tissue surrounding germinal centres. Scale bars = 50 µm B) High magnification detail of caecal tonsil germinal centre (dotted line). Scale bars = 10 µm. C) Representative image of immunolocalization of EGFP, Lysozyme and TIMD4 in adult spleen.

**Figure S3.**
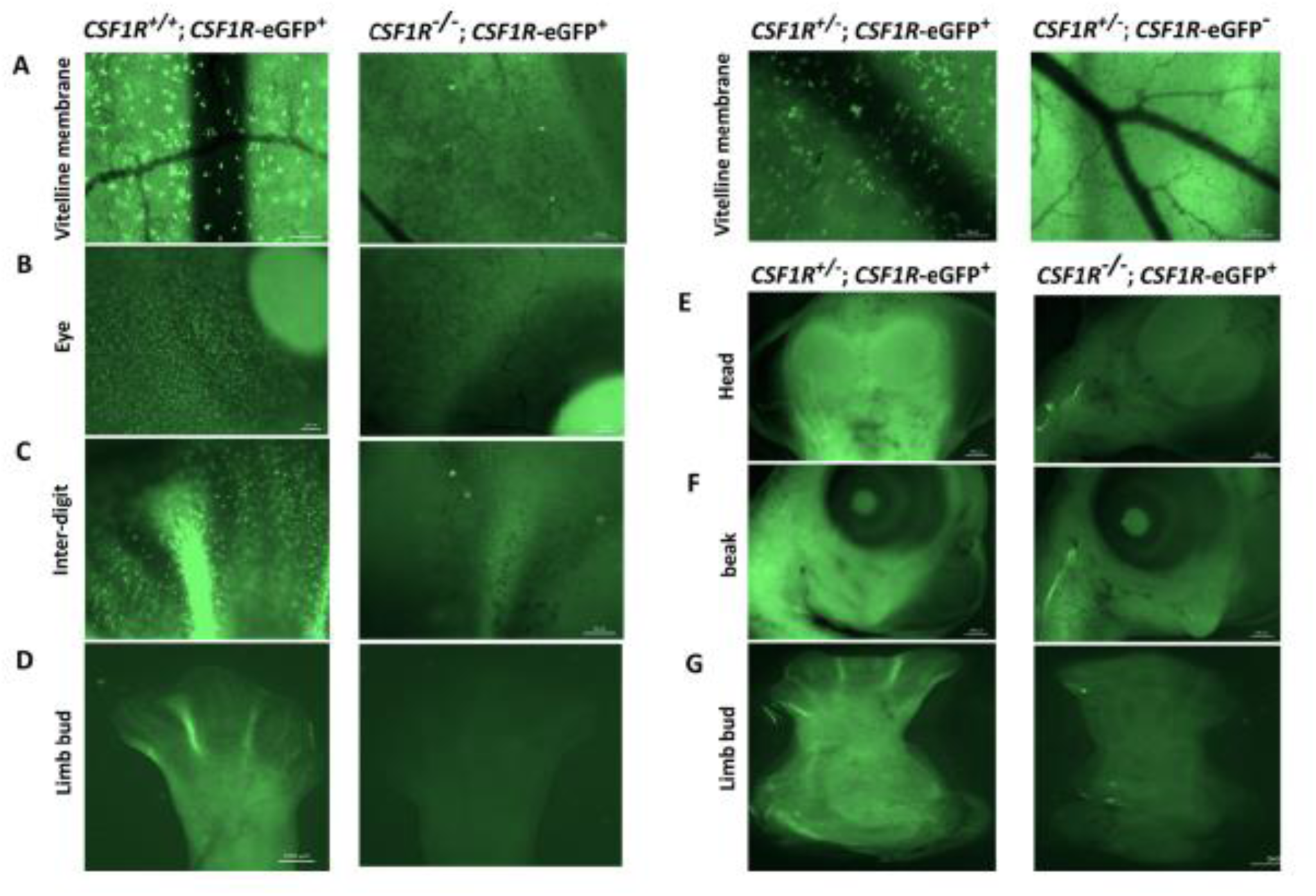
Whole mount imaging of *CSF1R-*EGFP in *CSF1RKO* embryos. A cohort of embryos from *CSF1R^+/-^, CSF1R*-EGFP^+^ matings was imaged as whole mounts at day 8 of incubation and subsequently genotyped for both the mutation and EGFP. CSF1R^-/-^embryos were devoid of detectable EGFP whereas WT and heterozygous embryos were macroscopically fluorescent throughout the body as shown.

**Figure S4.**
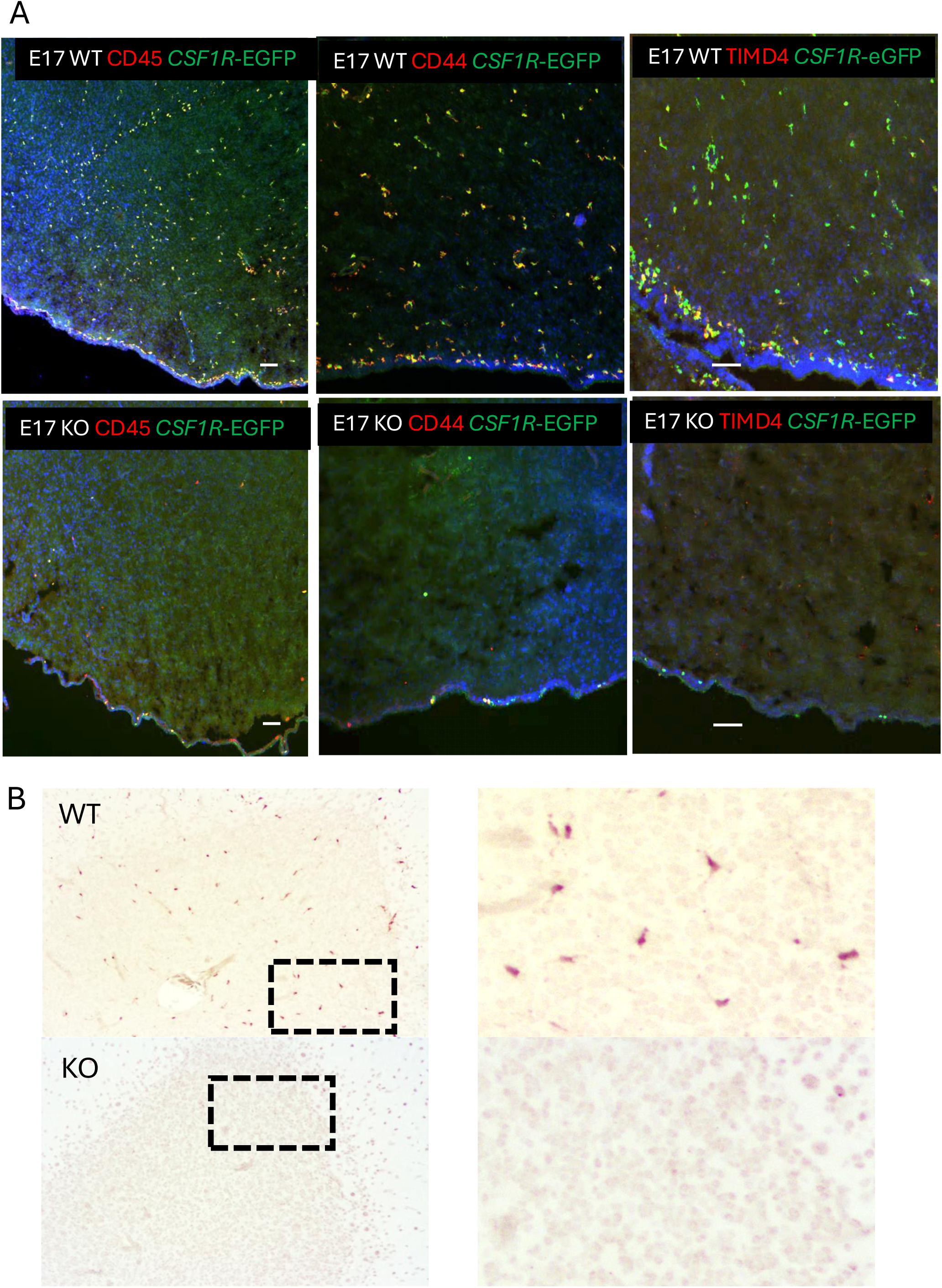
Loss of microglia in *CSF1RKO* chicks. A) Representative images of brain cortex and meninges of E17 WT and *CSF1RKO* embryos stained to detect CD45, CD44, TIMD4 and EGFP as indicated. B) Immunohistochemical localisation of GFP in WT and *CSF1RKO* birds at postnatal day 3. Panel at right shows higher magnification.

