## Supplementary material for "CSF1R-dependent macrophages control B cell development and function in the chicken immune system": Table S1

DE_markers

| | p_val | avg_logFC | pct.1 | pct.2 | p_val_adj | cluster | gene |
|---|---|---|---|---|---|---|---|
| **HSP90AA1** | 0 | -0.871751838363643 | 0.619 | 0.848 | 0 | 0 | HSP90AA1 |
| **CXCL13** | 0 | -1.78458819148731 | 0.126 | 0.528 | 0 | 0 | CXCL13 |
| **ENSGALG00000026970** | 7.26620849865874e-304 | -0.693230567231047 | 0.246 | 0.641 | 9.19974658015184e-300 | 0 | ENSGALG00000026970 |
| **HSPA8** | 5.28653765929335e-298 | -0.705126755600707 | 0.393 | 0.706 | 6.69328533043131e-294 | 0 | HSPA8 |
| **TGM4** | 1.80231724226195e-289 | -1.00079226222568 | 0.056 | 0.438 | 2.28191386042785e-285 | 0 | TGM4 |
| **EXFABP** | 2.89838449495155e-280 | -1.49727564969044 | 0.045 | 0.303 | 3.66964460905816e-276 | 0 | EXFABP |
| **ACTB** | 8.87195276689931e-279 | -0.512284413567881 | 0.964 | 0.986 | 1.12327793981712e-274 | 0 | ACTB |
| **DDX5** | 6.04140424681077e-266 | -0.613331114426264 | 0.241 | 0.619 | 7.64902191688712e-262 | 0 | DDX5 |
| **FOS** | 1.26728879108497e-264 | -0.731238956748866 | 0.121 | 0.491 | 1.60451433839268e-260 | 0 | FOS |
| **TMSB4X** | 1.61913360584327e-256 | -0.520384004243545 | 0.995 | 0.993 | 2.04998505835816e-252 | 0 | TMSB4X |
| **MARCO** | 1.43245862779689e-255 | -0.848431874312103 | 0.172 | 0.455 | 1.81363586865364e-251 | 0 | MARCO |
| **ENSGALG00000011190** | 1.93639024182226e-248 | -0.741997353051794 | 0.16 | 0.477 | 2.45166368517117e-244 | 0 | ENSGALG00000011190 |
| **S100A10** | 2.49985084744799e-248 | -0.805957421852848 | 0.19 | 0.456 | 3.1650611579539e-244 | 0 | S100A10 |
| **RPS11** | 1.97969623602343e-207 | -0.666307984102253 | 0.74 | 0.802 | 2.50649340442926e-203 | 0 | RPS11 |
| **ENSGALG00000004509** | 1.04365481356908e-205 | -0.513873238081065 | 0.794 | 0.892 | 1.32137135945982e-201 | 0 | ENSGALG00000004509 |
| **S100A6** | 5.91095612974476e-205 | -0.854723968485176 | 0.142 | 0.34 | 7.48386155586984e-201 | 0 | S100A6 |
| **JUN** | 1.85016683666587e-190 | -0.623949986566869 | 0.042 | 0.416 | 2.34249623190266e-186 | 0 | JUN |
| **RPL37** | 5.45855649051435e-178 | -0.541568246175426 | 0.609 | 0.748 | 6.91107837264022e-174 | 0 | RPL37 |
| **TPT1** | 1.18489979923132e-177 | -0.570260895807812 | 0.755 | 0.816 | 1.50020163580678e-173 | 0 | TPT1 |
| **BTG2** | 1.64640177600007e-177 | -0.511309696606765 | 0.076 | 0.402 | 2.08450928859369e-173 | 0 | BTG2 |
| **S100A4** | 1.99740367997712e-176 | -0.838376039694724 | 0.038 | 0.258 | 2.52891279921903e-172 | 0 | S100A4 |
| **RPLP2** | 2.09624414690204e-175 | -0.571375085584381 | 0.72 | 0.794 | 2.65405471439268e-171 | 0 | RPLP2 |
| **RPS19** | 5.88120106442289e-172 | -0.554181500629265 | 0.822 | 0.859 | 7.44618866766582e-168 | 0 | RPS19 |
| **RPS14** | 2.53849220279214e-170 | -0.539207129607357 | 0.748 | 0.819 | 3.21398497795513e-166 | 0 | RPS14 |
| **CRIP1** | 8.55277310554663e-170 | -0.640194016312521 | 0.049 | 0.308 | 1.08286660289326e-165 | 0 | CRIP1 |
| **B2M** | 8.70465398152506e-170 | -0.398834660681551 | 0.896 | 0.945 | 1.10209624060089e-165 | 0 | B2M |
| **PPDPF** | 9.22943895035737e-167 | -0.700847202193419 | 0.271 | 0.394 | 1.16853926550475e-162 | 0 | PPDPF |
| **RPL39L** | 9.36290516249876e-167 | -0.45801584387439 | 0.746 | 0.861 | 1.18543742262397e-162 | 0 | RPL39L |
| **RPS17** | 2.03973000456312e-162 | -0.507286324396046 | 0.784 | 0.84 | 2.58250215877736e-158 | 0 | RPS17 |
| **RPS8** | 1.61452087742566e-161 | -0.494743109333021 | 0.754 | 0.846 | 2.04414488290863e-157 | 0 | RPS8 |
| **SH3BGRL3** | 6.09305152327644e-158 | -0.413033147385671 | 0.16 | 0.475 | 7.71441253362031e-154 | 0 | SH3BGRL3 |
| **RPS3A** | 1.14401997840979e-155 | -0.501669581945512 | 0.629 | 0.752 | 1.44844369466464e-151 | 0 | RPS3A |
| **EEF2** | 6.32986648428857e-155 | -0.475374672029934 | 0.367 | 0.578 | 8.01424395575775e-151 | 0 | EEF2 |
| **RPS15** | 3.0909493702551e-154 | -0.514549598042858 | 0.711 | 0.792 | 3.91345099767999e-150 | 0 | RPS15 |
| **ENSGALG00000015183** | 7.84527632420637e-150 | -0.426376067642604 | 0.072 | 0.379 | 9.93290435407769e-146 | 0 | ENSGALG00000015183 |
| **EEF1A1** | 1.39322461345942e-149 | -0.471796999866729 | 0.769 | 0.836 | 1.76396168310097e-145 | 0 | EEF1A1 |
| **RPL27A** | 2.63424017510062e-149 | -0.517761051920075 | 0.584 | 0.701 | 3.3352114856949e-145 | 0 | RPL27A |
| **RPL37A** | 7.85646338667898e-149 | -0.48881926436345 | 0.681 | 0.774 | 9.94706829387425e-145 | 0 | RPL37A |
| **RPL24** | 5.56712389505135e-148 | -0.48486191792371 | 0.601 | 0.714 | 7.04853556352451e-144 | 0 | RPL24 |
| **RPS27A** | 6.72732635338863e-148 | -0.499027923120234 | 0.629 | 0.73 | 8.51746789602534e-144 | 0 | RPS27A |
| **LY86** | 2.51747362757995e-145 | -0.506221294346005 | 0.077 | 0.304 | 3.18737335987898e-141 | 0 | LY86 |
| **ITGB1BP3** | 1.57756615790286e-144 | -0.440116601967775 | 0.37 | 0.594 | 1.99735651252081e-140 | 0 | ITGB1BP3 |
| **TSC22D4** | 1.83902434167163e-144 | -0.420060319732392 | 0.076 | 0.356 | 2.32838871899046e-140 | 0 | TSC22D4 |
| **ANKRD11** | 1.20886921979414e-143 | -0.398542650393917 | 0.117 | 0.408 | 1.53054931918137e-139 | 0 | ANKRD11 |
| **RPL8** | 1.27224132534076e-141 | -0.468275693708631 | 0.519 | 0.675 | 1.61078474201393e-137 | 0 | RPL8 |
| **RPS23** | 8.88718757582433e-141 | -0.474850049681539 | 0.628 | 0.728 | 1.12520681897512e-136 | 0 | RPS23 |
| **RPL26L1** | 1.60693591621993e-140 | -0.480336660319147 | 0.54 | 0.676 | 2.03454156352605e-136 | 0 | RPL26L1 |
| **EEF1D** | 2.2373904794096e-140 | -0.479353543263281 | 0.329 | 0.508 | 2.83276008598049e-136 | 0 | EEF1D |
| **RPL17** | 2.40502911067671e-140 | -0.456387143466459 | 0.673 | 0.778 | 3.04500735702778e-136 | 0 | RPL17 |
| **RPS15A** | 1.3136078183443e-139 | -0.470868781658722 | 0.725 | 0.8 | 1.66315885880572e-135 | 0 | RPS15A |
| **RPL35A** | 4.47967509739469e-139 | -0.470658585038894 | 0.657 | 0.766 | 5.67171664081142e-135 | 0 | RPL35A |
| **RPS16** | 1.63171485423154e-138 | -0.463771347025172 | 0.57 | 0.692 | 2.06591417694256e-134 | 0 | RPS16 |

| | | | | | | | |
|---|---|---|---|---|---|---|---|
| **DDX3X** | 1.88420940334202e-138 | -0.366692038630225 | 0.123 | 0.418 | 2.38559752557133e-134 | 0 | DDX3X |
| **ENSGALG00000037441** | 2.11534187158552e-138 | -0.490699377618552 | 0.738 | 0.793 | 2.67823434361442e-134 | 0 | ENSGALG00000037441 |
| **RPL13** | 7.72871515443552e-138 | -0.455495546030628 | 0.514 | 0.671 | 9.78532625703082e-134 | 0 | RPL13 |
| **RPL34** | 8.1656855147995e-138 | -0.476615013036923 | 0.671 | 0.745 | 1.03385744302876e-133 | 0 | RPL34 |
| **RPL15** | 2.21924035123782e-137 | -0.462418686858187 | 0.704 | 0.796 | 2.8097802087022e-133 | 0 | RPL15 |
| **RPS4Y1** | 4.49824363961125e-136 | -0.438356999384183 | 0.386 | 0.584 | 5.69522627211181e-132 | 0 | RPS4Y1 |
| **CSF1R** | 6.34854926199129e-136 | -0.368608409591764 | 0.206 | 0.487 | 8.03789822060717e-132 | 0 | CSF1R |
| **GNB2L1** | 8.20312807811645e-136 | -0.476949788798282 | 0.543 | 0.661 | 1.03859804597032e-131 | 0 | GNB2L1 |
| **MGST1** | 4.98664633868148e-135 | -0.459451599210763 | 0.048 | 0.295 | 6.31359292940462e-131 | 0 | MGST1 |
| **LDHA** | 1.02111758019152e-133 | -0.479062295686073 | 0.042 | 0.28 | 1.29283696828048e-129 | 0 | LDHA |
| **RPSAP58** | 6.81032828376302e-133 | -0.403728822480631 | 0.432 | 0.638 | 8.62255664007236e-129 | 0 | RPSAP58 |
| **RPS6** | 7.40755072665094e-133 | -0.435675697730201 | 0.512 | 0.666 | 9.37869997501276e-129 | 0 | RPS6 |
| **ENSGALG00000030007** | 1.15890642741409e-132 | -0.493241113851102 | 0.032 | 0.285 | 1.46729142774898e-128 | 0 | ENSGALG00000030007 |
| **VIM** | 4.17561617673876e-132 | -0.527514280891364 | 0.029 | 0.279 | 5.28674764136894e-128 | 0 | VIM |
| **RPL23** | 8.31779803696596e-129 | -0.455236859799121 | 0.764 | 0.809 | 1.05311640946026e-124 | 0 | RPL23 |
| **RPL19** | 1.94950113518013e-128 | -0.440939176183599 | 0.555 | 0.682 | 2.46826338725156e-124 | 0 | RPL19 |
| **RPL27** | 1.17268840561923e-127 | -0.413956574412501 | 0.393 | 0.587 | 1.48474079035451e-123 | 0 | RPL27 |
| **RPS24** | 2.96084959113989e-126 | -0.43771439441727 | 0.605 | 0.709 | 3.74873166734221e-122 | 0 | RPS24 |
| **RPLP1** | 1.06193599589632e-125 | -0.444778150201922 | 0.904 | 0.906 | 1.34451716440433e-121 | 0 | RPLP1 |
| **RPL35** | 1.94338528998279e-125 | -0.416279784793483 | 0.501 | 0.662 | 2.46052011564721e-121 | 0 | RPL35 |
| **HSPA5** | 3.11537111418758e-125 | -0.461912747763067 | 0.25 | 0.455 | 3.94437136767289e-121 | 0 | HSPA5 |
| **RPL21** | 6.47939708037776e-125 | -0.434716918809276 | 0.78 | 0.846 | 8.20356464346629e-121 | 0 | RPL21 |
| **RPS20** | 1.5495676979991e-124 | -0.424599442002211 | 0.691 | 0.773 | 1.96190766243666e-120 | 0 | RPS20 |
| **RPS10** | 1.06071987139953e-123 | -0.446910846539812 | 0.566 | 0.67 | 1.34297742917894e-119 | 0 | RPS10 |
| **ENSGALG00000045199** | 4.15138272260494e-123 | -0.383644431125933 | 0.037 | 0.345 | 5.25606566509012e-119 | 0 | ENSGALG00000045199 |
| **RPL5** | 1.31108456785333e-122 | -0.389968663832401 | 0.334 | 0.541 | 1.6599641713591e-118 | 0 | RPL5 |
| **RPL7A** | 1.30809069653707e-121 | -0.430772978706823 | 0.482 | 0.627 | 1.65617363088558e-117 | 0 | RPL7A |
| **H3F3B** | 3.02393279076177e-121 | -0.369716269594735 | 0.469 | 0.67 | 3.82860130638348e-117 | 0 | H3F3B |
| **RPL11** | 5.65388940980137e-118 | -0.404114261766775 | 0.554 | 0.691 | 7.15838938174952e-114 | 0 | RPL11 |
| **TMSB15B** | 1.48909460398738e-117 | 0.442995920384414 | 0.927 | 0.632 | 1.88534267810842e-113 | 0 | TMSB15B |
| **HTRA1** | 4.97198553561773e-117 | -0.371922801299107 | 0.053 | 0.288 | 6.29503088664561e-113 | 0 | HTRA1 |
| **RPS7** | 1.54533631742911e-116 | -0.396877193016859 | 0.438 | 0.602 | 1.956550311497e-112 | 0 | RPS7 |
| **RPS25** | 1.54826997348414e-116 | -0.409566540147735 | 0.528 | 0.671 | 1.96026461342827e-112 | 0 | RPS25 |
| **ANXA1** | 2.27453822896954e-116 | -0.77058817632952 | 0.012 | 0.272 | 2.87979285169833e-112 | 0 | ANXA1 |
| **ATP6V0C** | 6.10059841087823e-115 | -0.404421719763741 | 0.448 | 0.594 | 7.72396764801293e-111 | 0 | ATP6V0C |
| **FAM49A** | 1.38313012013455e-113 | -0.327513693917368 | 0.221 | 0.472 | 1.75118104510235e-109 | 0 | FAM49A |
| **DNAJA1** | 1.90194162736874e-113 | -0.321117439899898 | 0.127 | 0.379 | 2.40804829441156e-109 | 0 | DNAJA1 |
| **ZEB2** | 6.8091432344294e-113 | -0.353351941347294 | 0.149 | 0.367 | 8.62105624911107e-109 | 0 | ZEB2 |
| **RPL18A** | 2.94401905093851e-111 | -0.404600508020391 | 0.463 | 0.603 | 3.72742252039325e-107 | 0 | RPL18A |
| **FN1** | 6.77320674814402e-111 | -0.872004594869402 | 0.009 | 0.276 | 8.57555706382514e-107 | 0 | FN1 |
| **RPL7** | 1.43160394176722e-110 | -0.404828931754093 | 0.498 | 0.622 | 1.81255375067148e-106 | 0 | RPL7 |
| **CD81** | 3.03942156205984e-110 | -0.328508401841417 | 0.092 | 0.322 | 3.84821163972397e-106 | 0 | CD81 |
| **RPL32** | 3.54588057542853e-110 | -0.399237549735968 | 0.573 | 0.701 | 4.48943939655006e-106 | 0 | RPL32 |
| **RPL36** | 3.7288780540599e-109 | -0.393887883049605 | 0.423 | 0.576 | 4.72113250424524e-105 | 0 | RPL36 |
| **RPL23A** | 4.01360032224992e-109 | -0.405659683327138 | 0.533 | 0.641 | 5.08161936800062e-105 | 0 | RPL23A |
| **NFKBIA** | 4.16795977945754e-109 | -0.351105074720557 | 0.18 | 0.392 | 5.27705387677119e-105 | 0 | NFKBIA |
| **RPL9** | 4.17271439965195e-108 | -0.381004389390398 | 0.459 | 0.622 | 5.28307370139933e-104 | 0 | RPL9 |
| **RPL36AL** | 8.48970706372492e-107 | -0.385740051522017 | 0.53 | 0.658 | 1.07488181133821e-102 | 0 | RPL36AL |
| **RPS12** | 4.52737998538994e-106 | -0.401471465730058 | 0.435 | 0.571 | 5.73211579950221e-102 | 0 | RPS12 |
| **PTPRC** | 1.81404322676652e-105 | -0.313452500213681 | 0.193 | 0.446 | 2.29676012940909e-101 | 0 | PTPRC |
| **RPS3** | 4.10480228103186e-105 | -0.377196768458057 | 0.402 | 0.56 | 5.19709016801444e-101 | 0 | RPS3 |
| **RPL31** | 5.25002315662257e-105 | -0.385881385724798 | 0.52 | 0.642 | 6.64705431859984e-101 | 0 | RPL31 |
| **CD47** | 1.43836815840063e-104 | -0.320042012544912 | 0.194 | 0.421 | 1.82111792535104e-100 | 0 | CD47 |
| **RPS28** | 1.91062589373754e-104 | -0.393980684012895 | 0.523 | 0.633 | 2.4190434440611e-100 | 0 | RPS28 |

| | | | | | | | |
|---|---|---|---|---|---|---|---|
| **DSTN** | 8.21580792437179e-104 | -0.334772699754944 | 0.519 | 0.694 | 1.04020344130471e-99 | 0 | DSTN |
| **ACTR3** | 8.32272014593778e-101 | -0.294301953602521 | 0.205 | 0.45 | 1.05373959767718e-96 | 0 | ACTR3 |
| **MT-CO1** | 8.33219408893756e-101 | -0.372344962934479 | 0.972 | 0.978 | 1.05493909360038e-96 | 0 | MT-CO1 |
| **SPIC** | 1.46045364497817e-100 | -0.374559192601918 | 0.02 | 0.271 | 1.84908035990686e-96 | 0 | SPIC |
| **FYB** | 3.53945087643102e-100 | -0.290897384035811 | 0.14 | 0.376 | 4.48129875464931e-96 | 0 | FYB |
| **ITM2A** | 1.80329614896243e-99 | -0.372780142847852 | 0.446 | 0.575 | 2.28315325420133e-95 | 0 | ITM2A |
| **DUSP1** | 3.42204819082293e-99 | -0.352049549691741 | 0.028 | 0.255 | 4.33265521440091e-95 | 0 | DUSP1 |
| **PTPN6** | 6.05889492793263e-99 | -0.279298962993799 | 0.16 | 0.402 | 7.6711668682555e-95 | 0 | PTPN6 |
| **RPL30** | 1.11871388605921e-98 | -0.377316170396734 | 0.486 | 0.609 | 1.41640365113956e-94 | 0 | RPL30 |
| **RPL22** | 3.45025444481369e-98 | -0.361904803314498 | 0.501 | 0.631 | 4.36836715257861e-94 | 0 | RPL22 |
| **RPS26** | 7.99481101986235e-98 | -0.40697017473927 | 0.684 | 0.72 | 1.01222302322477e-93 | 0 | RPS26 |
| **RPL4** | 2.31274926189125e-97 | -0.341406214781806 | 0.249 | 0.428 | 2.92817184048051e-93 | 0 | RPL4 |
| **SMAP2** | 3.93291707375294e-97 | -0.285415644325629 | 0.189 | 0.423 | 4.9794663070786e-93 | 0 | SMAP2 |
| **RPL29** | 3.79642432337767e-95 | -0.353395778683098 | 0.527 | 0.661 | 4.80665283582846e-91 | 0 | RPL29 |
| **UBA52** | 1.5084329823678e-94 | -0.355972012774478 | 0.482 | 0.604 | 1.90982699897587e-90 | 0 | UBA52 |
| **CCNI** | 2.53291551085601e-93 | -0.265470104408477 | 0.1 | 0.325 | 3.2069243282948e-89 | 0 | CCNI |
| **ENSGALG00000028749** | 1.0100169341603e-92 | -0.3122539683051 | 0.542 | 0.706 | 1.27878244034036e-88 | 0 | ENSGALG00000028749 |
| **MYH9** | 1.99429931468265e-92 | -0.302697001998153 | 0.289 | 0.511 | 2.5249823623197e-88 | 0 | MYH9 |
| **RPS21** | 2.3619923840248e-92 | -0.358577375003234 | 0.432 | 0.57 | 2.9905185574138e-88 | 0 | RPS21 |
| **ENO1** | 9.96362526383749e-92 | -0.289865178829342 | 0.261 | 0.462 | 1.26149459465446e-87 | 0 | ENO1 |
| **CORO1C** | 8.97287516676714e-91 | -0.305745677904567 | 0.309 | 0.506 | 1.13605572486439e-86 | 0 | CORO1C |
| **RAB1B** | 1.51938696617033e-90 | -0.268336983370338 | 0.171 | 0.393 | 1.92369583786825e-86 | 0 | RAB1B |
| **RPS13** | 1.77619424300938e-89 | -0.343686055976924 | 0.525 | 0.639 | 2.24883953107418e-85 | 0 | RPS13 |
| **ENSGALG00000052354** | 9.56548938669156e-89 | -0.254500287058875 | 0.078 | 0.292 | 1.21108661124902e-84 | 0 | ENSGALG00000052354 |
| **GAPDH** | 4.35274820723064e-88 | -0.349837098334344 | 0.429 | 0.559 | 5.51101450517471e-84 | 0 | GAPDH |
| **GNG5** | 1.28783423837144e-87 | -0.291176273952465 | 0.348 | 0.544 | 1.63052692920208e-83 | 0 | GNG5 |
| **RPL6** | 1.10410736602694e-86 | -0.327174391570393 | 0.461 | 0.602 | 1.39791033612671e-82 | 0 | RPL6 |
| **ARPC4** | 1.15473376040643e-86 | -0.282628558139871 | 0.352 | 0.55 | 1.46200841405057e-82 | 0 | ARPC4 |
| **CST3** | 4.25586989687856e-86 | -0.322323969297747 | 0.625 | 0.742 | 5.38835687643795e-82 | 0 | CST3 |
| **RPL12** | 4.53885011501933e-86 | -0.353374992181258 | 0.526 | 0.626 | 5.74663813062598e-82 | 0 | RPL12 |
| **GNAI2** | 2.77651740683953e-85 | -0.258356483294397 | 0.13 | 0.332 | 3.51534868879953e-81 | 0 | GNAI2 |
| **RPL14** | 4.11788714500232e-85 | -0.323659040157663 | 0.465 | 0.593 | 5.21365691428743e-81 | 0 | RPL14 |
| **MCL1** | 5.63690174883925e-85 | -0.250935380577476 | 0.092 | 0.294 | 7.13688130420538e-81 | 0 | MCL1 |
| **RPS29** | 2.99210533482056e-84 | -0.332823606267752 | 0.429 | 0.555 | 3.78830456441631e-80 | 0 | RPS29 |
| **CALM2** | 1.54418078231491e-83 | -0.285314673618967 | 0.282 | 0.474 | 1.95508728848891e-79 | 0 | CALM2 |
| **RPL10A** | 1.82904000941061e-83 | -0.32487121276248 | 0.454 | 0.581 | 2.31574755591478e-79 | 0 | RPL10A |
| **FAU** | 1.93019382825982e-83 | -0.331065413475367 | 0.431 | 0.548 | 2.44381840595976e-79 | 0 | FAU |
| **YBX1** | 5.79332280432308e-83 | -0.277774428679604 | 0.36 | 0.573 | 7.33492600255345e-79 | 0 | YBX1 |
| **SLC40A1** | 1.1872333389878e-82 | 0.349857873862355 | 0.718 | 0.317 | 1.50315613049245e-78 | 0 | SLC40A1 |
| **ENSGALG00000047532** | 2.21861200809296e-82 | -0.263776452583058 | 0.213 | 0.415 | 2.80898466344649e-78 | 0 | ENSGALG00000047532 |
| **COX3** | 5.10919549447923e-82 | -0.31445659074961 | 0.987 | 0.99 | 6.46875241556015e-78 | 0 | COX3 |
| **EEF1B2** | 1.27888800028433e-81 | -0.300350494421623 | 0.505 | 0.642 | 1.61920009715998e-77 | 0 | EEF1B2 |
| **LCP1** | 5.06243883276154e-81 | -0.278901728696713 | 0.407 | 0.597 | 6.40955380615939e-77 | 0 | LCP1 |
| **PABPC1** | 7.38174890828599e-81 | -0.299792868651414 | 0.48 | 0.625 | 9.3460322927809e-77 | 0 | PABPC1 |
| **PNRC1** | 7.7023258870652e-80 | -0.326208556688944 | 0.457 | 0.574 | 9.75191480561325e-76 | 0 | PNRC1 |
| **RASSF2** | 2.02285973959375e-79 | -0.260400546187937 | 0.182 | 0.376 | 2.56114271629965e-75 | 0 | RASSF2 |
| **NME2** | 1.6849322197787e-78 | -0.302029129282866 | 0.371 | 0.522 | 2.13329268346182e-74 | 0 | NME2 |
| **MYO9B** | 4.29867890146702e-78 | -0.252741712626676 | 0.196 | 0.415 | 5.44255735714739e-74 | 0 | MYO9B |
| **LAPTM5** | 1.04438208405012e-76 | -0.262217448472722 | 0.343 | 0.54 | 1.32229215661585e-72 | 0 | LAPTM5 |
| **RPS2** | 8.1052223349771e-72 | -0.300075411288928 | 0.546 | 0.642 | 1.02620219983145e-67 | 0 | RPS2 |
| **RPL3** | 8.95388834504211e-72 | -0.289993138619696 | 0.412 | 0.537 | 1.13365180336578e-67 | 0 | RPL3 |
| **BTG1** | 1.6497449480703e-70 | -0.260169130050525 | 0.203 | 0.368 | 2.08874207875181e-66 | 0 | BTG1 |
| **ATP6** | 6.64597907700394e-70 | -0.303574990109418 | 0.961 | 0.974 | 8.41447410939469e-66 | 0 | ATP6 |
| **ENSGALG00000050515** | 4.49603480991018e-62 | -0.305509110124083 | 0.15 | 0.271 | 5.69242967282728e-58 | 0 | ENSGALG00000050515 |

| | | | | | | | |
|---|---|---|---|---|---|---|---|
| **MAFB** | 1.68088995688174e-60 | -0.312179108610124 | 0.577 | 0.634 | 2.12817477440797e-56 | 0 | MAFB |
| **RPL38** | 1.93442550548844e-60 | -0.258563595988916 | 0.454 | 0.567 | 2.44917613249892e-56 | 0 | RPL38 |
| **HSP90B1** | 2.26057365516179e-54 | -0.255964592100829 | 0.338 | 0.476 | 2.86211230480034e-50 | 0 | HSP90B1 |
| **SAT1** | 1.15443418897722e-40 | -0.264184591954805 | 0.573 | 0.55 | 1.46162912666406e-36 | 0 | SAT1 |
| **MAFB.1** | 0 | 1.0370668939763 | 0.948 | 0.555 | 0 | 1 | MAFB |
| **SAT1.1** | 0 | 1.00673044821409 | 0.936 | 0.483 | 0 | 1 | SAT1 |
| **HSP90AA1.1** | 0 | 0.927756363922119 | 0.979 | 0.75 | 0 | 1 | HSP90AA1 |
| **CD83** | 0 | 0.557747096498081 | 0.542 | 0.121 | 0 | 1 | CD83 |
| **PDCD1LG2** | 0 | 0.525950469884227 | 0.549 | 0.127 | 0 | 1 | PDCD1LG2 |
| **CXCL13.1** | 6.84946685697951e-255 | -1.53564441253172 | 0.22 | 0.461 | 8.67210998762176e-251 | 1 | CXCL13 |
| **MERTK** | 4.43934355772587e-254 | 0.41701983781722 | 0.481 | 0.125 | 5.62065287843673e-250 | 1 | MERTK |
| **C1QC** | 1.08426614414345e-251 | 0.70030443915323 | 0.883 | 0.482 | 1.37278936510002e-247 | 1 | C1QC |
| **RSFR** | 6.91191919903135e-240 | 0.525832726084952 | 0.62 | 0.239 | 8.7511808978936e-236 | 1 | RSFR |
| **PNRC1.1** | 1.15311403443656e-237 | 0.611055411033334 | 0.859 | 0.482 | 1.45995767900013e-233 | 1 | PNRC1 |
| **HAVCR1** | 1.31557657160562e-236 | 0.414016668045776 | 0.541 | 0.175 | 1.66565149730987e-232 | 1 | HAVCR1 |
| **HSPA8.1** | 2.29620286569961e-232 | 0.706460323451103 | 0.913 | 0.567 | 2.90722244826228e-228 | 1 | HSPA8 |
| **MYLIP** | 8.93936025935869e-232 | 0.384920076373224 | 0.403 | 0.093 | 1.1318124024374e-227 | 1 | MYLIP |
| **BIRC3** | 2.65668162815109e-227 | 0.393223839785715 | 0.406 | 0.087 | 3.36362460940209e-223 | 1 | BIRC3 |
| **C1QA** | 2.53592732104652e-225 | 0.53107486103739 | 0.765 | 0.409 | 3.210737581177e-221 | 1 | C1QA |
| **ARGLU1** | 4.96166896902628e-225 | 0.436901271311151 | 0.552 | 0.178 | 6.28196908168417e-221 | 1 | ARGLU1 |
| **ENSGALG00000014585** | 2.75288385334963e-222 | 0.672214334912321 | 0.737 | 0.314 | 3.48542624672597e-218 | 1 | ENSGALG00000014585 |
| **ENSGALG00000053121** | 3.27568482672457e-211 | 0.480756133691274 | 0.353 | 0.058 | 4.14734455911597e-207 | 1 | ENSGALG00000053121 |
| **C1QB** | 5.73475574847269e-207 | 0.6053252689318 | 0.78 | 0.37 | 7.26077425314127e-203 | 1 | C1QB |
| **ENSGALG00000004509.1** | 3.27044335871247e-192 | 0.548801672177015 | 0.964 | 0.847 | 4.14070833646585e-188 | 1 | ENSGALG00000004509 |
| **CD47.1** | 2.55370844622569e-191 | 0.469537925927841 | 0.685 | 0.298 | 3.23325026376634e-187 | 1 | CD47 |
| **CDH2** | 2.32427290795394e-189 | 0.339464673718207 | 0.392 | 0.096 | 2.94276192876049e-185 | 1 | CDH2 |
| **NFKBIA.1** | 2.73976755030959e-186 | 0.531133447979991 | 0.682 | 0.269 | 3.46881969544697e-182 | 1 | NFKBIA |
| **SASH1** | 5.33843760375758e-185 | 0.268225745714104 | 0.308 | 0.046 | 6.75899585011748e-181 | 1 | SASH1 |
| **CTSS** | 8.50381775333361e-184 | 0.577158639214621 | 0.918 | 0.605 | 1.07666836853499e-179 | 1 | CTSS |
| **DNAJA1.1** | 7.21594753933589e-178 | 0.431646603056787 | 0.584 | 0.26 | 9.13611117955316e-174 | 1 | DNAJA1 |
| **CYTH1** | 6.87846749987138e-160 | 0.318229931121992 | 0.437 | 0.149 | 8.70882770158716e-156 | 1 | CYTH1 |
| **TGM4.1** | 7.56358501993981e-157 | -0.847399520901478 | 0.097 | 0.383 | 9.5762549937458e-153 | 1 | TGM4 |
| **LITAF** | 4.32357990238894e-148 | 0.322089343686775 | 0.464 | 0.162 | 5.47408451441463e-144 | 1 | LITAF |
| **PLAU** | 5.03746468186788e-148 | 0.26600880499055 | 0.285 | 0.061 | 6.37793403371293e-144 | 1 | PLAU |
| **DUSP7** | 1.99047342186428e-147 | 0.305016529378184 | 0.406 | 0.123 | 2.52013839942237e-143 | 1 | DUSP7 |
| **SCPEP1** | 6.57298196329167e-147 | 0.391349215756663 | 0.571 | 0.245 | 8.32205246372358e-143 | 1 | SCPEP1 |
| **IFI30** | 4.83253255261056e-143 | 0.604045723331348 | 0.997 | 0.792 | 6.11846946486023e-139 | 1 | IFI30 |
| **INSIG1** | 1.82735319762519e-142 | 0.405067225755823 | 0.44 | 0.157 | 2.31361188351325e-138 | 1 | INSIG1 |
| **BLB1** | 9.48865538871445e-141 | 0.524376212343572 | 0.972 | 0.839 | 1.20135865876514e-136 | 1 | BLB1 |
| **PNISR** | 6.14170616913583e-137 | 0.313216213298143 | 0.436 | 0.14 | 7.77601418074287e-133 | 1 | PNISR |
| **DNASE2B** | 5.65975802701795e-135 | 0.294254307155422 | 0.398 | 0.149 | 7.16581963800742e-131 | 1 | DNASE2B |
| **DCLK3** | 1.95315492407969e-134 | 0.35381798651786 | 0.542 | 0.248 | 2.4728894493773e-130 | 1 | DCLK3 |
| **PMAIP1** | 2.61172815241586e-132 | 0.304555042184809 | 0.27 | 0.052 | 3.30670901377372e-128 | 1 | PMAIP1 |
| **RGS1** | 7.1040665453901e-132 | 0.300387519989955 | 0.335 | 0.082 | 8.99445865311841e-128 | 1 | RGS1 |
| **SDC4** | 1.44414655824891e-128 | 0.250179377239466 | 0.295 | 0.076 | 1.82843395739894e-124 | 1 | SDC4 |
| **STAB1** | 5.72802862456459e-126 | 0.298583440608565 | 0.426 | 0.144 | 7.25225704156122e-122 | 1 | STAB1 |
| **ITSN1** | 5.17546450118824e-124 | 0.301024083904696 | 0.44 | 0.168 | 6.55265560495443e-120 | 1 | ITSN1 |
| **ENSGALG00000011190.1** | 3.20292389667731e-119 | -0.62836940191837 | 0.193 | 0.432 | 4.05522194558314e-115 | 1 | ENSGALG00000011190 |
| **TMEM123** | 3.08558286665024e-117 | 0.310504732200613 | 0.445 | 0.173 | 3.90665646746586e-113 | 1 | TMEM123 |
| **SLC40A1.1** | 1.24785244386669e-114 | 0.490584747795936 | 0.719 | 0.366 | 1.57990597917961e-110 | 1 | SLC40A1 |
| **ATP2B1** | 1.25308683122388e-114 | 0.274438177299714 | 0.383 | 0.136 | 1.58653323701256e-110 | 1 | ATP2B1 |
| **DDX3X.1** | 1.5256297319851e-111 | 0.352635630503787 | 0.6 | 0.289 | 1.93159980366634e-107 | 1 | DDX3X |
| **ZEB2.1** | 9.3379753336065e-104 | 0.412494186135766 | 0.623 | 0.249 | 1.18228105698792e-99 | 1 | ZEB2 |
| **KIAA0232** | 5.00655314227779e-103 | 0.253950098702385 | 0.333 | 0.113 | 6.33879693343791e-99 | 1 | KIAA0232 |

| | | | | | | | |
|---|---|---|---|---|---|---|---|
| **DDX5.1** | 4.98487732876471e-102 | 0.43067840810343 | 0.818 | 0.461 | 6.31135318594899e-98 | 1 | DDX5 |
| **SGK1** | 1.24028294708692e-99 | 0.31196155462913 | 0.415 | 0.134 | 1.57032223930675e-95 | 1 | SGK1 |
| **ENSGALG00000045115** | 3.62388622793867e-98 | 0.345617079192277 | 0.483 | 0.226 | 4.58820235319315e-94 | 1 | ENSGALG00000045115 |
| **DNAJA4** | 5.05052820484133e-98 | 0.25015564738644 | 0.254 | 0.066 | 6.39447376014961e-94 | 1 | DNAJA4 |
| **TIMD4** | 5.3363595671508e-98 | 0.297421319011022 | 0.404 | 0.144 | 6.75636484796963e-94 | 1 | TIMD4 |
| **MARCO.1** | 1.48760397078317e-97 | -0.656119483933395 | 0.264 | 0.402 | 1.88345538740857e-93 | 1 | MARCO |
| **TIMP2** | 2.624238720515e-97 | 0.293393307021528 | 0.512 | 0.265 | 3.32254864404405e-93 | 1 | TIMP2 |
| **ENSGALG00000002431** | 9.9412052891286e-97 | 0.310911195659948 | 0.383 | 0.14 | 1.25865600165657e-92 | 1 | ENSGALG00000002431 |
| **IFIT5** | 1.56434340998522e-93 | 0.428469879862345 | 0.405 | 0.183 | 1.98061519138228e-89 | 1 | IFIT5 |
| **CRIP1.1** | 3.66235042055036e-93 | -0.551954637439116 | 0.062 | 0.274 | 4.63690186745882e-89 | 1 | CRIP1 |
| **TPP1** | 4.6437764142008e-91 | 0.331815187551193 | 0.66 | 0.378 | 5.87948531801963e-87 | 1 | TPP1 |
| **JUN.1** | 8.11377496905553e-90 | 0.508342449180966 | 0.649 | 0.253 | 1.02728504883212e-85 | 1 | JUN |
| **RNF213** | 2.34330691721311e-88 | 0.266655275822043 | 0.342 | 0.129 | 2.96686088788352e-84 | 1 | RNF213 |
| **LYN** | 6.51685816803826e-88 | 0.301238385251099 | 0.643 | 0.394 | 8.25099412655324e-84 | 1 | LYN |
| **FAM49A.1** | 3.24953113768172e-87 | 0.314897140840056 | 0.661 | 0.356 | 4.11423137341883e-83 | 1 | FAM49A |
| **HSPA5.1** | 3.07164134384642e-85 | 0.493915892972227 | 0.611 | 0.36 | 3.88900510544396e-81 | 1 | HSPA5 |
| **MBNL3** | 2.00484247101488e-83 | 0.288196311079994 | 0.582 | 0.336 | 2.53833105255194e-79 | 1 | MBNL3 |
| **ENSGALG00000046720** | 1.75148866657998e-81 | 0.258323986708645 | 0.397 | 0.176 | 2.21755980075691e-77 | 1 | ENSGALG00000046720 |
| **S100A10.1** | 9.20259813207624e-81 | -0.577384402038283 | 0.258 | 0.411 | 1.16514094950217e-76 | 1 | S100A10 |
| **FOS.1** | 4.91105876841344e-78 | 0.525925451760583 | 0.723 | 0.329 | 6.21789150668825e-74 | 1 | FOS |
| **ENSGALG00000030007.1** | 2.23187140142206e-76 | -0.437760626499343 | 0.04 | 0.252 | 2.82577238134047e-72 | 1 | ENSGALG00000030007 |
| **ENSGALG00000005257** | 1.69955922668941e-74 | 0.339390512766082 | 0.553 | 0.275 | 2.15181193691146e-70 | 1 | ENSGALG00000005257 |
| **TMSB4X.1** | 2.75148029190445e-72 | -0.338952360524697 | 0.985 | 0.995 | 3.48364919758023e-68 | 1 | TMSB4X |
| **BTG2.1** | 5.91073853123221e-72 | 0.399394938154112 | 0.603 | 0.26 | 7.4835860543931e-68 | 1 | BTG2 |
| **RPL39L.1** | 9.25494335413233e-72 | -0.365108303094744 | 0.768 | 0.842 | 1.17176837806669e-67 | 1 | RPL39L |
| **CD74** | 7.40537122252536e-68 | 0.342379273702299 | 1 | 0.994 | 9.37594050483936e-64 | 1 | CD74 |
| **GRN** | 1.84753069436804e-67 | 0.340863492939398 | 0.839 | 0.588 | 2.33915861213938e-63 | 1 | GRN |
| **CTSB** | 2.83380944099206e-65 | 0.301241529834095 | 0.592 | 0.337 | 3.58788613324004e-61 | 1 | CTSB |
| **RPS11.1** | 5.03663384567254e-64 | -0.452404336093945 | 0.784 | 0.786 | 6.376882112006e-60 | 1 | RPS11 |
| **ENSGALG00000015183.1** | 6.95173976356013e-63 | -0.314298133190609 | 0.129 | 0.33 | 8.80159771464348e-59 | 1 | ENSGALG00000015183 |
| **ENSGALG00000045199.1** | 2.34744039188356e-62 | -0.28703434199695 | 0.07 | 0.301 | 2.97209428016377e-58 | 1 | ENSGALG00000045199 |
| **TSC22D4.1** | 7.55166265242122e-62 | 0.334976310133104 | 0.555 | 0.229 | 9.56116008423051e-58 | 1 | TSC22D4 |
| **PTGS2** | 1.31296028570172e-59 | 0.296885165535599 | 0.284 | 0.088 | 1.66233901772695e-55 | 1 | PTGS2 |
| **BLB2** | 6.73750835341711e-57 | 0.307514596805033 | 0.966 | 0.844 | 8.53035932626141e-53 | 1 | BLB2 |
| **RPS15.1** | 2.93349777789933e-56 | -0.376077163208527 | 0.747 | 0.775 | 3.71410153659834e-52 | 1 | RPS15 |
| **HTRA1.1** | 3.80376429978433e-56 | -0.29828760478413 | 0.073 | 0.255 | 4.81594597995694e-52 | 1 | HTRA1 |
| **RPLP1.1** | 1.07467811300577e-55 | -0.357039308048767 | 0.876 | 0.911 | 1.36064995887661e-51 | 1 | RPLP1 |
| **HSP90B1.1** | 7.45156082239241e-54 | 0.304766360191949 | 0.577 | 0.413 | 9.43442115723103e-50 | 1 | HSP90B1 |
| **RPS26.1** | 2.91473321199326e-53 | -0.361598731533691 | 0.661 | 0.72 | 3.69034371970467e-49 | 1 | RPS26 |
| **SH3BGRL3.1** | 4.24595233643386e-51 | -0.27334460933815 | 0.245 | 0.42 | 5.37580025315891e-47 | 1 | SH3BGRL3 |
| **ATP6V0C.1** | 2.74301730627791e-50 | 0.317588985654104 | 0.77 | 0.514 | 3.47293421147846e-46 | 1 | ATP6V0C |
| **RPS19.1** | 3.4392425257275e-49 | -0.358613363456324 | 0.844 | 0.85 | 4.35442496182359e-45 | 1 | RPS19 |
| **RPL27A.1** | 4.56843009060913e-49 | -0.359110742187978 | 0.627 | 0.678 | 5.78408933772023e-45 | 1 | RPL27A |
| **RPS17.1** | 1.28151818752768e-48 | -0.335082457250533 | 0.809 | 0.828 | 1.62253017722879e-44 | 1 | RPS17 |
| **TMSB15B.1** | 8.82925930250003e-48 | 0.343171463540402 | 0.897 | 0.674 | 1.11787252028953e-43 | 1 | TMSB15B |
| **RPS8.1** | 8.46155659015524e-47 | -0.320986081776829 | 0.8 | 0.826 | 1.07131767987956e-42 | 1 | RPS8 |
| **RPL8.1** | 7.87105376975257e-46 | -0.320883649399039 | 0.578 | 0.644 | 9.96554117788373e-42 | 1 | RPL8 |
| **ENSGALG00000037441.1** | 1.09792881887574e-45 | -0.339785603830639 | 0.774 | 0.779 | 1.39008767757857e-41 | 1 | ENSGALG00000037441 |
| **RPLP2.1** | 2.31608093028102e-45 | -0.352010691206708 | 0.758 | 0.778 | 2.9323900658288e-41 | 1 | RPLP2 |
| **HMOX1** | 2.25465893491276e-44 | 0.408920971542593 | 0.54 | 0.322 | 2.85462367749305e-40 | 1 | HMOX1 |
| **PPDPF.1** | 5.11731347995031e-44 | -0.457755375282168 | 0.35 | 0.364 | 6.47903059696509e-40 | 1 | PPDPF |
| **RPS3A.1** | 9.18308481954273e-43 | -0.31675481206557 | 0.691 | 0.725 | 1.16267036900231e-38 | 1 | RPS3A |
| **LY86.1** | 3.51950759654427e-42 | -0.333515805450597 | 0.141 | 0.264 | 4.4560485679847e-38 | 1 | LY86 |
| **RPL35A.1** | 1.23020430884152e-41 | -0.309577452497914 | 0.714 | 0.742 | 1.55756167542425e-37 | 1 | RPL35A |

| | | | | | | | |
|---|---|---|---|---|---|---|---|
| **RPL17.1** | 2.46455735469761e-41 | -0.297347142061275 | 0.697 | 0.761 | 3.12037606678264e-37 | 1 | RPL17 |
| **RPL24.1** | 6.21935020626899e-41 | -0.306810504529635 | 0.634 | 0.694 | 7.87431929615716e-37 | 1 | RPL24 |
| **RPS14.1** | 3.38849392571879e-39 | -0.311685035871883 | 0.805 | 0.799 | 4.29017215935256e-35 | 1 | RPS14 |
| **RPL21.1** | 3.50134674462248e-39 | -0.291857657563513 | 0.825 | 0.829 | 4.43305511336652e-35 | 1 | RPL21 |
| **RPL35.1** | 3.69285309172002e-39 | -0.278363970175315 | 0.554 | 0.632 | 4.67552129942672e-35 | 1 | RPL35 |
| **RPL37A.1** | 6.70684937234867e-39 | -0.300261017344579 | 0.742 | 0.751 | 8.49154199033065e-35 | 1 | RPL37A |
| **RPS12.1** | 1.55818133647796e-38 | -0.290227522206156 | 0.47 | 0.548 | 1.97281339011475e-34 | 1 | RPS12 |
| **RPS24.1** | 2.46933044429008e-38 | -0.289042638202929 | 0.657 | 0.686 | 3.12641927551567e-34 | 1 | RPS24 |
| **CTSD** | 5.6744380705626e-38 | 0.443269586342544 | 0.565 | 0.473 | 7.18440604113931e-34 | 1 | CTSD |
| **RPL19.1** | 9.33824279348049e-38 | -0.286446957937328 | 0.622 | 0.654 | 1.18231492008256e-33 | 1 | RPL19 |
| **RPS27A.1** | 7.93558567105574e-37 | -0.299325460914522 | 0.687 | 0.707 | 1.00472450181237e-32 | 1 | RPS27A |
| **RPS20.1** | 8.62064829177421e-37 | -0.275993899183325 | 0.731 | 0.755 | 1.09146028022153e-32 | 1 | RPS20 |
| **EXFABP.1** | 9.91992728744821e-37 | -0.832465149258037 | 0.146 | 0.252 | 1.25596199386382e-32 | 1 | EXFABP |
| **S100A6.1** | 1.51476485572505e-36 | -0.493637044793047 | 0.244 | 0.296 | 1.91784378383349e-32 | 1 | S100A6 |
| **ITGB1BP3.1** | 2.58587649010947e-36 | -0.263171373597961 | 0.444 | 0.552 | 3.2739782241276e-32 | 1 | ITGB1BP3 |
| **RPS4Y1.1** | 4.01264875165437e-36 | -0.270205810618436 | 0.471 | 0.543 | 5.0804145844696e-32 | 1 | RPS4Y1 |
| **RPS15A.1** | 4.90488886000795e-36 | -0.286617944612446 | 0.767 | 0.783 | 6.21007978565606e-32 | 1 | RPS15A |
| **RPS28.1** | 1.01471065955644e-35 | -0.27558340732097 | 0.557 | 0.613 | 1.28472516606441e-31 | 1 | RPS28 |
| **RPL32.1** | 1.24502801654852e-35 | -0.271222532575019 | 0.647 | 0.671 | 1.57632997175208e-31 | 1 | RPL32 |
| **RPL36AL.1** | 6.5878116802827e-35 | -0.263298245228104 | 0.569 | 0.634 | 8.34082836840593e-31 | 1 | RPL36AL |
| **RPL26L1.1** | 1.8852662848066e-34 | -0.285437590262025 | 0.622 | 0.643 | 2.38693564319363e-30 | 1 | RPL26L1 |
| **ENSGALG00000026970.1** | 2.19620410891844e-34 | -0.274558083849756 | 0.426 | 0.558 | 2.78061402230164e-30 | 1 | ENSGALG00000026970 |
| **GNB2L1.1** | 4.25280749113713e-34 | -0.286840105322935 | 0.603 | 0.635 | 5.38447956452871e-30 | 1 | GNB2L1 |
| **RPL36.1** | 5.35511332607374e-34 | -0.262625402224486 | 0.471 | 0.548 | 6.78010898214197e-30 | 1 | RPL36 |
| **RPL15.1** | 1.7112073128005e-33 | -0.272875983613182 | 0.77 | 0.772 | 2.16655957873671e-29 | 1 | RPL15 |
| **RPS16.1** | 1.90276908994421e-33 | -0.272650254233997 | 0.64 | 0.663 | 2.40909594477836e-29 | 1 | RPS16 |
| **EEF1D.1** | 2.66235522739713e-33 | -0.282220689648447 | 0.415 | 0.47 | 3.37080795340751e-29 | 1 | EEF1D |
| **RPL13.1** | 4.19133941587886e-33 | -0.266833625021052 | 0.6 | 0.635 | 5.30665483444422e-29 | 1 | RPL13 |
| **RPL23.1** | 1.17344000782126e-32 | -0.273844080096887 | 0.795 | 0.797 | 1.4856923939025e-28 | 1 | RPL23 |
| **RPL37.1** | 1.987101087773e-32 | -0.277619874737822 | 0.708 | 0.711 | 2.51586868722939e-28 | 1 | RPL37 |
| **FTH1** | 5.18164713312241e-32 | 0.260138716174388 | 0.999 | 0.989 | 6.56048343524628e-28 | 1 | FTH1 |
| **RPL7A.1** | 4.45421732712811e-31 | -0.260049425052025 | 0.572 | 0.592 | 5.6394845578769e-27 | 1 | RPL7A |
| **TPT1.1** | 1.67538804873765e-29 | -0.278988555903191 | 0.826 | 0.795 | 2.12119923506739e-25 | 1 | TPT1 |
| **RPS10.1** | 1.05224961446241e-28 | -0.256783150969003 | 0.625 | 0.645 | 1.33225323687085e-24 | 1 | RPS10 |
| **RPL34.1** | 3.44317626357022e-27 | -0.252114476045318 | 0.73 | 0.724 | 4.35940546730625e-23 | 1 | RPL34 |
| **FTH1.1** | 0 | -1.27904773403702 | 0.978 | 0.993 | 0 | 2 | FTH1 |
| **IFI30.1** | 0 | -1.47338808305559 | 0.38 | 0.888 | 0 | 2 | IFI30 |
| **CD74.1** | 2.12006436293163e-257 | -0.760603064716941 | 0.988 | 0.996 | 2.68421348990774e-253 | 2 | CD74 |
| **BLB1.1** | 5.7573641318223e-239 | -0.842189086930395 | 0.664 | 0.888 | 7.28939872730021e-235 | 2 | BLB1 |
| **CTSS.1** | 8.23312311932366e-220 | -0.951262323078631 | 0.142 | 0.728 | 1.04239571813757e-215 | 2 | CTSS |
| **TMSB15B.2** | 7.63462797482715e-215 | -0.871746767552239 | 0.356 | 0.761 | 9.66620247892865e-211 | 2 | TMSB15B |
| **ENSGALG00000011190.2** | 1.05770321390804e-185 | 0.958716435254397 | 0.947 | 0.315 | 1.33915803912896e-181 | 2 | ENSGALG00000011190 |
| **GRN.1** | 1.16463526295414e-179 | -0.763828306644568 | 0.198 | 0.689 | 1.47454470642623e-175 | 2 | GRN |
| **SAT1.2** | 2.08463341559177e-163 | -0.804057125997857 | 0.137 | 0.615 | 2.63935436748074e-159 | 2 | SAT1 |
| **TMSB4X.2** | 5.20384145044592e-163 | 0.538406151026107 | 1 | 0.993 | 6.58858366040958e-159 | 2 | TMSB4X |
| **BLB2.1** | 3.96535620263e-158 | -0.626973596799696 | 0.72 | 0.884 | 5.02053748814984e-154 | 2 | BLB2 |
| **C1QC.1** | 1.80251409011289e-154 | -0.852328085136441 | 0.065 | 0.615 | 2.28216308949194e-150 | 2 | C1QC |
| **HSP90AA1.2** | 3.9396247433312e-134 | -0.703598413674028 | 0.67 | 0.804 | 4.98795888753164e-130 | 2 | HSP90AA1 |
| **PNRC1.2** | 5.870059854541e-128 | -0.606188247784601 | 0.213 | 0.59 | 7.43208278183436e-124 | 2 | PNRC1 |
| **ENSGALG00000015183.2** | 3.48782286866739e-127 | 0.507448446962926 | 0.674 | 0.244 | 4.41593253401979e-123 | 2 | ENSGALG00000015183 |
| **SPIC.1** | 3.72916415954804e-124 | 0.535510567835057 | 0.63 | 0.144 | 4.72149474240378e-120 | 2 | SPIC |
| **COX3.1** | 1.153479074341e-121 | -0.519058380906631 | 0.983 | 0.99 | 1.46041985602314e-117 | 2 | COX3 |
| **EXFABP.2** | 2.6343032218937e-119 | -1.37031231224093 | 0.043 | 0.262 | 3.33529130923961e-115 | 2 | EXFABP |
| **RPS11.2** | 2.59580849208575e-117 | -0.68726806709437 | 0.689 | 0.799 | 3.28655313182977e-113 | 2 | RPS11 |

| | | | | | | | |
|---|---|---|---|---|---|---|---|
| **GRP** | 4.05951789614005e-117 | 0.479770518029825 | 0.557 | 0.112 | 5.13975560830292e-113 | 2 | GRP |
| **PPDPF.2** | 1.29899053464897e-114 | -0.865316864977852 | 0.052 | 0.405 | 1.64465191591906e-110 | 2 | PPDPF |
| **ATP6V0C.2** | 1.39722472357205e-113 | -0.568359714964913 | 0.287 | 0.593 | 1.76902622251458e-109 | 2 | ATP6V0C |
| **SLC40A1.2** | 1.72771004243251e-107 | -0.623275434859051 | 0.069 | 0.473 | 2.18745368472381e-103 | 2 | SLC40A1 |
| **RPS19.2** | 1.15304198899969e-106 | -0.598894169662699 | 0.775 | 0.86 | 1.45986646227251e-102 | 2 | RPS19 |
| **ITM2A.1** | 1.32011456577529e-102 | -0.531825105991975 | 0.286 | 0.577 | 1.67139705172809e-98 | 2 | ITM2A |
| **TGM4.2** | 6.45663360773764e-102 | 0.970653417661379 | 0.967 | 0.248 | 8.17474381075663e-98 | 2 | TGM4 |
| **C1QB.1** | 4.29737929578753e-101 | -0.643955551172922 | 0.046 | 0.492 | 5.44091192639659e-97 | 2 | C1QB |
| **FOS.2** | 1.17612312622799e-100 | -0.608776059092394 | 0.155 | 0.426 | 1.48908949011725e-96 | 2 | FOS |
| **S100A6.2** | 7.77293368639901e-98 | -0.889384545399389 | 0.037 | 0.323 | 9.84131134034979e-94 | 2 | S100A6 |
| **TNFSF13B** | 1.25158317429964e-97 | 0.349705464810893 | 0.469 | 0.113 | 1.58462945698078e-93 | 2 | TNFSF13B |
| **HMOX1.1** | 5.55324056190711e-97 | -0.65061779697509 | 0.053 | 0.4 | 7.03095787543059e-93 | 2 | HMOX1 |
| **HSPA8.2** | 1.68046544551252e-96 | -0.554560547475591 | 0.439 | 0.649 | 2.1276373005634e-92 | 2 | HSPA8 |
| **TPT1.2** | 6.57124407182156e-95 | -0.571558175160137 | 0.717 | 0.812 | 8.31985211933327e-91 | 2 | TPT1 |
| **ENSGALG00000014585.1** | 5.3035449803002e-93 | -0.578849062276315 | 0.053 | 0.429 | 6.71481829955808e-89 | 2 | ENSGALG00000014585 |
| **HSPA5.2** | 1.08248574223634e-92 | -0.54645774536594 | 0.148 | 0.437 | 1.37053519824543e-88 | 2 | HSPA5 |
| **TPP1.1** | 4.12480307488648e-91 | -0.492082778959877 | 0.077 | 0.473 | 5.22241317311377e-87 | 2 | TPP1 |
| **RPLP1.2** | 3.32871868370581e-90 | -0.514867588131428 | 0.862 | 0.912 | 4.21449072543992e-86 | 2 | RPLP1 |
| **ATP6.1** | 3.7687605603405e-90 | -0.468740867275941 | 0.949 | 0.974 | 4.7716277454471e-86 | 2 | ATP6 |
| **MT-CO1.1** | 7.55631185114781e-90 | -0.476983311231906 | 0.965 | 0.978 | 9.56704643473824e-86 | 2 | MT-CO1 |
| **RPLP2.2** | 9.37673265264297e-90 | -0.559928228290101 | 0.688 | 0.787 | 1.18718812115113e-85 | 2 | RPLP2 |
| **CXCL13.2** | 1.11460697123296e-89 | 1.29750747793605 | 0.999 | 0.34 | 1.41120388627805e-85 | 2 | CXCL13 |
| **C1QA.1** | 2.40239501048817e-89 | -0.454554465959776 | 0.188 | 0.506 | 3.04167232277907e-85 | 2 | C1QA |
| **COII** | 1.68888770218364e-87 | -0.434983148433492 | 0.977 | 0.986 | 2.13830071973471e-83 | 2 | COII |
| **ASS1** | 7.751867329339e-87 | -0.526014700348741 | 0.057 | 0.421 | 9.81463922567611e-83 | 2 | ASS1 |
| **MAFB.2** | 2.62357272542322e-86 | -0.509814147564934 | 0.435 | 0.645 | 3.32170542765834e-82 | 2 | MAFB |
| **ND3** | 4.46397538668521e-86 | -0.487507564466882 | 0.682 | 0.843 | 5.65183923708214e-82 | 2 | ND3 |
| **RPL34.2** | 2.21247326671687e-85 | -0.514122511124896 | 0.589 | 0.745 | 2.80121240299023e-81 | 2 | RPL34 |
| **CSTA** | 2.31715859436591e-85 | 0.265492146110855 | 0.344 | 0.077 | 2.93375449632668e-81 | 2 | CSTA |
| **CTSD.1** | 5.01254418362618e-85 | -0.661714117124048 | 0.367 | 0.505 | 6.3463821908891e-81 | 2 | CTSD |
| **RPS23.1** | 1.13584151882019e-84 | -0.505410414905543 | 0.554 | 0.722 | 1.43808894697824e-80 | 2 | RPS23 |
| **RPS26.2** | 2.78359173402227e-81 | -0.505834176105574 | 0.605 | 0.726 | 3.5243054944456e-77 | 2 | RPS26 |
| **TXN** | 3.01563536219709e-81 | -0.458867563333478 | 0.258 | 0.526 | 3.81809593207774e-77 | 2 | TXN |
| **RPL23.2** | 2.88170003754834e-79 | -0.488411139352944 | 0.717 | 0.808 | 3.64852041753995e-75 | 2 | RPL23 |
| **JUN.2** | 5.09006985566176e-79 | -0.551626772303103 | 0.036 | 0.357 | 6.44453744425336e-75 | 2 | JUN |
| **LSP1P1** | 1.04839627168265e-78 | -0.435686403448663 | 0.105 | 0.407 | 1.3273745195774e-74 | 2 | LSP1P1 |
| **GNB2L1.2** | 9.49530509213951e-77 | -0.489570512418098 | 0.491 | 0.649 | 1.20220057771578e-72 | 2 | GNB2L1 |
| **RPS10.2** | 1.00106029238512e-76 | -0.481114897271338 | 0.497 | 0.662 | 1.26744243618881e-72 | 2 | RPS10 |
| **RPS14.2** | 2.10060228416803e-76 | -0.494271265135326 | 0.752 | 0.807 | 2.65957255198514e-72 | 2 | RPS14 |
| **CTSB.1** | 3.92092909684027e-76 | -0.487766752038379 | 0.038 | 0.426 | 4.96428832950947e-72 | 2 | CTSB |
| **CTSA** | 7.15435626676514e-76 | -0.440929790020149 | 0.058 | 0.406 | 9.05813046935134e-72 | 2 | CTSA |
| **RPS15.2** | 2.63852268435722e-75 | -0.491300108081049 | 0.707 | 0.78 | 3.34063357066468e-71 | 2 | RPS15 |
| **PSAP** | 2.61143646028137e-74 | -0.397634830141539 | 0.095 | 0.412 | 3.30633970236224e-70 | 2 | PSAP |
| **BTG2.2** | 2.21400424377363e-73 | -0.442232642983837 | 0.084 | 0.348 | 2.8031507730418e-69 | 2 | BTG2 |
| **SH3BGRL3.2** | 5.45932698839416e-73 | 0.34959239834937 | 0.658 | 0.354 | 6.91205390000584e-69 | 2 | SH3BGRL3 |
| **RPL27A.2** | 6.82058953650724e-72 | -0.489898925355883 | 0.584 | 0.682 | 8.63554841217181e-68 | 2 | RPL27A |
| **DDX5.2** | 8.12431237351079e-71 | -0.424125699765179 | 0.332 | 0.545 | 1.0286191896102e-66 | 2 | DDX5 |
| **ENSGALG00000037441.2** | 1.23083208673227e-69 | -0.474253969128197 | 0.713 | 0.788 | 1.55835650501172e-65 | 2 | ENSGALG00000037441 |
| **RPL37A.2** | 2.47463813013581e-69 | -0.455900868424227 | 0.656 | 0.763 | 3.13313933656495e-65 | 2 | RPL37A |
| **RPS12.2** | 3.97009456242575e-69 | -0.441728158768128 | 0.383 | 0.556 | 5.02653672548724e-65 | 2 | RPS12 |
| **CTSZ** | 4.62370561896656e-69 | -0.375155150529377 | 0.136 | 0.414 | 5.85407368417356e-65 | 2 | CTSZ |
| **ZEB2.2** | 8.88134525239875e-68 | -0.418734910334226 | 0.051 | 0.346 | 1.12446712240621e-63 | 2 | ZEB2 |
| **ENSGALG00000004509.2** | 9.02905059808778e-68 | -0.404828858645993 | 0.801 | 0.875 | 1.14316809622389e-63 | 2 | ENSGALG00000004509 |
| **RPL31.1** | 1.10430998890657e-67 | -0.424258966019984 | 0.454 | 0.632 | 1.39816687695461e-63 | 2 | RPL31 |

| | | | | | | | |
|---|---|---|---|---|---|---|---|
| **RPL26L1.2** | 6.98232319438482e-67 | -0.451834125630688 | 0.543 | 0.654 | 8.84031939641063e-63 | 2 | RPL26L1 |
| **CTSC** | 1.58760758438393e-66 | -0.362695228091515 | 0.074 | 0.38 | 2.01006996258849e-62 | 2 | CTSC |
| **TNFAIP2** | 4.09200321659515e-66 | 0.345858800910562 | 0.485 | 0.134 | 5.18088527253112e-62 | 2 | TNFAIP2 |
| **LY86.2** | 7.57593889165298e-66 | -0.476025746684117 | 0.049 | 0.271 | 9.59189623072184e-62 | 2 | LY86 |
| **RPS27A.2** | 7.76613964595525e-66 | -0.454010267470746 | 0.623 | 0.715 | 9.83270940574394e-62 | 2 | RPS27A |
| **LAMP1** | 1.30220964699878e-65 | -0.356528650640897 | 0.077 | 0.378 | 1.64872763406516e-61 | 2 | LAMP1 |
| **MARCO.2** | 2.18543287570166e-65 | 0.730706870960969 | 0.903 | 0.306 | 2.76697656392587e-61 | 2 | MARCO |
| **NFKBIA.2** | 4.93675587389582e-65 | -0.37808970314411 | 0.111 | 0.367 | 6.2504266119395e-61 | 2 | NFKBIA |
| **RPL15.2** | 1.15716612767498e-64 | -0.432756964915369 | 0.706 | 0.781 | 1.46508803424929e-60 | 2 | RPL15 |
| **RPL21.2** | 1.80179633123454e-64 | -0.425774562798541 | 0.788 | 0.834 | 2.28125433497605e-60 | 2 | RPL21 |
| **EEF1D.2** | 3.70351476811732e-64 | -0.439886325623989 | 0.315 | 0.481 | 4.68902004791334e-60 | 2 | EEF1D |
| **CD47.2** | 3.71307237097224e-64 | -0.354120616039801 | 0.13 | 0.393 | 4.70112092888796e-60 | 2 | CD47 |
| **EEF1A1.1** | 4.11435863754529e-64 | -0.421665171436165 | 0.75 | 0.828 | 5.20918947099609e-60 | 2 | EEF1A1 |
| **S100A10.2** | 8.60370977195955e-64 | -0.57230376879881 | 0.292 | 0.399 | 1.0893156942278e-59 | 2 | S100A10 |
| **RPS15A.2** | 1.32607648052085e-63 | -0.433251500685576 | 0.725 | 0.788 | 1.67894543198745e-59 | 2 | RPS15A |
| **ND6** | 1.42915402682444e-63 | -0.403029537769273 | 0.767 | 0.888 | 1.80945191336242e-59 | 2 | ND6 |
| **RPS16.2** | 5.98219130547288e-63 | -0.426299033628622 | 0.556 | 0.674 | 7.57405241185922e-59 | 2 | RPS16 |
| **BTG1.1** | 1.71531524569904e-62 | -0.352860917189892 | 0.096 | 0.356 | 2.17176063257956e-58 | 2 | BTG1 |
| **RPL37.2** | 5.93886770177643e-62 | -0.436419834927744 | 0.633 | 0.722 | 7.51920039721914e-58 | 2 | RPL37 |
| **RPS17.2** | 3.4295133518344e-61 | -0.424618853493201 | 0.791 | 0.83 | 4.34210685475753e-57 | 2 | RPS17 |
| **RPS28.2** | 4.67853996437633e-61 | -0.410231985509228 | 0.481 | 0.621 | 5.92349944889687e-57 | 2 | RPS28 |
| **RPL30.1** | 5.15577452791054e-61 | -0.404295380521287 | 0.434 | 0.596 | 6.52772612978754e-57 | 2 | RPL30 |
| **RPL35A.2** | 9.19240139742557e-60 | -0.420068903591153 | 0.665 | 0.747 | 1.16384994092805e-55 | 2 | RPL35A |
| **RPL23A.1** | 1.14220782607219e-59 | -0.408084482463118 | 0.489 | 0.63 | 1.44614932859e-55 | 2 | RPL23A |
| **RPL18A.1** | 1.86464886527817e-59 | -0.402477863005792 | 0.418 | 0.587 | 2.36083192832869e-55 | 2 | RPL18A |
| **RPS21.1** | 1.89642396622186e-58 | -0.3883930273551 | 0.38 | 0.555 | 2.4010623836335e-54 | 2 | RPS21 |
| **GSTA3** | 2.40897503452405e-58 | -0.380365606858348 | 0.035 | 0.343 | 3.0500032912109e-54 | 2 | GSTA3 |
| **TSC22D4.2** | 2.63345186894751e-58 | -0.416241797075307 | 0.031 | 0.317 | 3.33421341127445e-54 | 2 | TSC22D4 |
| **RSFR.1** | 5.2583375321303e-57 | -0.398289237407779 | 0.03 | 0.339 | 6.65758114943017e-53 | 2 | RSFR |
| **SCPEP1.1** | 7.67106980275068e-57 | -0.320817879200812 | 0.07 | 0.33 | 9.71234147726263e-53 | 2 | SCPEP1 |
| **RPS24.2** | 2.87661141154835e-56 | -0.397366635349325 | 0.593 | 0.694 | 3.64207770816137e-52 | 2 | RPS24 |
| **RPS20.2** | 5.66542330329555e-56 | -0.387423923411092 | 0.667 | 0.763 | 7.1729924443025e-52 | 2 | RPS20 |
| **BRI3** | 1.3404854910086e-55 | -0.306593293494934 | 0.155 | 0.407 | 1.69718868016599e-51 | 2 | BRI3 |
| **CRIP1.2** | 2.3591727429366e-54 | -0.471064350703379 | 0.117 | 0.257 | 2.98694860983203e-50 | 2 | CRIP1 |
| **MGST1.1** | 6.7127495527614e-54 | -0.37268647653809 | 0.073 | 0.251 | 8.4990122087512e-50 | 2 | MGST1 |
| **CYTB** | 1.69144258816365e-53 | -0.348535427009866 | 0.797 | 0.894 | 2.14153546087399e-49 | 2 | CYTB |
| **SPI1** | 2.15379164070848e-53 | -0.293372509387076 | 0.091 | 0.344 | 2.72691559630101e-49 | 2 | SPI1 |
| **RPL7.1** | 2.58056944361546e-53 | -0.381383376786037 | 0.476 | 0.605 | 3.26725897256154e-49 | 2 | RPL7 |
| **RPL24.2** | 5.92228305684942e-53 | -0.394181296983402 | 0.61 | 0.695 | 7.49820257827706e-49 | 2 | RPL24 |
| **RPS8.2** | 6.03778937880769e-53 | -0.385133642553546 | 0.794 | 0.826 | 7.64444513250842e-49 | 2 | RPS8 |
| **PRDX1** | 1.65157675442491e-52 | -0.301875945155997 | 0.19 | 0.424 | 2.09106132877738e-48 | 2 | PRDX1 |
| **PPIB** | 8.14340800189619e-52 | -0.296172361179314 | 0.154 | 0.391 | 1.03103688712008e-47 | 2 | PPIB |
| **HSP90B1.2** | 3.62677385562997e-51 | -0.338501393791088 | 0.258 | 0.465 | 4.59185837861311e-47 | 2 | HSP90B1 |
| **CTSV** | 1.03799990732456e-50 | -0.372073934237108 | 0.023 | 0.322 | 1.31421168266363e-46 | 2 | CTSV |
| **GAPDH.1** | 1.58782081544424e-50 | -0.358848041608018 | 0.385 | 0.544 | 2.01033993443396e-46 | 2 | GAPDH |
| **RPL12.1** | 5.09798089692683e-50 | -0.365062037437596 | 0.482 | 0.616 | 6.45455361359906e-46 | 2 | RPL12 |
| **RPL7A.2** | 5.34580834377754e-50 | -0.374156262296988 | 0.487 | 0.603 | 6.76832794405674e-46 | 2 | RPL7A |
| **RPS3A.2** | 2.27816648256478e-49 | -0.383468774980501 | 0.687 | 0.724 | 2.88438658357527e-45 | 2 | RPS3A |
| **RPL36.2** | 2.30767899860824e-49 | -0.358659204688694 | 0.42 | 0.552 | 2.92175238013789e-45 | 2 | RPL36 |
| **CST3.1** | 3.32738424079455e-49 | -0.332484777480863 | 0.605 | 0.726 | 4.21280118726998e-45 | 2 | CST3 |
| **GNS** | 3.43522738383198e-49 | -0.276163354400026 | 0.14 | 0.371 | 4.34934139066968e-45 | 2 | GNS |
| **HTRA1.2** | 3.84864873586334e-49 | 0.363142707173577 | 0.558 | 0.179 | 4.87277416447657e-45 | 2 | HTRA1 |
| **ANKRD11.1** | 1.1190735450406e-48 | -0.306612783251783 | 0.148 | 0.357 | 1.4168590153759e-44 | 2 | ANKRD11 |
| **ND5** | 1.47317562726772e-48 | -0.337131786161094 | 0.759 | 0.858 | 1.86518766168366e-44 | 2 | ND5 |

| | | | | | | | |
|---|---|---|---|---|---|---|---|
| **RPL13.2** | 4.97925196992768e-48 | -0.364318624548865 | 0.54 | 0.642 | 6.30423091912543e-44 | 2 | RPL13 |
| **ATP6AP1** | 1.45449916023828e-47 | -0.269432920481344 | 0.056 | 0.294 | 1.84154138677769e-43 | 2 | ATP6AP1 |
| **CREG1** | 1.5244510733637e-46 | -0.272987742954894 | 0.044 | 0.286 | 1.93010750398578e-42 | 2 | CREG1 |
| **RPL19.2** | 1.5608821581799e-46 | -0.359130439509261 | 0.576 | 0.659 | 1.97623290047157e-42 | 2 | RPL19 |
| **RPS3.1** | 3.29529684688226e-46 | -0.338081612399799 | 0.399 | 0.535 | 4.17217533783763e-42 | 2 | RPS3 |
| **RPS29.1** | 4.04604201548791e-46 | -0.333354353487638 | 0.397 | 0.539 | 5.12269379580924e-42 | 2 | RPS29 |
| **FGL2** | 4.39202825537768e-46 | -0.276505123818661 | 0.05 | 0.266 | 5.56074697413368e-42 | 2 | FGL2 |
| **ENSGALG00000045115.1** | 1.6354613217579e-45 | -0.355898399715085 | 0.019 | 0.302 | 2.07065757947768e-41 | 2 | ENSGALG00000045115 |
| **FAU.1** | 1.93105521369859e-45 | -0.330572493919525 | 0.396 | 0.534 | 2.44490900606379e-41 | 2 | FAU |
| **ND1** | 2.41620341652995e-45 | -0.318966505490984 | 0.721 | 0.821 | 3.05915514566857e-41 | 2 | ND1 |
| **RPL32.2** | 2.04939874858221e-44 | -0.34260304669305 | 0.589 | 0.678 | 2.59474375557993e-40 | 2 | RPL32 |
| **RPS25.1** | 1.77811211438595e-43 | -0.337988840520143 | 0.546 | 0.645 | 2.25126774802405e-39 | 2 | RPS25 |
| **RPS7.1** | 2.15469875157997e-43 | -0.326927087074675 | 0.44 | 0.576 | 2.7280640893754e-39 | 2 | RPS7 |
| **SLC25A6** | 2.29376130716214e-43 | -0.276144221225753 | 0.179 | 0.358 | 2.90413119099798e-39 | 2 | SLC25A6 |
| **UBA52.1** | 2.81962650625949e-43 | -0.325493123722117 | 0.475 | 0.585 | 3.56992911957514e-39 | 2 | UBA52 |
| **RPL22.1** | 9.2247713003994e-43 | -0.322747476209253 | 0.491 | 0.612 | 1.16794829434357e-38 | 2 | RPL22 |
| **HAVCR1.1** | 3.06455217689307e-42 | -0.263202846441406 | 0.031 | 0.263 | 3.88002951116431e-38 | 2 | HAVCR1 |
| **ENSGALG00000047532.1** | 1.35004274321172e-41 | -0.255626865371469 | 0.187 | 0.386 | 1.70928911718036e-37 | 2 | ENSGALG00000047532 |
| **LIPA** | 8.87284235556509e-41 | -0.253220060301539 | 0.031 | 0.253 | 1.1233905706381e-36 | 2 | LIPA |
| **RPS13.1** | 3.98696386751412e-40 | -0.310545987413999 | 0.508 | 0.623 | 5.04789495265963e-36 | 2 | RPS13 |
| **RPL29.1** | 4.03259111184612e-40 | -0.309752739022679 | 0.549 | 0.636 | 5.10566360670837e-36 | 2 | RPL29 |
| **RPL27.1** | 5.80939383436442e-40 | -0.311688278974829 | 0.438 | 0.549 | 7.35527353368879e-36 | 2 | RPL27 |
| **CD81.1** | 1.81847425419492e-39 | -0.258260336880095 | 0.114 | 0.282 | 2.30237025323619e-35 | 2 | CD81 |
| **HMGB1** | 1.84524176458812e-39 | -0.26730096758783 | 0.181 | 0.363 | 2.33626059814502e-35 | 2 | HMGB1 |
| **RPS6.1** | 2.11260093802741e-39 | -0.319608811952554 | 0.548 | 0.636 | 2.6747640476365e-35 | 2 | RPS6 |
| **RPL11.1** | 4.78499654021429e-39 | -0.313158404817734 | 0.587 | 0.664 | 6.05828411956532e-35 | 2 | RPL11 |
| **RPL9.1** | 5.58931728830817e-39 | -0.307921351610922 | 0.486 | 0.592 | 7.07663461872697e-35 | 2 | RPL9 |
| **RPL8.2** | 6.76185749284721e-39 | -0.330899797263266 | 0.571 | 0.642 | 8.56118777169385e-35 | 2 | RPL8 |
| **ND4** | 1.34715659648478e-38 | -0.295049905482383 | 0.668 | 0.786 | 1.70563496680938e-34 | 2 | ND4 |
| **RPS2.1** | 3.41303039132352e-37 | -0.291422230781203 | 0.528 | 0.629 | 4.32123777845471e-33 | 2 | RPS2 |
| **EEF2.1** | 3.65543105523922e-37 | -0.313309799597804 | 0.431 | 0.535 | 4.62814125903837e-33 | 2 | EEF2 |
| **RPL4.1** | 1.79915076205215e-35 | -0.27565942129639 | 0.265 | 0.397 | 2.27790477983422e-31 | 2 | RPL4 |
| **RPL36AL.2** | 2.0427316664894e-35 | -0.298322761302202 | 0.547 | 0.635 | 2.58630256294223e-31 | 2 | RPL36AL |
| **PABPC1.1** | 3.50818975723667e-35 | -0.266614604349952 | 0.478 | 0.602 | 4.44171905163735e-31 | 2 | PABPC1 |
| **RPL6.1** | 1.91078502222971e-34 | -0.276985995841997 | 0.477 | 0.577 | 2.41924491664503e-30 | 2 | RPL6 |
| **RPS4Y1.2** | 3.39263698514484e-33 | -0.290104342653967 | 0.46 | 0.542 | 4.29541768689188e-29 | 2 | RPS4Y1 |
| **RPL10A.1** | 3.79877443655647e-33 | -0.274483768999296 | 0.463 | 0.559 | 4.80962831412415e-29 | 2 | RPL10A |
| **RPL17.2** | 9.627590739822366e-33 | -0.295596526276453 | 0.741 | 0.752 | 1.21894926356891e-28 | 2 | RPL17 |
| **ENSGALG00000030007.2** | 2.11779616714668e-30 | 0.381156615544651 | 0.583 | 0.166 | 2.68134172722441e-26 | 2 | ENSGALG00000030007 |
| **RPL3.1** | 5.66364294571381e-30 | -0.250559175631748 | 0.418 | 0.516 | 7.17073833356826e-26 | 2 | RPL3 |
| **RPL35.2** | 1.54500883435477e-26 | -0.25445442910349 | 0.599 | 0.622 | 1.95613568517657e-22 | 2 | RPL35 |
| **S100A10.3** | 0 | 1.38740968997284 | 0.932 | 0.311 | 0 | 3 | S100A10 |
| **RPS11.3** | 0 | 1.18653849169421 | 0.984 | 0.758 | 0 | 3 | RPS11 |
| **RPS19.3** | 0 | 1.14228202020039 | 0.992 | 0.83 | 0 | 3 | RPS19 |
| **RPLP1.3** | 0 | 1.13296714821774 | 0.99 | 0.894 | 0 | 3 | RPLP1 |
| **RPS15.3** | 0 | 1.11145325760831 | 0.983 | 0.742 | 0 | 3 | RPS15 |
| **RPLP2.3** | 0 | 1.09347442776177 | 0.985 | 0.746 | 0 | 3 | RPLP2 |
| **LY86.3** | 0 | 1.05081178541153 | 0.83 | 0.163 | 0 | 3 | LY86 |
| **RPL27A.3** | 0 | 1.05039738978895 | 0.951 | 0.631 | 0 | 3 | RPL27A |
| **ENSGALG00000037441.3** | 0 | 1.04801382504697 | 0.977 | 0.751 | 0 | 3 | ENSGALG00000037441 |
| **RPS14.3** | 0 | 1.04512384439305 | 0.987 | 0.774 | 0 | 3 | RPS14 |
| **RPL37A.3** | 0 | 1.01995544758317 | 0.975 | 0.718 | 0 | 3 | RPL37A |
| **RPS26.3** | 0 | 1.01492448337655 | 0.965 | 0.676 | 0 | 3 | RPS26 |
| **RPS17.3** | 0 | 1.00624903124494 | 0.988 | 0.803 | 0 | 3 | RPS17 |

| | | | | | | | | |
|---|---|---|---|---|---|---|---|---|
| **RPL34.3** | | 0 | 0.997546429967307 | 0.963 | 0.693 | 0 | 3 | RPL34 |
| **GNB2L1.3** | | 0 | 0.99743927821264 | 0.96 | 0.584 | 0 | 3 | GNB2L1 |
| **RPL37.3** | | 0 | 0.986955349913359 | 0.969 | 0.675 | 0 | 3 | RPL37 |
| **RPL23.3** | | 0 | 0.966171810921028 | 0.977 | 0.772 | 0 | 3 | RPL23 |
| **TPT1.3** | | 0 | 0.960670843133842 | 0.976 | 0.776 | 0 | 3 | TPT1 |
| **RPS27A.3** | | 0 | 0.951744214785595 | 0.964 | 0.668 | 0 | 3 | RPS27A |
| **RPS10.3** | | 0 | 0.951617787935649 | 0.965 | 0.598 | 0 | 3 | RPS10 |
| **RPL35A.3** | | 0 | 0.947101748913782 | 0.974 | 0.705 | 0 | 3 | RPL35A |
| **RPS3A.3** | | 0 | 0.94514250732466 | 0.969 | 0.685 | 0 | 3 | RPS3A |
| **RPS8.3** | | 0 | 0.944981736429041 | 0.985 | 0.799 | 0 | 3 | RPS8 |
| **RPL26L1.3** | | 0 | 0.944134758173715 | 0.95 | 0.597 | 0 | 3 | RPL26L1 |
| **EEF1D.3** | | 0 | 0.944127142084523 | 0.902 | 0.4 | 0 | 3 | EEF1D |
| **RPL24.3** | | 0 | 0.942950588304163 | 0.96 | 0.647 | 0 | 3 | RPL24 |
| **RPL21.3** | | 0 | 0.940128400430242 | 0.979 | 0.807 | 0 | 3 | RPL21 |
| **RPS12.3** | | 0 | 0.934175106617973 | 0.929 | 0.481 | 0 | 3 | RPS12 |
| **RPS15A.3** | | 0 | 0.934166064925359 | 0.968 | 0.754 | 0 | 3 | RPS15A |
| **RPS23.2** | | 0 | 0.933786760719462 | 0.966 | 0.665 | 0 | 3 | RPS23 |
| **RPL15.3** | | 0 | 0.931929925129564 | 0.974 | 0.744 | 0 | 3 | RPL15 |
| **RPS28.3** | | 0 | 0.929755500657331 | 0.945 | 0.557 | 0 | 3 | RPS28 |
| **EEF1A1.2** | | 0 | 0.927309897782744 | 0.971 | 0.797 | 0 | 3 | EEF1A1 |
| **RPL13.3** | | 0 | 0.9058855461935 | 0.946 | 0.586 | 0 | 3 | RPL13 |
| **RPL8.3** | | 0 | 0.905470571576765 | 0.948 | 0.59 | 0 | 3 | RPL8 |
| **RPL36.3** | | 0 | 0.896138471496638 | 0.923 | 0.482 | 0 | 3 | RPL36 |
| **MT-CO1.2** | | 0 | 0.889333597600432 | 0.998 | 0.974 | 0 | 3 | MT-CO1 |
| **RPS24.3** | | 0 | 0.888380930842364 | 0.947 | 0.645 | 0 | 3 | RPS24 |
| **RPL23A.2** | | 0 | 0.888328038700893 | 0.932 | 0.569 | 0 | 3 | RPL23A |
| **RPS16.3** | | 0 | 0.887753809529168 | 0.956 | 0.619 | 0 | 3 | RPS16 |
| **ANXA2** | | 0 | 0.884888973345885 | 0.742 | 0.08 | 0 | 3 | ANXA2 |
| **RPL7A.3** | | 0 | 0.878790319289657 | 0.916 | 0.544 | 0 | 3 | RPL7A |
| **RPL7.2** | | 0 | 0.877243108908742 | 0.917 | 0.544 | 0 | 3 | RPL7 |
| **RPL19.3** | | 0 | 0.872051892788445 | 0.942 | 0.608 | 0 | 3 | RPL19 |
| **RPL17.3** | | 0 | 0.868161187074608 | 0.96 | 0.722 | 0 | 3 | RPL17 |
| **RPS20.3** | | 0 | 0.86149595294091 | 0.965 | 0.722 | 0 | 3 | RPS20 |
| **RPS29.2** | | 0 | 0.86046927717185 | 0.895 | 0.471 | 0 | 3 | RPS29 |
| **RPL30.2** | | 0 | 0.858893354651682 | 0.909 | 0.53 | 0 | 3 | RPL30 |
| **RPL12.2** | | 0 | 0.850021088196888 | 0.931 | 0.554 | 0 | 3 | RPL12 |
| **RPL18A.2** | | 0 | 0.849498759857385 | 0.914 | 0.518 | 0 | 3 | RPL18A |
| **ND3.1** | | 0 | 0.848760207299186 | 0.992 | 0.799 | 0 | 3 | ND3 |
| **RPS6.2** | | 0 | 0.845147796482596 | 0.94 | 0.582 | 0 | 3 | RPS6 |
| **GAPDH.2** | | 0 | 0.835469701510984 | 0.87 | 0.477 | 0 | 3 | GAPDH |
| **RPL32.3** | | 0 | 0.829345089400613 | 0.949 | 0.628 | 0 | 3 | RPL32 |
| **RPS25.2** | | 0 | 0.82927319546273 | 0.935 | 0.592 | 0 | 3 | RPS25 |
| **RPS21.2** | | 0 | 0.826857335887547 | 0.907 | 0.482 | 0 | 3 | RPS21 |
| **RPL22.2** | | 0 | 0.825171527512235 | 0.916 | 0.553 | 0 | 3 | RPL22 |
| **COX3.2** | | 0 | 0.820722292312227 | 1 | 0.987 | 0 | 3 | COX3 |
| **RPS4Y1.3** | | 0 | 0.818131436675521 | 0.907 | 0.48 | 0 | 3 | RPS4Y1 |
| **UBA52.2** | | 0 | 0.81567701322521 | 0.917 | 0.524 | 0 | 3 | UBA52 |
| **EEF2.2** | | 0 | 0.8112247191658 | 0.914 | 0.468 | 0 | 3 | EEF2 |
| **ATP6.2** | | 0 | 0.811150056226549 | 1 | 0.967 | 0 | 3 | ATP6 |
| **RPL35.3** | | 0 | 0.804973208131675 | 0.927 | 0.577 | 0 | 3 | RPL35 |
| **RPS3.2** | | 0 | 0.80244578735787 | 0.879 | 0.469 | 0 | 3 | RPS3 |
| **RPL11.2** | | 0 | 0.799042400992633 | 0.938 | 0.616 | 0 | 3 | RPL11 |
| **RPL31.2** | | 0 | 0.79308108903725 | 0.901 | 0.57 | 0 | 3 | RPL31 |
| **RPL36AL.3** | | 0 | 0.788135192543184 | 0.921 | 0.583 | 0 | 3 | RPL36AL |

| | | | | | | | |
|---|---|---|---|---|---|---|---|
| **ITM2A.2** | 0 | 0.786243266271827 | 0.927 | 0.488 | 0 | 3 | ITM2A |
| **RPL4.2** | 0 | 0.785902333586141 | 0.838 | 0.318 | 0 | 3 | RPL4 |
| **ITGB1BP3.2** | 0 | 0.783996034070477 | 0.892 | 0.486 | 0 | 3 | ITGB1BP3 |
| **RPL39L.2** | 0 | 0.763500722253398 | 0.969 | 0.811 | 0 | 3 | RPL39L |
| **FAU.2** | 0 | 0.763205721093054 | 0.873 | 0.468 | 0 | 3 | FAU |
| **RPL9.2** | 0 | 0.75668721277115 | 0.877 | 0.538 | 0 | 3 | RPL9 |
| **RPL29.2** | 0 | 0.756645274964911 | 0.92 | 0.585 | 0 | 3 | RPL29 |
| **RPS2.2** | 0 | 0.754529029398666 | 0.891 | 0.579 | 0 | 3 | RPS2 |
| **RPL14.1** | 0 | 0.71337642329728 | 0.874 | 0.516 | 0 | 3 | RPL14 |
| **GPX1** | 0 | 0.690404401847363 | 0.677 | 0.113 | 0 | 3 | GPX1 |
| **RPL38.1** | 0 | 0.672192640105596 | 0.868 | 0.491 | 0 | 3 | RPL38 |
| **CDK5RAP1** | 0 | 0.64959282744001 | 0.692 | 0.132 | 0 | 3 | CDK5RAP1 |
| **RPLP0** | 0 | 0.644238676089206 | 0.719 | 0.207 | 0 | 3 | RPLP0 |
| **C3** | 0 | 0.516744541501453 | 0.546 | 0.084 | 0 | 3 | C3 |
| **COII.1** | 5.53445451770866e-304 | 0.738759624119961 | 1 | 0.983 | 7.00717286487094e-300 | 3 | COII |
| **RPL6.2** | 9.46857781304473e-304 | 0.707925140330937 | 0.905 | 0.518 | 1.19881663690959e-299 | 3 | RPL6 |
| **RPL10A.2** | 9.56802970105957e-304 | 0.723266713339812 | 0.863 | 0.504 | 1.21140824045115e-299 | 3 | RPL10A |
| **RPL27.2** | 2.58143420757333e-298 | 0.750836104624658 | 0.885 | 0.488 | 3.26835385020859e-294 | 3 | RPL27 |
| **RPS13.2** | 2.10532622795832e-297 | 0.735503401988139 | 0.901 | 0.569 | 2.66555353721803e-293 | 3 | RPS13 |
| **MGST1.2** | 6.12152281917521e-292 | 0.895835988285457 | 0.799 | 0.151 | 7.75046004135773e-288 | 3 | MGST1 |
| **RPL3.2** | 2.57818576558039e-288 | 0.681291710110858 | 0.844 | 0.457 | 3.26424099780134e-284 | 3 | RPL3 |
| **VIM.1** | 1.66610194794224e-284 | 1.03899722079615 | 0.85 | 0.125 | 2.10945167628967e-280 | 3 | VIM |
| **RPSAP58.1** | 1.70865549774462e-280 | 0.678518578120081 | 0.881 | 0.542 | 2.16332872569446e-276 | 3 | RPSAP58 |
| **RPS7.2** | 1.14442335742132e-279 | 0.736101248547656 | 0.881 | 0.514 | 1.44895441283113e-275 | 3 | RPS7 |
| **S100A4.1** | 4.49849368368534e-279 | 1.6617707182535 | 0.896 | 0.104 | 5.69554285291401e-275 | 3 | S100A4 |
| **PPDPF.3** | 3.95544110474587e-276 | 1.31725885500611 | 0.953 | 0.28 | 5.00798398271874e-272 | 3 | PPDPF |
| **RPL5.1** | 8.67682291184471e-274 | 0.693087052823857 | 0.859 | 0.435 | 1.09857254886866e-269 | 3 | RPL5 |
| **ENSGALG00000054657** | 3.49567533458157e-271 | 0.413895759469629 | 0.412 | 0.035 | 4.42587454111373e-267 | 3 | ENSGALG00000054657 |
| **HMGB1.1** | 3.95839679041713e-269 | 0.627258276370633 | 0.737 | 0.286 | 5.01172617634713e-265 | 3 | HMGB1 |
| **GNAI3** | 4.58627403291174e-268 | 0.487225657705982 | 0.593 | 0.156 | 5.80668155306956e-264 | 3 | GNAI3 |
| **NME2.1** | 8.58072876519867e-267 | 0.660280766979051 | 0.815 | 0.437 | 1.0864060689618e-262 | 3 | NME2 |
| **LSP1P1.1** | 9.03177591885804e-261 | 0.676972793280445 | 0.809 | 0.31 | 1.14351314908662e-256 | 3 | LSP1P1 |
| **ENSGALG00000052026** | 4.15091713943185e-260 | 0.775447335768737 | 0.531 | 0.052 | 5.25547619023466e-256 | 3 | ENSGALG00000052026 |
| **FABP3** | 1.08596314638478e-259 | 0.838630286247302 | 0.683 | 0.071 | 1.37493793963777e-255 | 3 | FABP3 |
| **VCAN** | 2.83006505616676e-250 | 1.12507283631736 | 0.862 | 0.084 | 3.58314536761274e-246 | 3 | VCAN |
| **CSF3R** | 3.07018886968133e-248 | 0.403685140576727 | 0.42 | 0.043 | 3.88716612790353e-244 | 3 | CSF3R |
| **URM1** | 1.07125378061971e-241 | 0.515915869687446 | 0.611 | 0.188 | 1.35631441164261e-237 | 3 | URM1 |
| **CD99** | 4.69378739908171e-240 | 0.551501140959457 | 0.547 | 0.056 | 5.94280422597735e-236 | 3 | CD99 |
| **SLC25A6.1** | 1.06462327505524e-238 | 0.578056589316349 | 0.709 | 0.285 | 1.34791952854744e-234 | 3 | SLC25A6 |
| **PABPC1.2** | 1.13218016880675e-234 | 0.593950210675886 | 0.889 | 0.546 | 1.43345331172622e-230 | 3 | PABPC1 |
| **CD3D** | 2.83340115215257e-232 | 0.500398386216054 | 0.479 | 0.09 | 3.58736919874037e-228 | 3 | CD3D |
| **ND1.1** | 5.60109571458692e-230 | 0.63122338204148 | 0.982 | 0.785 | 7.0915472842385e-226 | 3 | ND1 |
| **EMP1** | 6.79859599994772e-227 | 0.518656816495564 | 0.478 | 0.049 | 8.6077023955338e-223 | 3 | EMP1 |
| **LDHA.1** | 7.94838603041722e-224 | 0.902401405908491 | 0.792 | 0.138 | 1.00634515531112e-219 | 3 | LDHA |
| **S100A6.3** | 1.45364671404868e-223 | 1.47292794310361 | 0.952 | 0.196 | 1.84046210465703e-219 | 3 | S100A6 |
| **ENSGALG00000008518** | 2.81490135013299e-221 | 0.487521342664512 | 0.554 | 0.107 | 3.56394659940338e-217 | 3 | ENSGALG00000008518 |
| **EIF3F** | 1.58474073080875e-220 | 0.467534919816886 | 0.533 | 0.131 | 2.00644023927696e-216 | 3 | EIF3F |
| **CRIP1.3** | 3.01247139547118e-219 | 1.13310849807118 | 0.852 | 0.155 | 3.81409003380606e-215 | 3 | CRIP1 |
| **FGL2.1** | 1.08894480202024e-218 | 0.515474465236585 | 0.645 | 0.183 | 1.37871301383782e-214 | 3 | FGL2 |
| **SPTBN1** | 2.33536321714991e-218 | 0.527382183186768 | 0.527 | 0.067 | 2.95680336923351e-214 | 3 | SPTBN1 |
| **BTG1.2** | 7.52148035682998e-218 | 0.57011511377104 | 0.73 | 0.268 | 9.52294627978244e-214 | 3 | BTG1 |
| **CALM2.1** | 1.03285986412739e-217 | 0.542004180492555 | 0.755 | 0.377 | 1.30770387397169e-213 | 3 | CALM2 |
| **TPM3** | 5.35938647237419e-216 | 0.496735790693858 | 0.626 | 0.192 | 6.78551921267296e-212 | 3 | TPM3 |
| **CD81.2** | 1.26231677921676e-213 | 0.581952844203462 | 0.684 | 0.203 | 1.59821927416634e-209 | 3 | CD81 |

| | | | | | | | |
|---|---|---|---|---|---|---|---|
| **S100A11** | 5.46488375272992e-211 | 0.532004320372856 | 0.558 | 0.14 | 6.91908931933135e-207 | 3 | S100A11 |
| **CORO1C.1** | 5.30431024560247e-210 | 0.551906381501128 | 0.77 | 0.411 | 6.71578720195728e-206 | 3 | CORO1C |
| **CD74.2** | 1.22283978476085e-209 | 0.654538677970116 | 1 | 0.994 | 1.54823745148571e-205 | 3 | CD74 |
| **CXCL13.3** | 1.40603310979584e-209 | -1.57525625312755 | 0.126 | 0.462 | 1.78017852031251e-205 | 3 | CXCL13 |
| **ND5.1** | 3.62139825782138e-207 | 0.625946056651214 | 0.985 | 0.827 | 4.58505233422765e-203 | 3 | ND5 |
| **SLC1A6** | 1.00035305755843e-206 | 0.353506624254162 | 0.36 | 0.03 | 1.26654700617472e-202 | 3 | SLC1A6 |
| **ND4.1** | 8.59472646778578e-206 | 0.608175664570576 | 0.973 | 0.743 | 1.08817831808636e-201 | 3 | ND4 |
| **ALDOC** | 6.84349183093658e-202 | 0.402483982027 | 0.471 | 0.117 | 8.66454500714881e-198 | 3 | ALDOC |
| **HNRNPA1** | 1.13035236089462e-200 | 0.456210784089685 | 0.589 | 0.212 | 1.43113912412868e-196 | 3 | HNRNPA1 |
| **ANXA1.1** | 2.43146572519034e-198 | 1.36385182640736 | 0.92 | 0.104 | 3.07847875466349e-194 | 3 | ANXA1 |
| **EMB** | 7.40236989319698e-195 | 0.336613455477357 | 0.338 | 0.035 | 9.37214052177669e-191 | 3 | EMB |
| **TXN.1** | 1.39520044479057e-193 | 0.650213058619209 | 0.825 | 0.447 | 1.76646328314934e-189 | 3 | TXN |
| **RAMP2** | 1.89250137642737e-193 | 0.382545517036603 | 0.379 | 0.038 | 2.3960959926947e-189 | 3 | RAMP2 |
| **ANKRD11.2** | 1.05733614073778e-192 | 0.573755880723859 | 0.739 | 0.275 | 1.3386932877881e-188 | 3 | ANKRD11 |
| **GNAI2.1** | 3.72473927121994e-191 | 0.463481937581663 | 0.622 | 0.231 | 4.71589239129157e-187 | 3 | GNAI2 |
| **CDKN1B** | 7.76858752775546e-190 | 0.41457203509298 | 0.483 | 0.114 | 9.83580866889119e-186 | 3 | CDKN1B |
| **MYL6** | 6.81013599394255e-189 | 0.39831157677416 | 0.519 | 0.165 | 8.62231318193066e-185 | 3 | MYL6 |
| **ZNF593** | 8.6294114146229e-188 | 0.447160323269357 | 0.637 | 0.272 | 1.09256977920541e-183 | 3 | ZNF593 |
| **SH3BGRL** | 1.0364110155506e-186 | 0.428402610773022 | 0.501 | 0.136 | 1.31219998678862e-182 | 3 | SH3BGRL |
| **ANXA5** | 1.27565956562805e-186 | 0.380839225531474 | 0.453 | 0.101 | 1.61511257604168e-182 | 3 | ANXA5 |
| **GATD3AL1** | 2.68190926204236e-186 | 0.416400201702725 | 0.522 | 0.154 | 3.39556531667184e-182 | 3 | GATD3AL1 |
| **MECR** | 1.94764379595274e-185 | 0.391839027922314 | 0.482 | 0.108 | 2.46591181005577e-181 | 3 | MECR |
| **ATP6V0C.3** | 2.56166510047315e-184 | 0.64844334916767 | 0.897 | 0.509 | 3.24332418370905e-180 | 3 | ATP6V0C |
| **CYBA** | 4.55185988682969e-183 | 0.475362226175391 | 0.662 | 0.305 | 5.76310980271507e-179 | 3 | CYBA |
| **EEF1B2.1** | 9.56917546016547e-181 | 0.533771983857595 | 0.854 | 0.571 | 1.21155330501155e-176 | 3 | EEF1B2 |
| **ASS1.1** | 1.8388421016316e-180 | 0.6309183180072 | 0.808 | 0.316 | 2.32815798487577e-176 | 3 | ASS1 |
| **ENSGALG00000003348** | 5.40412424931475e-177 | 0.414900631720109 | 0.561 | 0.224 | 6.84216171205741e-173 | 3 | ENSGALG00000003348 |
| **RASSF2.1** | 2.38386247818238e-170 | 0.467069259800471 | 0.663 | 0.278 | 3.01820828362671e-166 | 3 | RASSF2 |
| **ENSGALG00000028749.1** | 1.57549993011262e-169 | 0.497922668178933 | 0.885 | 0.632 | 1.99474046151558e-165 | 3 | ENSGALG00000028749 |
| **TSC22D4.3** | 4.54239678302269e-169 | 0.596861555090628 | 0.713 | 0.222 | 5.75112856698502e-165 | 3 | TSC22D4 |
| **DSTN.1** | 4.55206500133506e-167 | 0.502846456186404 | 0.867 | 0.618 | 5.76336949819031e-163 | 3 | DSTN |
| **PRDX6** | 1.66069013348406e-165 | 0.339518186934765 | 0.439 | 0.108 | 2.10259977800417e-161 | 3 | PRDX6 |
| **BNIP3** | 2.17527878177885e-165 | 0.318490573285545 | 0.38 | 0.073 | 2.75412046561021e-161 | 3 | BNIP3 |
| **PPIA** | 1.12554582841431e-164 | 0.489459145459714 | 0.829 | 0.572 | 1.42505357335536e-160 | 3 | PPIA |
| **IL6R** | 8.67002510009673e-160 | 0.321038780623655 | 0.384 | 0.078 | 1.09771187792325e-155 | 3 | IL6R |
| **COX6A1** | 1.59651555999915e-158 | 0.445395512752424 | 0.731 | 0.392 | 2.02134835051492e-154 | 3 | COX6A1 |
| **KLF6** | 2.40161858145416e-158 | 0.479964400848739 | 0.509 | 0.127 | 3.04068928597911e-154 | 3 | KLF6 |
| **ENSGALG00000051266** | 3.05424351766713e-155 | 0.355153964900762 | 0.47 | 0.165 | 3.86697771771835e-151 | 3 | ENSGALG00000051266 |
| **FKBP1A** | 3.55415773562964e-154 | 0.349907616815997 | 0.436 | 0.137 | 4.49991910908069e-150 | 3 | FKBP1A |
| **ATP5F1D** | 1.01566244065874e-151 | 0.418127749098934 | 0.608 | 0.276 | 1.28593021611803e-147 | 3 | ATP5F1D |
| **FMNL1** | 1.0835878964581e-149 | 0.298084038739727 | 0.349 | 0.075 | 1.37193063570561e-145 | 3 | FMNL1 |
| **TLR7** | 3.53964056771905e-149 | 0.305507121237436 | 0.36 | 0.067 | 4.48153892278909e-145 | 3 | TLR7 |
| **H3F3C** | 1.391628277774e-148 | 0.458891880391663 | 0.8 | 0.467 | 1.76194056248966e-144 | 3 | H3F3C |
| **ENSGALG00000043329** | 9.9890353870555e-148 | 0.463301294831596 | 0.731 | 0.431 | 1.2647117703551e-143 | 3 | ENSGALG00000043329 |
| **H3F3B.1** | 9.34506336848232e-147 | 0.490901404633508 | 0.862 | 0.583 | 1.18317847308355e-142 | 3 | H3F3B |
| **TMEM167A** | 5.52390174289716e-146 | 0.319816261664341 | 0.386 | 0.095 | 6.9938119966821e-142 | 3 | TMEM167A |
| **RHOA** | 6.00819386534542e-146 | 0.431746204600991 | 0.747 | 0.45 | 7.60697425291384e-142 | 3 | RHOA |
| **APLP2** | 2.24740055488664e-143 | 0.383919792719012 | 0.537 | 0.193 | 2.84543384254198e-139 | 3 | APLP2 |
| **ITGB1** | 4.52792176768325e-143 | 0.368804798964153 | 0.478 | 0.162 | 5.73280175006377e-139 | 3 | ITGB1 |
| **COX4I1** | 6.08675838506773e-141 | 0.420339575481864 | 0.666 | 0.343 | 7.70644479133426e-137 | 3 | COX4I1 |
| **FN1.1** | 1.12583366323967e-140 | 1.36278889457205 | 0.951 | 0.103 | 1.42541800102775e-136 | 3 | FN1 |
| **COTL1** | 1.25658244085769e-140 | 0.392598991891863 | 0.573 | 0.24 | 1.59095902836992e-136 | 3 | COTL1 |
| **KLF2** | 2.35258294721897e-139 | 0.327687892660427 | 0.349 | 0.055 | 2.97860526947394e-135 | 3 | KLF2 |
| **ND6.1** | 4.91899464534461e-138 | 0.555534581641992 | 0.983 | 0.857 | 6.22793912047082e-134 | 3 | ND6 |

| | | | | | | | |
|---|---|---|---|---|---|---|---|
| **SEPT7** | 6.44497210091743e-135 | 0.37883352457972 | 0.585 | 0.259 | 8.15997917697155e-131 | 3 | SEPT7 |
| **PGD** | 2.54919055841441e-134 | 0.291674488874189 | 0.351 | 0.077 | 3.22753016600849e-130 | 3 | PGD |
| **SLC38A2** | 2.67537550333107e-134 | 0.300453331857803 | 0.364 | 0.077 | 3.38729292476747e-130 | 3 | SLC38A2 |
| **OSBPL8** | 2.89156999520953e-134 | 0.273645923296093 | 0.323 | 0.063 | 3.66101677093478e-130 | 3 | OSBPL8 |
| **EIF3G** | 1.02951049775171e-133 | 0.306670036964022 | 0.414 | 0.128 | 1.30346324120343e-129 | 3 | EIF3G |
| **ENO1.1** | 1.43402649204984e-133 | 0.422323441784237 | 0.684 | 0.371 | 1.8156209415843e-129 | 3 | ENO1 |
| **EIF3H** | 3.82629313726914e-133 | 0.35187791114583 | 0.499 | 0.195 | 4.84446974109646e-129 | 3 | EIF3H |
| **HK1** | 1.08063694738113e-132 | 0.282512150554305 | 0.362 | 0.093 | 1.36819443907924e-128 | 3 | HK1 |
| **RABGAP1** | 1.76598982895251e-132 | 0.274564661307728 | 0.328 | 0.06 | 2.23591972243678e-128 | 3 | RABGAP1 |
| **ENSGALG00000004509.3** | 4.04115428087265e-132 | 0.52211549679844 | 0.966 | 0.852 | 5.11650543501287e-128 | 3 | ENSGALG00000004509 |
| **LFNG** | 8.51616782220975e-131 | 0.297206408357611 | 0.381 | 0.086 | 1.07823200796998e-126 | 3 | LFNG |
| **DAP** | 9.55356192562156e-131 | 0.276248452617946 | 0.343 | 0.081 | 1.20957647540295e-126 | 3 | DAP |
| **CTSA.1** | 3.916412942591e-129 | 0.450372488088629 | 0.688 | 0.318 | 4.95857042661446e-125 | 3 | CTSA |
| **B2M.1** | 1.94858427922503e-128 | 0.428054214760381 | 0.987 | 0.924 | 2.46710255592681e-124 | 3 | B2M |
| **GINM1** | 2.38381548308267e-127 | 0.264509008080506 | 0.341 | 0.075 | 3.01814878313097e-123 | 3 | GINM1 |
| **CTSZ.1** | 7.83405178298125e-127 | 0.451445724445633 | 0.686 | 0.338 | 9.91869296243255e-123 | 3 | CTSZ |
| **ARHGDIB** | 5.99136677804004e-126 | 0.31101745171732 | 0.419 | 0.128 | 7.58566947767649e-122 | 3 | ARHGDIB |
| **MARCO.3** | 9.88325770757103e-126 | -0.893782380830593 | 0.057 | 0.424 | 1.25131925835557e-121 | 3 | MARCO |
| **SHFM1** | 6.90054041666836e-125 | 0.335599217410895 | 0.482 | 0.183 | 8.73677422154381e-121 | 3 | SHFM1 |
| **RCSD1** | 7.56366968651091e-125 | 0.351848457235498 | 0.461 | 0.135 | 9.57636219009146e-121 | 3 | RCSD1 |
| **EIF3M** | 1.35706542294875e-124 | 0.339450968212775 | 0.476 | 0.173 | 1.71818053199541e-120 | 3 | EIF3M |
| **ARF6** | 1.65292614413601e-124 | 0.36032036561696 | 0.552 | 0.217 | 2.09276979109061e-120 | 3 | ARF6 |
| **BTG2.3** | 5.05039402388591e-122 | 0.581420476560749 | 0.718 | 0.26 | 6.39430387364195e-118 | 3 | BTG2 |
| **NFE2L2** | 2.57686712586507e-121 | 0.316896453189122 | 0.43 | 0.122 | 3.26257146805776e-117 | 3 | NFE2L2 |
| **GNG5.1** | 5.54916726071325e-121 | 0.412920787189424 | 0.756 | 0.456 | 7.02580066878905e-117 | 3 | GNG5 |
| **PPIB.1** | 7.42866739417685e-120 | 0.399345164623386 | 0.639 | 0.323 | 9.40543578776731e-116 | 3 | PPIB |
| **TPI1** | 3.46476676064039e-119 | 0.311091638445348 | 0.453 | 0.158 | 4.38674119564679e-115 | 3 | TPI1 |
| **YWHAQ** | 3.56712399487537e-119 | 0.34719344169075 | 0.539 | 0.224 | 4.5163356899117e-115 | 3 | YWHAQ |
| **EIF4A2** | 5.66901047146256e-119 | 0.390192480377634 | 0.765 | 0.529 | 7.17753415791875e-115 | 3 | EIF4A2 |
| **PRNP** | 1.22581576788497e-118 | 0.317391698252103 | 0.417 | 0.108 | 1.55200534371916e-114 | 3 | PRNP |
| **ENSGALG00000006899** | 1.9075549827949e-118 | 0.345071803125576 | 0.574 | 0.291 | 2.41515536371663e-114 | 3 | ENSGALG00000006899 |
| **PGLS** | 6.66971872783304e-118 | 0.285212595060113 | 0.4 | 0.121 | 8.44453088130941e-114 | 3 | PGLS |
| **AP1S2** | 7.74595187156195e-116 | 0.339710627047859 | 0.486 | 0.182 | 9.80714966458458e-112 | 3 | AP1S2 |
| **DDX5.3** | 9.67004949748492e-116 | 0.514318119247114 | 0.853 | 0.473 | 1.22432496687657e-111 | 3 | DDX5 |
| **TGM4.3** | 1.09814038241823e-114 | -0.825190722549306 | 0.067 | 0.374 | 1.39035553817972e-110 | 3 | TGM4 |
| **MYH9.1** | 2.77759073285424e-114 | 0.415391781973213 | 0.709 | 0.417 | 3.51670762686675e-110 | 3 | MYH9 |
| **JUN.3** | 3.15642291935624e-113 | 0.647708583889747 | 0.793 | 0.252 | 3.99634705819693e-109 | 3 | JUN |
| **ATP6V1E1** | 1.01744281866786e-112 | 0.263448829956762 | 0.342 | 0.088 | 1.28818435271538e-108 | 3 | ATP6V1E1 |
| **BZW2** | 1.17276413665762e-112 | 0.284780948215957 | 0.368 | 0.099 | 1.48483667342222e-108 | 3 | BZW2 |
| **PKLR** | 1.17981065316966e-112 | 0.269135217380132 | 0.376 | 0.12 | 1.49375826797811e-108 | 3 | PKLR |
| **HIF1A** | 3.14931040504179e-112 | 0.258373718231898 | 0.33 | 0.086 | 3.98734190382342e-108 | 3 | HIF1A |
| **EIF3L** | 8.37874440301525e-111 | 0.272384346244587 | 0.377 | 0.121 | 1.06083278165761e-106 | 3 | EIF3L |
| **EIF3E** | 1.91796355549321e-110 | 0.332996358436095 | 0.513 | 0.231 | 2.42833365760995e-106 | 3 | EIF3E |
| **CTSD.2** | 4.71024533711083e-110 | -0.753708303785423 | 0.248 | 0.521 | 5.96364162131603e-106 | 3 | CTSD |
| **ENSGALG00000026970.2** | 7.44426771958589e-110 | 0.548631812716359 | 0.849 | 0.494 | 9.42518735976 77e-106 | 3 | ENSGALG00000026970 |
| **LTA4H** | 8.16293063615008e-110 | 0.362594259786304 | 0.418 | 0.079 | 1.03350864784296e-105 | 3 | LTA4H |
| **TLN1** | 1.54741369274387e-109 | 0.333266373288552 | 0.505 | 0.222 | 1.95918047638302e-105 | 3 | TLN1 |
| **TLR1B** | 5.68732349581935e-109 | 0.282387121491909 | 0.356 | 0.089 | 7.20072027805688e-105 | 3 | TLR1B |
| **DMB2** | 2.18171115082805e-108 | 0.305351714181642 | 0.459 | 0.181 | 2.7622644880634e-104 | 3 | DMB2 |
| **DUSP1.1** | 2.41910253852169e-108 | 0.511908985694606 | 0.562 | 0.144 | 3.06282572402231e-104 | 3 | DUSP1 |
| **BLB2.2** | 3.11382595284939e-108 | 0.468979572326404 | 0.982 | 0.847 | 3.94241503890261e-104 | 3 | BLB2 |
| **NDUFB2** | 3.95598205014483e-108 | 0.332834438312184 | 0.584 | 0.302 | 5.00866887368837e-104 | 3 | NDUFB2 |
| **ANXA11** | 1.59458778345913e-106 | 0.271675150625412 | 0.367 | 0.112 | 2.01890759263761e-102 | 3 | ANXA11 |
| **NPM1** | 6.5897659337302e-106 | 0.352030108087689 | 0.549 | 0.276 | 8.34330264869581e-102 | 3 | NPM1 |

| | | | | | | | |
|---|---|---|---|---|---|---|---|
| **ALCAM** | 1.92229474521761e-105 | 0.269105142715074 | 0.316 | 0.075 | 2.43381737692002e-101 | 3 | ALCAM |
| **ACTB.1** | 4.37572366888165e-105 | 0.409801944492822 | 0.994 | 0.978 | 5.54010373717106e-101 | 3 | ACTB |
| **CYTB.1** | 4.87474110920849e-105 | 0.45754329563941 | 0.972 | 0.869 | 6.17190971836887e-101 | 3 | CYTB |
| **TNFAIP8** | 5.46577412894479e-105 | 0.285813391719668 | 0.421 | 0.144 | 6.920216624657e-101 | 3 | TNFAIP8 |
| **AIF1L** | 1.01814766129889e-104 | 0.371003926923025 | 0.685 | 0.415 | 1.28907675397052e-100 | 3 | AIF1L |
| **ENSGALG00000030086** | 2.98115506364236e-104 | 0.289291730246569 | 0.459 | 0.189 | 3.7744404260776e-100 | 3 | ENSGALG00000030086 |
| **SPEN** | 1.17028736505348e-103 | 0.346321895254944 | 0.504 | 0.2 | 1.48170083289421e-99 | 3 | SPEN |
| **TOP2B** | 2.66447900890844e-103 | 0.289024400685785 | 0.429 | 0.146 | 3.37349687317898e-99 | 3 | TOP2B |
| **ZEB2.3** | 2.69367234044944e-102 | 0.467428138287928 | 0.663 | 0.261 | 3.41045855024303e-98 | 3 | ZEB2 |
| **EIF3D** | 2.52358382120706e-101 | 0.277401811837839 | 0.392 | 0.141 | 3.19510947603026e-97 | 3 | EIF3D |
| **RAB1B.1** | 3.04336295558481e-100 | 0.340681038860375 | 0.597 | 0.298 | 3.85320183806593e-96 | 3 | RAB1B |
| **PLXNC1** | 2.89239260525181e-99 | 0.321025583683837 | 0.49 | 0.199 | 3.66205827750931e-95 | 3 | PLXNC1 |
| **SPCS3** | 5.56931538953386e-99 | 0.300109097990505 | 0.459 | 0.198 | 7.05131021468882e-95 | 3 | SPCS3 |
| **ENSGALG00000047532.2** | 2.59886927142156e-98 | 0.35507117075908 | 0.639 | 0.324 | 3.29042838454683e-94 | 3 | ENSGALG00000047532 |
| **HHEX** | 4.77728064929553e-97 | 0.273525804837767 | 0.374 | 0.121 | 6.04851503007307e-93 | 3 | HHEX |
| **CG-1B** | 8.60031031408285e-94 | 0.503496301956193 | 0.547 | 0.126 | 1.08888528886603e-89 | 3 | CG-1B |
| **MSN** | 9.59397941017055e-94 | 0.34619975092182 | 0.565 | 0.3 | 1.21469373312169e-89 | 3 | MSN |
| **SERP1** | 7.2297414341197e-93 | 0.308790241147696 | 0.485 | 0.231 | 9.15357562973895e-89 | 3 | SERP1 |
| **NDUFS8** | 3.91946883754624e-92 | 0.263475231438929 | 0.387 | 0.14 | 4.96243949521729e-88 | 3 | NDUFS8 |
| **EIF5A2** | 2.17148444730749e-91 | 0.270470323201619 | 0.4 | 0.147 | 2.74931645873601e-87 | 3 | EIF5A2 |
| **MMR1L4** | 2.49296227148212e-91 | 0.257713040819226 | 0.321 | 0.083 | 3.15633953192352e-87 | 3 | MMR1L4 |
| **NRK** | 3.79283115637974e-91 | 0.286919900138259 | 0.435 | 0.177 | 4.80210352709239e-87 | 3 | NRK |
| **SERBP1** | 9.9394697776933e-91 | 0.308101147521531 | 0.546 | 0.284 | 1.25843626855375e-86 | 3 | SERBP1 |
| **ADAM10** | 2.75789672779811e-90 | 0.286090345046849 | 0.459 | 0.198 | 3.49177304706519e-86 | 3 | ADAM10 |
| **COX7C** | 1.03351804722287e-88 | 0.319295934255426 | 0.614 | 0.376 | 1.30853719958887e-84 | 3 | COX7C |
| **ZNF706** | 1.10604343307034e-88 | 0.307843976692352 | 0.559 | 0.283 | 1.40036159061036e-84 | 3 | ZNF706 |
| **ENSGALG00000029330** | 3.65039918932965e-88 | 0.26331341777305 | 0.43 | 0.186 | 4.62177041361027e-84 | 3 | ENSGALG00000029330 |
| **ARPC4.1** | 1.69226671941203e-87 | 0.343484393444677 | 0.731 | 0.465 | 2.14257889344757e-83 | 3 | ARPC4 |
| **DBI** | 2.446235693337e-87 | 0.288485600555371 | 0.434 | 0.18 | 3.09717901133397e-83 | 3 | DBI |
| **CALM1** | 1.35307766220787e-85 | 0.346642650238231 | 0.742 | 0.493 | 1.71313162812139e-81 | 3 | CALM1 |
| **NCOR2** | 3.57609721101091e-85 | 0.286112772537245 | 0.465 | 0.197 | 4.52769667886092e-81 | 3 | NCOR2 |
| **PRDX1.1** | 6.83018675916492e-85 | 0.344328243554438 | 0.642 | 0.361 | 8.6476994557787e-81 | 3 | PRDX1 |
| **MTPN** | 8.37880054706619e-85 | 0.285758261072493 | 0.492 | 0.236 | 1.06083993726405e-80 | 3 | MTPN |
| **IFI30.2** | 1.59791176452988e-84 | 0.534103917172067 | 0.994 | 0.802 | 2.02311608507128e-80 | 3 | IFI30 |
| **NSA2** | 4.67992856233429e-84 | 0.279176845861661 | 0.492 | 0.245 | 5.92525755277145e-80 | 3 | NSA2 |
| **PSAP.1** | 1.7040984532713e-83 | 0.362550968047663 | 0.629 | 0.337 | 2.15755905168679e-79 | 3 | PSAP |
| **ENSGALG00000011190.3** | 5.11394437158044e-83 | -0.595404655111084 | 0.185 | 0.421 | 6.47476496885799e-79 | 3 | ENSGALG00000011190 |
| **ENSGALG00000047864** | 8.66460651686273e-82 | 0.269720250198924 | 0.412 | 0.168 | 1.09702583109999e-77 | 3 | ENSGALG00000047864 |
| **SLC40A1.3** | 1.22193635885277e-81 | -0.477962747407415 | 0.175 | 0.457 | 1.54709362394349e-77 | 3 | SLC40A1 |
| **CDV3** | 1.69230289075934e-81 | 0.264789534813874 | 0.394 | 0.153 | 2.1426246899904e-77 | 3 | CDV3 |
| **TMED10** | 4.9410075547803e-81 | 0.282480779297575 | 0.465 | 0.217 | 6.25580966510734e-77 | 3 | TMED10 |
| **COX5A** | 5.68453878073285e-81 | 0.30055416881392 | 0.543 | 0.3 | 7.19719455028586e-77 | 3 | COX5A |
| **CST3.2** | 9.25060049426248e-81 | 0.396395354699211 | 0.89 | 0.686 | 1.17121852857857e-76 | 3 | CST3 |
| **CLIC2** | 3.00429249317119e-80 | 0.253025276304257 | 0.41 | 0.173 | 3.80373472560404e-76 | 3 | CLIC2 |
| **ENSGALG00000038672** | 1.21576883458246e-79 | 0.280890746313813 | 0.498 | 0.268 | 1.53928492146486e-75 | 3 | ENSGALG00000038672 |
| **MYCBP2** | 4.28233527717466e-78 | 0.252369225674089 | 0.386 | 0.15 | 5.42186469443084e-74 | 3 | MYCBP2 |
| **HNRNPA3** | 8.28468913586792e-78 | 0.300851405694135 | 0.545 | 0.288 | 1.04892449149224e-73 | 3 | HNRNPA3 |
| **YWHAZ** | 1.15125153118869e-77 | 0.250171296891561 | 0.398 | 0.163 | 1.457599563638e-73 | 3 | YWHAZ |
| **PRCP** | 2.10846631163554e-77 | 0.268417541527559 | 0.364 | 0.108 | 2.66952919716175e-73 | 3 | PRCP |
| **MCL1.1** | 1.1129088291383e-76 | 0.306282283446955 | 0.484 | 0.207 | 1.409053868572e-72 | 3 | MCL1 |
| **C15orf48** | 1.95294642949408e-75 | 0.257772937198507 | 0.35 | 0.1 | 2.47262547438245e-71 | 3 | C15orf48 |
| **FOS.3** | 4.26032309688983e-75 | 0.591658994428717 | 0.812 | 0.335 | 5.39399507297222e-71 | 3 | FOS |
| **PTBP1** | 4.14549154519824e-74 | 0.263788735091429 | 0.44 | 0.189 | 5.24860684537549e-70 | 3 | PTBP1 |
| **BF2** | 8.42143849625337e-74 | 0.274735564418087 | 0.453 | 0.221 | 1.06623832801064e-69 | 3 | BF2 |

| | | | | | | | |
|---|---|---|---|---|---|---|---|
| **CCL4** | 2.10314973700369e-73 | 0.536507857769632 | 0.503 | 0.113 | 2.66279788202037e-69 | 3 | CCL4 |
| **PRRC2C** | 2.24318619443011e-73 | 0.256137702259048 | 0.432 | 0.181 | 2.84009804076796e-69 | 3 | PRRC2C |
| **DYL1** | 7.49818973007563e-73 | 0.259106092975216 | 0.425 | 0.196 | 9.49345801724875e-69 | 3 | DYL1 |
| **BRI3.1** | 2.1536623589896e-72 | 0.307892867645947 | 0.605 | 0.345 | 2.72675191271674e-68 | 3 | BRI3 |
| **SMAP2.1** | 3.14169449505405e-70 | 0.297052911366947 | 0.585 | 0.331 | 3.97769940018793e-66 | 3 | SMAP2 |
| **EXFABP.3** | 1.85833241547107e-69 | 1.56014089085201 | 0.912 | 0.141 | 2.35283467122793e-65 | 3 | EXFABP |
| **MYO1F** | 2.79417069606301e-66 | 0.251699304195578 | 0.433 | 0.199 | 3.53769951828538e-62 | 3 | MYO1F |
| **GRN.2** | 7.51340736209731e-66 | 0.379447576987957 | 0.848 | 0.598 | 9.5127250611514e-62 | 3 | GRN |
| **S100A12** | 3.77446715731077e-65 | 0.952798882747179 | 0.382 | 0.023 | 4.77885286787116e-61 | 3 | S100A12 |
| **EIF4G2** | 1.08880729591544e-64 | 0.267277112784959 | 0.583 | 0.337 | 1.37853891735854e-60 | 3 | EIF4G2 |
| **BLB1.2** | 1.14826313307634e-64 | 0.409878601345032 | 0.98 | 0.844 | 1.45381595278795e-60 | 3 | BLB1 |
| **PNRC1.3** | 1.34940716179585e-64 | 0.388974684040221 | 0.818 | 0.506 | 1.70848440754973e-60 | 3 | PNRC1 |
| **SPI1.1** | 5.5744166789316e-63 | 0.277366988644196 | 0.543 | 0.281 | 7.0577689571953e-59 | 3 | SPI1 |
| **COX7B** | 1.48989737152018e-62 | 0.267763901189009 | 0.551 | 0.327 | 1.8863590620817e-58 | 3 | COX7B |
| **NFKBIA.3** | 7.58742603175373e-62 | 0.369827823242207 | 0.596 | 0.3 | 9.60644009880339e-58 | 3 | NFKBIA |
| **ATF4** | 2.79470374426684e-61 | 0.259260521149413 | 0.497 | 0.246 | 3.53837441061624e-57 | 3 | ATF4 |
| **HNRNPH2** | 2.7636348138455e-60 | 0.269432389628562 | 0.562 | 0.327 | 3.49903803780979e-56 | 3 | HNRNPH2 |
| **HSPA5.3** | 5.0080762794803e-60 | 0.477006054489224 | 0.7 | 0.36 | 6.34072537745001e-56 | 3 | HSPA5 |
| **EIF1** | 5.05787620867367e-60 | 0.262871825114513 | 0.543 | 0.328 | 6.40377706780173e-56 | 3 | EIF1 |
| **LAPTM5.1** | 1.14749592657843e-59 | 0.28416227245424 | 0.709 | 0.457 | 1.45284459264095e-55 | 3 | LAPTM5 |
| **TGFBI** | 1.23135635503371e-59 | 0.307571901910107 | 0.567 | 0.285 | 1.55902028110818e-55 | 3 | TGFBI |
| **RNASET2** | 2.28747317528853e-57 | 0.253705254510514 | 0.468 | 0.242 | 2.89616978723281e-53 | 3 | RNASET2 |
| **ARPC1B** | 1.58400782280933e-56 | 0.266421200195626 | 0.57 | 0.361 | 2.00551230445889e-52 | 3 | ARPC1B |
| **ENSGALG00000052354.1** | 4.32265128270224e-56 | 0.254869575557191 | 0.453 | 0.206 | 5.47290878902931e-52 | 3 | ENSGALG00000052354 |
| **CTSC.1** | 1.7542274927154e-55 | 0.275843327900903 | 0.566 | 0.311 | 2.22102742852697e-51 | 3 | CTSC |
| **MT-ND2** | 9.63828343569212e-55 | 0.307031996497316 | 0.776 | 0.583 | 1.22030306579298e-50 | 3 | MT-ND2 |
| **GNS 0.10** | 1.35165476394151e-53 | 0.255897741316172 | 0.541 | 0.315 | 1.71133009662635e-49 | 3 | GNS |
| **YBX1.1** | 4.47409608992744e-53 | 0.273945796444109 | 0.675 | 0.495 | 5.66465305945714e-49 | 3 | YBX1 |
| **PTPRC.1** | 8.40011524186649e-52 | 0.278678121592075 | 0.593 | 0.349 | 1.06353859077272e-47 | 3 | PTPRC |
| **SGK1.1** | 2.90435763460122e-50 | 0.25862957311946 | 0.407 | 0.149 | 3.67720720116861e-46 | 3 | SGK1 |
| **ENSGALG00000005257.1** | 2.94722492654336e-49 | -0.312085229000055 | 0.113 | 0.348 | 3.73148147949655e-45 | 3 | ENSGALG00000005257 |
| **HSPA8.3** | 9.39514856961854e-49 | 0.391952731971632 | 0.889 | 0.586 | 1.1895197603994e-44 | 3 | HSPA8 |
| **CSF1R.1** | 1.07791627988809e-47 | 0.272984501005777 | 0.631 | 0.383 | 1.36474980196631e-43 | 3 | CSF1R |
| **LCP1.1** | 1.4157985807524e-46 | 0.262724428994866 | 0.733 | 0.521 | 1.79254258309061e-42 | 3 | LCP1 |
| **CD47.3** | 1.05628595495337e-45 | 0.277055718896373 | 0.597 | 0.328 | 1.33736364756646e-41 | 3 | CD47 |
| **CTSV.1** | 2.4655538732811e-44 | 0.269430199841946 | 0.528 | 0.252 | 3.1216377589612e-40 | 3 | CTSV |
| **FTH1.2** | 4.10585223273163e-43 | 0.342233827285242 | 1 | 0.99 | 5.19841951186152e-39 | 3 | FTH1 |
| **HSP90AA1.3** | 2.64814010223662e-42 | 0.3931002825058 | 0.965 | 0.763 | 3.35281018344179e-38 | 3 | HSP90AA1 |
| **ENSGALG00000014585.2** | 1.37925474866013e-41 | -0.334867003834506 | 0.256 | 0.4 | 1.74627443727859e-37 | 3 | ENSGALG00000014585 |
| **HTRA1.3** | 2.75742569375181e-41 | -0.368266386570258 | 0.014 | 0.255 | 3.49117667085916e-37 | 3 | HTRA1 |
| **C1QA.2** | 7.37544414436808e-41 | -0.28983760105563 | 0.308 | 0.488 | 9.33804983118442e-37 | 3 | C1QA |
| **C1QB.2** | 5.43188749685663e-40 | -0.318064068861567 | 0.305 | 0.455 | 6.87731275977018e-36 | 3 | C1QB |
| **HSP90B1.3** | 5.97296083920189e-39 | 0.293611888177329 | 0.643 | 0.412 | 7.56236571851351e-35 | 3 | HSP90B1 |
| **AVD** | 6.55656519876287e-35 | 0.393768491604742 | 0.388 | 0.046 | 8.30126719815367e-31 | 3 | AVD |
| **HMOX1.2** | 2.99111417350476e-30 | -0.336396328684109 | 0.312 | 0.364 | 3.78704965507438e-26 | 3 | HMOX1 |
| **MAFB.3** | 3.17602993495074e-23 | -0.258700369155519 | 0.571 | 0.625 | 4.02117150064113e-19 | 3 | MAFB |
| **ENSGALG00000030007.3** | 0 | 1.22638216383825 | 0.942 | 0.148 | 0 | 4 | ENSGALG00000030007 |
| **HTRA1.4** | 0 | 0.893003090067906 | 0.815 | 0.169 | 0 | 4 | HTRA1 |
| **SH3PXD2A** | 0 | 0.586243408498591 | 0.557 | 0.061 | 0 | 4 | SH3PXD2A |
| **VSIG4** | 0 | 0.569876984725429 | 0.549 | 0.058 | 0 | 4 | VSIG4 |
| **IFI30.3** | 0 | -1.30740254204876 | 0.488 | 0.858 | 0 | 4 | IFI30 |
| **FTH1.3** | 0 | -1.32309770270743 | 0.957 | 0.994 | 0 | 4 | FTH1 |
| **MARCO.4** | 1.49058658556194e-282 | 1.52539693313143 | 0.983 | 0.322 | 1.88723167597997e-278 | 4 | MARCO |
| **ADGRG2** | 1.68445626125053e-256 | 0.421022572649987 | 0.414 | 0.026 | 2.1326900723693e-252 | 4 | ADGRG2 |

| | | | | | | | |
|---|---|---|---|---|---|---|---|
| **ENSGALG00000049402** | 1.73944431364429e-223 | 0.481046090908664 | 0.524 | 0.106 | 2.20231044550503e-219 | 4 | ENSGALG00000049402 |
| **SEMA3D** | 1.92117861681121e-212 | 0.343845934607863 | 0.361 | 0.025 | 2.43240424674467e-208 | 4 | SEMA3D |
| **TNFAIP2.1** | 3.64708834757612e-175 | 0.616431674848304 | 0.643 | 0.133 | 4.61757855686612e-171 | 4 | TNFAIP2 |
| **ENSGALG00000005257.[illegible]** | 1.51029924959746e-172 | 0.64415029453489 | 0.691 | 0.284 | 1.91218987991535e-168 | 4 | ENSGALG00000005257 |
| **ZDHHC23** | 6.82262560422204e-170 | 0.280777069372355 | 0.296 | 0.023 | 8.63812627750553e-166 | 4 | ZDHHC23 |
| **TNFSF13B.1** | 7.46589868830277e-168 | 0.513236637048667 | 0.558 | 0.118 | 9.45257432926014e-164 | 4 | TNFSF13B |
| **FAM49A.2** | 3.49292333024695e-167 | 0.531476280795825 | 0.715 | 0.376 | 4.42239022842567e-163 | 4 | FAM49A |
| **IL8L1** | 8.91945430891342e-162 | 0.671158393050492 | 0.364 | 0.026 | 1.12929211005153e-157 | 4 | IL8L1 |
| **ITPR1** | 1.35478724270753e-161 | 0.275989081769856 | 0.289 | 0.026 | 1.715296127992e-157 | 4 | ITPR1 |
| **CD82** | 3.72904043426943e-160 | 0.342045945022902 | 0.403 | 0.084 | 4.72133809382853e-156 | 4 | CD82 |
| **IGFBP7** | 6.66727109122645e-158 | 0.361108639303647 | 0.402 | 0.058 | 8.44143192860181e-154 | 4 | IGFBP7 |
| **CTSS.2** | 2.04843030291632e-147 | -0.84786441617154 | 0.219 | 0.698 | 2.59351760652235e-143 | 4 | CTSS |
| **ENSGALG00000046843** | 5.50879888084878e-147 | 0.311802435404556 | 0.367 | 0.075 | 6.97469026304264e-143 | 4 | ENSGALG00000046843 |
| **ENSGALG00000048767** | 6.79188237508383e-146 | 0.402153692905858 | 0.522 | 0.171 | 8.59920227509364e-142 | 4 | ENSGALG00000048767 |
| **FYB.1** | 1.48875069500304e-143 | 0.481149445848992 | 0.653 | 0.281 | 1.88490725494335e-139 | 4 | FYB |
| **TMSB15B.3** | 3.49439193002612e-130 | -0.784494907124525 | 0.382 | 0.742 | 4.42424962260607e-126 | 4 | TMSB15B |
| **KTN1** | 3.87962716588506e-126 | 0.40345082557937 | 0.514 | 0.177 | 4.91199595472708e-122 | 4 | KTN1 |
| **CCDC88C** | 6.3185423577106e-125 | 0.378196381263299 | 0.443 | 0.126 | 7.99990647909738e-121 | 4 | CCDC88C |
| **GRP.1** | 4.12389586367662e-120 | 0.580819530530494 | 0.603 | 0.125 | 5.22126455300096e-116 | 4 | GRP |
| **TTYH3** | 5.0814349200665e-120 | 0.327202133682754 | 0.411 | 0.121 | 6.43360475229619e-116 | 4 | TTYH3 |
| **ITGAV** | 7.9806191139275e-118 | 0.364794246748396 | 0.493 | 0.187 | 1.01042618601436e-113 | 4 | ITGAV |
| **DNASE1L3** | 1.21405781916534e-116 | 0.381540023601747 | 0.477 | 0.13 | 1.53711860484524e-112 | 4 | DNASE1L3 |
| **CSF1R.2** | 3.97680757567641e-116 | 0.469968027297827 | 0.754 | 0.38 | 5.03503607156391e-112 | 4 | CSF1R |
| **MTMR10** | 1.80228508743631e-115 | 0.279753498337586 | 0.319 | 0.072 | 2.28187314920311e-111 | 4 | MTMR10 |
| **CXCL13.4** | 5.83733951599087e-115 | 1.68423237579033 | 0.999 | 0.366 | 7.39065556119604e-111 | 4 | CXCL13 |
| **MCF2L** | 7.15779047192228e-114 | 0.311350215807153 | 0.372 | 0.089 | 9.06247851650079e-110 | 4 | MCF2L |
| **CST7** | 8.39820119894381e-113 | 0.420426898311475 | 0.52 | 0.146 | 1.06329625379828e-108 | 4 | CST7 |
| **C1QC.2** | 1.13557047004156e-108 | -0.825375091596798 | 0.081 | 0.592 | 1.43774577211961e-104 | 4 | C1QC |
| **GRN.3** | 5.70048644593837e-108 | -0.652688049724949 | 0.279 | 0.662 | 7.21738588920257e-104 | 4 | GRN |
| **SAT1.3** | 1.8089752220591e-107 | -0.725894196626987 | 0.187 | 0.592 | 2.29034352864902e-103 | 4 | SAT1 |
| **ACTN1** | 1.62383791888615e-97 | 0.255253254775216 | 0.325 | 0.077 | 2.05594118910175e-93 | 4 | ACTN1 |
| **TGM4.4** | 1.64496472128105e-97 | 1.12635658207438 | 0.963 | 0.276 | 2.08268983361394e-93 | 4 | TGM4 |
| **CCPG1** | 2.6552386113878e-96 | 0.338751620539966 | 0.459 | 0.163 | 3.36179760587809e-92 | 4 | CCPG1 |
| **WIPF1** | 2.8312012078578e-93 | 0.333019698527384 | 0.477 | 0.173 | 3.58458384926876e-89 | 4 | WIPF1 |
| **PTPRC.2** | 1.40769624775448e-92 | 0.41868365274561 | 0.683 | 0.349 | 1.78228421928195e-88 | 4 | PTPRC |
| **SPIC.2** | 2.21849611709175e-92 | 0.565932459047328 | 0.644 | 0.162 | 2.80883793384987e-88 | 4 | SPIC |
| **BLB1.3** | 1.56812789312658e-91 | -0.597083566006762 | 0.749 | 0.871 | 1.98540672548757e-87 | 4 | BLB1 |
| **NR1H3** | 1.4648952945988e-84 | 0.274075875279264 | 0.369 | 0.13 | 1.85470393249155e-80 | 4 | NR1H3 |
| **EXFABP.4** | 3.58605024989081e-84 | -1.34013785614493 | 0.043 | 0.253 | 4.54029822138675e-80 | 4 | EXFABP |
| **FGD4** | 6.31992578139044e-84 | 0.284016384648322 | 0.343 | 0.104 | 8.00165803181844e-80 | 4 | FGD4 |
| **PTGS2.1** | 3.64346180141994e-82 | 0.486432437732333 | 0.419 | 0.091 | 4.61298698677778e-78 | 4 | PTGS2 |
| **BAG5** | 3.76291284382764e-79 | 0.292476940641816 | 0.398 | 0.141 | 4.76422395157018e-75 | 4 | BAG5 |
| **IL8** | 7.15090530058224e-79 | 0.330018538573141 | 0.298 | 0.043 | 9.05376120106717e-75 | 4 | IL8 |
| **C1QB.3** | 4.83622214342668e-75 | -0.605620114112965 | 0.066 | 0.472 | 6.12314085579251e-71 | 4 | C1QB |
| **RPS26.4** | 3.37962191012496e-74 | -0.567388457321642 | 0.501 | 0.731 | 4.27893930040922e-70 | 4 | RPS26 |
| **S100A6.4** | 3.44612330363177e-72 | -0.856309954974225 | 0.045 | 0.311 | 4.36313671472818e-68 | 4 | S100A6 |
| **RASGEF1A** | 4.54067863110516e-72 | 0.29380531110782 | 0.372 | 0.114 | 5.74895321484225e-68 | 4 | RASGEF1A |
| **RPS11.4** | 6.04757420765331e-72 | -0.626822586991034 | 0.698 | 0.794 | 7.65683370430986e-68 | 4 | RPS11 |
| **SLC40A1.4** | 1.14568662297549e-71 | -0.624112742027883 | 0.047 | 0.459 | 1.45055383334927e-67 | 4 | SLC40A1 |
| **PPDPF.4** | 1.49402881670942e-70 | -0.862036931729026 | 0.032 | 0.393 | 1.89158988483579e-66 | 4 | PPDPF |
| **HMOX1.3** | 8.06156659986509e-69 | -0.622194098914531 | 0.051 | 0.387 | 1.02067494720892e-64 | 4 | HMOX1 |
| **ENSGALG00000014585.[illegible]** | 6.14265331036407e-66 | -0.552864307870765 | 0.066 | 0.413 | 7.77721335625195e-62 | 4 | ENSGALG00000014585 |
| **ACTR3.1** | 4.90884432413332e-65 | 0.329295360036155 | 0.625 | 0.362 | 6.21508779878519e-61 | 4 | ACTR3 |
| **TPT1.4** | 1.02261771911046e-63 | -0.544983208729439 | 0.696 | 0.81 | 1.29473629416575e-59 | 4 | TPT1 |

| | | | | | | | |
|---|---|---|---|---|---|---|---|
| **ASS1.2** | 1.46544892256681e-61 | -0.492145084826516 | 0.072 | 0.405 | 1.85540488086184e-57 | 4 | ASS1 |
| **TXN.2** | 2.87995371794647e-61 | -0.466826390841126 | 0.247 | 0.516 | 3.64630940229202e-57 | 4 | TXN |
| **RPS19.4** | 3.02075217920384e-61 | -0.526847998015421 | 0.767 | 0.857 | 3.82457433408998e-57 | 4 | RPS19 |
| **RPLP1.4** | 5.34593515745259e-61 | -0.492569838342315 | 0.84 | 0.912 | 6.76848850285072e-57 | 4 | RPLP1 |
| **TPP1.2** | 7.07940680896741e-61 | -0.431075685135592 | 0.116 | 0.454 | 8.96323696083364e-57 | 4 | TPP1 |
| **NADK** | 1.88358056698935e-60 | 0.292773445606956 | 0.499 | 0.251 | 2.38480135586522e-56 | 4 | NADK |
| **ENSGALG00000037441.4** | 1.59648312987287e-58 | -0.507338672229725 | 0.671 | 0.789 | 2.02130729073204e-54 | 4 | ENSGALG00000037441 |
| **CTSB.2** | 1.10596877755283e-54 | -0.466976010907241 | 0.049 | 0.41 | 1.40026706925964e-50 | 4 | CTSB |
| **LSP1P1.2** | 3.51423980803663e-53 | -0.410702192906864 | 0.117 | 0.394 | 4.44937902095517e-49 | 4 | LSP1P1 |
| **CTSA.2** | 7.55794473751851e-53 | -0.401649367333287 | 0.08 | 0.39 | 9.56911383217218e-49 | 4 | CTSA |
| **RPL23.4** | 2.01498462190823e-51 | -0.456585514751297 | 0.695 | 0.807 | 2.551172029798e-47 | 4 | RPL23 |
| **RPS15.4** | 4.10742306232609e-51 | -0.470098549660153 | 0.668 | 0.781 | 5.20040833921106e-47 | 4 | RPS15 |
| **CCNI.1** | 4.57422003902023e-49 | 0.275760317666454 | 0.474 | 0.245 | 5.79141999140351e-45 | 4 | CCNI |
| **PNRC1.4** | 1.61953498524469e-48 | -0.406700630859916 | 0.356 | 0.561 | 2.0504932448183e-44 | 4 | PNRC1 |
| **S100A10.4** | 2.14770623546135e-47 | -0.569845076952471 | 0.281 | 0.396 | 2.71921086471762e-43 | 4 | S100A10 |
| **RPL15.4** | 1.95612101329864e-46 | -0.426113073578336 | 0.664 | 0.782 | 2.47664481493741e-42 | 4 | RPL15 |
| **BLB2.3** | 2.45661905828428e-45 | -0.378799192677427 | 0.819 | 0.868 | 3.11032538969373e-41 | 4 | BLB2 |
| **RPL23A.3** | 2.78524440981087e-45 | -0.413594851083486 | 0.476 | 0.626 | 3.52639794726154e-41 | 4 | RPL23A |
| **GNB2L1.4** | 1.0964661906752e-44 | -0.431945919469432 | 0.514 | 0.641 | 1.38823584401387e-40 | 4 | GNB2L1 |
| **ATP6V0C.4** | 1.70846424244798e-44 | -0.399117699924881 | 0.392 | 0.571 | 2.16308657736339e-40 | 4 | ATP6V0C |
| **LAMP1.1** | 1.04818947291057e-43 | -0.32637137878722 | 0.092 | 0.365 | 1.32711269165207e-39 | 4 | LAMP1 |
| **RPS17.4** | 4.25114367995402e-43 | -0.413491475883105 | 0.739 | 0.833 | 5.38237301318978e-39 | 4 | RPS17 |
| **RPL21.4** | 5.62012348832602e-43 | -0.402954147001557 | 0.749 | 0.836 | 7.11563834856957e-39 | 4 | RPL21 |
| **RPLP2.4** | 6.54338424636614e-43 | -0.447083963149735 | 0.704 | 0.781 | 8.28457879432418e-39 | 4 | RPLP2 |
| **RPS10.4** | 1.10035192392738e-42 | -0.414186408116793 | 0.518 | 0.654 | 1.39315557088446e-38 | 4 | RPS10 |
| **RPL27A.4** | 1.21896753002435e-42 | -0.436607278342238 | 0.557 | 0.68 | 1.54333478976383e-38 | 4 | RPL27A |
| **RPL37A.4** | 1.43204816901818e-42 | -0.413315605309112 | 0.647 | 0.759 | 1.81311618679392e-38 | 4 | RPL37A |
| **RPS14.4** | 1.43892471728476e-42 | -0.426319887585576 | 0.712 | 0.808 | 1.82182258455424e-38 | 4 | RPS14 |
| **CRIP1.4** | 1.31898584633087e-40 | -0.468962653931503 | 0.108 | 0.252 | 1.66996798003952e-36 | 4 | CRIP1 |
| **RPL12.3** | 2.8255525408514e-40 | -0.381359046692976 | 0.456 | 0.614 | 3.57743207197195e-36 | 4 | RPL12 |
| **CTSV.2** | 1.52645331766672e-39 | -0.342931436495433 | 0.039 | 0.309 | 1.93264254549784e-35 | 4 | CTSV |
| **ENSGALG00000011190.4** | 3.21435591357352e-39 | 0.554991313689167 | 0.836 | 0.35 | 4.06969602217543e-35 | 4 | ENSGALG00000011190 |
| **EEF1D.4** | 7.20084904297963e-39 | -0.395775872481021 | 0.313 | 0.475 | 9.1169949733165e-35 | 4 | EEF1D |
| **GSTA3.1** | 8.9069883682195e-39 | -0.376209654039261 | 0.028 | 0.332 | 1.12771379730027e-34 | 4 | GSTA3 |
| **LYN.1** | 1.78254969018032e-38 | 0.260078989276697 | 0.633 | 0.415 | 2.25688616273731e-34 | 4 | LYN |
| **RSFR.2** | 2.74480997007838e-38 | -0.390577983253035 | 0.025 | 0.327 | 3.47520390311623e-34 | 4 | RSFR |
| **RPS13.3** | 6.087438306247e-38 | -0.353755161055127 | 0.466 | 0.623 | 7.70730563953932e-34 | 4 | RPS13 |
| **PRDX1.2** | 6.80117637465911e-38 | -0.298167856023725 | 0.187 | 0.415 | 8.6109694079559e-34 | 4 | PRDX1 |
| **PPIA.1** | 1.61830622090078e-37 | -0.31588849552999 | 0.422 | 0.621 | 2.04893750628248e-33 | 4 | PPIA |
| **CTSC.2** | 2.62072105198093e-37 | -0.289161165285797 | 0.129 | 0.362 | 3.31809492391306e-33 | 4 | CTSC |
| **ENSGALG00000045115.2** | 3.43372171378197e-37 | -0.323633306726703 | 0.034 | 0.29 | 4.34743506181936e-33 | 4 | ENSGALG00000045115 |
| **RPS27A.4** | 4.84589959690055e-37 | -0.392884072479353 | 0.599 | 0.713 | 6.13539347963578e-33 | 4 | RPS27A |
| **LY86.4** | 5.87621424510807e-37 | -0.392908154361152 | 0.109 | 0.257 | 7.43987485573133e-33 | 4 | LY86 |
| **RPL19.4** | 6.23525100004124e-37 | -0.371788738963521 | 0.522 | 0.661 | 7.89445129115222e-33 | 4 | RPL19 |
| **RPS28.4** | 1.38694108117309e-36 | -0.366318904157296 | 0.469 | 0.617 | 1.75600610287324e-32 | 4 | RPS28 |
| **BRI3.2** | 1.40166107056883e-36 | -0.285277513268978 | 0.166 | 0.396 | 1.7746430814472e-32 | 4 | BRI3 |
| **EEF1A1.3** | 1.89279710337733e-36 | -0.366440817685407 | 0.76 | 0.824 | 2.39647041258604e-32 | 4 | EEF1A1 |
| **RPS24.4** | 2.53274664394161e-36 | -0.369052925281327 | 0.576 | 0.692 | 3.20671052589447e-32 | 4 | RPS24 |
| **RPS3A.4** | 2.68088954359705e-36 | -0.381182789699179 | 0.641 | 0.727 | 3.39427425114822e-32 | 4 | RPS3A |
| **RPL34.4** | 3.24294433199669e-36 | -0.383882119916064 | 0.644 | 0.733 | 4.10589181874101e-32 | 4 | RPL34 |
| **FAU.3** | 3.91453160300242e-36 | -0.342887212923156 | 0.353 | 0.533 | 4.95618846256136e-32 | 4 | FAU |
| **ITM2A.3** | 6.82496073676683e-36 | -0.351878201306986 | 0.414 | 0.553 | 8.64108278882048e-32 | 4 | ITM2A |
| **RPL7.3** | 7.76575038738572e-36 | -0.361695127790092 | 0.444 | 0.603 | 9.83221656546906e-32 | 4 | RPL7 |
| **RPL35A.4** | 9.44953997111894e-36 | -0.37493407448536 | 0.666 | 0.744 | 1.19640625574337e-31 | 4 | RPL35A |

| | | | | | | | |
|---|---|---|---|---|---|---|---|
| **LCP1.2** | 2.42290617750941e-35 | 0.261781420168216 | 0.725 | 0.529 | 3.06764151134466e-31 | 4 | LCP1 |
| **RPS15A.4** | 2.50488131094552e-35 | -0.371991810402937 | 0.691 | 0.789 | 3.17143022778813e-31 | 4 | RPS15A |
| **RPS12.4** | 3.4406884224128e-35 | -0.361953415903225 | 0.414 | 0.547 | 4.35625561161685e-31 | 4 | RPS12 |
| **RPS21.3** | 1.56685225874394e-34 | -0.343739670330156 | 0.399 | 0.546 | 1.9837916447957e-30 | 4 | RPS21 |
| **RPL7A.4** | 1.64585880487845e-34 | -0.359661827706505 | 0.477 | 0.599 | 2.08382183285661e-30 | 4 | RPL7A |
| **RPL24.4** | 1.82718141473342e-34 | -0.367508518144223 | 0.597 | 0.693 | 2.31339438919398e-30 | 4 | RPL24 |
| **COII.2** | 9.06127852936428e-34 | -0.310478866509157 | 0.991 | 0.985 | 1.14724847460281e-29 | 4 | COII |
| **RPL18A.3** | 2.52654580876934e-33 | -0.346628403544179 | 0.459 | 0.576 | 3.19885964848286e-29 | 4 | RPL18A |
| **SH3BGRL3.3** | 3.58645998409362e-33 | 0.275790569554568 | 0.593 | 0.372 | 4.54081698586094e-29 | 4 | SH3BGRL3 |
| **BTG1.3** | 2.46221516427556e-32 | -0.278490878512468 | 0.152 | 0.341 | 3.11741061948929e-28 | 4 | BTG1 |
| **RPL26L1.4** | 3.609348634166e-32 | -0.359796489726595 | 0.537 | 0.65 | 4.56979630571758e-28 | 4 | RPL26L1 |
| **PSAP.2** | 1.08424573383958e-31 | -0.273476246219093 | 0.189 | 0.39 | 1.3727635236143e-27 | 4 | PSAP |
| **RPL11.3** | 1.57678220742772e-31 | -0.326981920869085 | 0.553 | 0.664 | 1.99636395282423e-27 | 4 | RPL11 |
| **C1QA.3** | 3.5656448271172e-31 | -0.290768393901589 | 0.307 | 0.482 | 4.51446291561309e-27 | 4 | C1QA |
| **SCPEP1.2** | 7.99758627444379e-31 | -0.254476678943483 | 0.122 | 0.315 | 1.01257439820733e-26 | 4 | SCPEP1 |
| **RPS2.3** | 8.84522358648021e-31 | -0.308582261703022 | 0.501 | 0.628 | 1.11989375828426e-26 | 4 | RPS2 |
| **RPS8.4** | 8.15898151274505e-30 | -0.332199694306303 | 0.77 | 0.827 | 1.03300864932865e-25 | 4 | RPS8 |
| **RPL32.4** | 1.05067593906276e-29 | -0.323388736292723 | 0.573 | 0.676 | 1.33026080644737e-25 | 4 | RPL32 |
| **RPL22.3** | 1.39786432058973e-29 | -0.310142050324014 | 0.494 | 0.607 | 1.76983601629866e-25 | 4 | RPL22 |
| **UBA52.3** | 2.26698980793872e-29 | -0.309570777422535 | 0.46 | 0.582 | 2.87023579583121e-25 | 4 | UBA52 |
| **RPL8.4** | 2.3008441096137e-29 | -0.332775566894123 | 0.543 | 0.642 | 2.91309872718191e-25 | 4 | RPL8 |
| **GAPDH.3** | 5.18711459017836e-29 | -0.312300497120985 | 0.409 | 0.536 | 6.56740578262482e-25 | 4 | GAPDH |
| **RPL13.4** | 5.72382959360413e-29 | -0.325639292476696 | 0.547 | 0.637 | 7.24694064846219e-25 | 4 | RPL13 |
| **ENSGALG00000004509.4** | 1.07302301238161e-28 | -0.29996839854255 | 0.813 | 0.871 | 1.35855443597636e-24 | 4 | ENSGALG00000004509 |
| **TSC22D4.4** | 1.09636596883089e-28 | -0.275259738769331 | 0.134 | 0.296 | 1.38810895313679e-24 | 4 | TSC22D4 |
| **RPL10A.3** | 1.30067010916226e-28 | -0.296882314688737 | 0.424 | 0.559 | 1.64677842521034e-24 | 4 | RPL10A |
| **RPL17.4** | 2.14188887926442e-28 | -0.320000678153036 | 0.7 | 0.755 | 2.71184551003668e-24 | 4 | RPL17 |
| **RPL36.4** | 7.35165219687712e-28 | -0.309324959297123 | 0.435 | 0.545 | 9.30792684646612e-24 | 4 | RPL36 |
| **SLC25A6.2** | 1.21065230710793e-27 | -0.252653705413331 | 0.181 | 0.351 | 1.53280688602935e-23 | 4 | SLC25A6 |
| **ZEB2.4** | 1.2444622528676e-27 | -0.268559585256435 | 0.168 | 0.323 | 1.57561365835567e-23 | 4 | ZEB2 |
| **ENSGALG00000015183.3** | 1.68621239915779e-27 | 0.283224285499712 | 0.556 | 0.273 | 2.13491351857368e-23 | 4 | ENSGALG00000015183 |
| **RPL30.3** | 5.93479508515296e-27 | -0.30622132175932 | 0.48 | 0.585 | 7.51404405731216e-23 | 4 | RPL30 |
| **CTSZ.2** | 1.18130847438399e-26 | -0.252727295973946 | 0.238 | 0.393 | 1.49565465941757e-22 | 4 | CTSZ |
| **RPS20.4** | 1.26126853360066e-25 | -0.298779539331099 | 0.677 | 0.758 | 1.59689209039179e-21 | 4 | RPS20 |
| **NME2.2** | 1.85173299551985e-25 | -0.267030771275716 | 0.353 | 0.495 | 2.34447914562768e-21 | 4 | NME2 |
| **RPS23.3** | 4.53072073592645e-25 | -0.310970968370062 | 0.648 | 0.707 | 5.73634552375648e-21 | 4 | RPS23 |
| **RPL36AL.4** | 5.56880477038046e-25 | -0.289038378282111 | 0.53 | 0.633 | 7.05066371977869e-21 | 4 | RPL36AL |
| **RPS25.3** | 7.32053643440864e-25 | -0.293004467165638 | 0.552 | 0.641 | 9.26853117960477e-21 | 4 | RPS25 |
| **RPS3.3** | 1.30149684457888e-24 | -0.282130216064871 | 0.411 | 0.529 | 1.64782515492132e-20 | 4 | RPS3 |
| **RPL3.3** | 3.21355730836188e-24 | -0.261167375014926 | 0.389 | 0.515 | 4.06868490811698e-20 | 4 | RPL3 |
| **ND3.2** | 4.17698333276317e-23 | -0.284220558460508 | 0.796 | 0.825 | 5.28847859761145e-19 | 4 | ND3 |
| **RPS4Y1.4** | 5.33964594244204e-22 | -0.271411970014993 | 0.452 | 0.539 | 6.76052572772587e-18 | 4 | RPS4Y1 |
| **RPL9.3** | 8.83663064077769e-22 | -0.262959298638003 | 0.484 | 0.588 | 1.11880580542886e-17 | 4 | RPL9 |
| **RPL31.3** | 1.84429398662182e-21 | -0.268360644303727 | 0.531 | 0.617 | 2.33506061646188e-17 | 4 | RPL31 |
| **RPL37.4** | 1.95009063854199e-21 | -0.290834694444476 | 0.699 | 0.712 | 2.46900975745802e-17 | 4 | RPL37 |
| **RPS6.3** | 3.7412861975269e-21 | -0.267241277973733 | 0.568 | 0.631 | 4.7368424546888e-17 | 4 | RPS6 |
| **RPS16.4** | 4.65785122430118e-20 | -0.271173360204781 | 0.619 | 0.664 | 5.89730543508772e-16 | 4 | RPS16 |
| **FOS.4** | 1.39239278963841e-15 | -0.274662284208398 | 0.367 | 0.395 | 1.76290851096119e-11 | 4 | FOS |
| **CTSD.3** | 9.01201994551277e-10 | -0.259203449088566 | 0.643 | 0.473 | 1.14101184530137e-05 | 4 | CTSD |
| **FTH1.4** | 3.30705999114945e-289 | -1.04406594934011 | 0.973 | 0.993 | 4.18706865479432e-285 | 5 | FTH1 |
| **IFI30.4** | 1.33892362471471e-272 | -1.19851039289545 | 0.506 | 0.856 | 1.6952112012513e-268 | 5 | IFI30 |
| **TMSB4X.3** | 1.19845940358707e-203 | 0.688531668361205 | 0.999 | 0.993 | 1.51736945088159e-199 | 5 | TMSB4X |
| **CD74.3** | 3.52869448304346e-144 | -0.673672281805426 | 0.975 | 0.997 | 4.46768008498132e-140 | 5 | CD74 |
| **CTSS.3** | 2.34930572269231e-143 | -0.830898315680725 | 0.208 | 0.699 | 2.97445597550073e-139 | 5 | CTSS |

| | | | | | | | |
|---|---|---|---|---|---|---|---|
| **ENSGALG00000015183.4** | 3.71532526648264e-123 | 0.590846970674656 | 0.689 | 0.26 | 4.70397331989367e-119 | 5 | ENSGALG00000015183 |
| **C1QC.3** | 2.02134272040076e-111 | -0.758695201224848 | 0.118 | 0.588 | 2.55922201829941e-107 | 5 | C1QC |
| **ENSGALG00000011190.5** | 4.37868705780341e-106 | 0.903828479601263 | 0.916 | 0.343 | 5.5438556838849e-102 | 5 | ENSGALG00000011190 |
| **BLB2.4** | 3.46981819885109e-101 | -0.589211284777397 | 0.633 | 0.886 | 4.39313682156536e-97 | 5 | BLB2 |
| **SAT1.4** | 3.68116995487082e-97 | -0.675399912288177 | 0.222 | 0.588 | 4.66072927986194e-93 | 5 | SAT1 |
| **BLB1.4** | 2.86371311150658e-96 | -0.616691159310964 | 0.668 | 0.879 | 3.62574717047848e-92 | 5 | BLB1 |
| **GRN.4** | 2.78826536643773e-85 | -0.560531883016152 | 0.284 | 0.661 | 3.5302227804468e-81 | 5 | GRN |
| **ITM2A.4** | 7.06344228839152e-85 | -0.583147032306494 | 0.224 | 0.571 | 8.9430242813325e-81 | 5 | ITM2A |
| **CTSD.4** | 1.47027891825787e-84 | -0.764757488081559 | 0.224 | 0.513 | 1.86152013840629e-80 | 5 | CTSD |
| **ATP6V0C.5** | 4.97739577330535e-84 | -0.579535632499847 | 0.246 | 0.585 | 6.30188078858191e-80 | 5 | ATP6V0C |
| **PPDPF.5** | 7.93517459064613e-83 | -0.826875725085676 | 0.064 | 0.39 | 1.00467245492171e-78 | 5 | PPDPF |
| **CSTA.1** | 6.47279902293451e-82 | 0.311466217319812 | 0.357 | 0.086 | 8.19521084293738e-78 | 5 | CSTA |
| **EXFABP.5** | 4.8513680534229e-76 | -1.35466968011312 | 0.029 | 0.254 | 6.14231709243874e-72 | 5 | EXFABP |
| **C1QB.4** | 1.98257352926553e-75 | -0.572317776371631 | 0.073 | 0.471 | 2.51013634540308e-71 | 5 | C1QB |
| **S100A6.5** | 1.24598571361994e-73 | -0.842117058550842 | 0.06 | 0.309 | 1.57754251201421e-69 | 5 | S100A6 |
| **SLC40A1.5** | 3.4608383447172e-69 | -0.525739351201676 | 0.127 | 0.451 | 4.38176742824645e-65 | 5 | SLC40A1 |
| **MAFB.4** | 4.90146048948187e-69 | -0.537019252020768 | 0.386 | 0.641 | 6.20573912573299e-65 | 5 | MAFB |
| **HSP90AA1.4** | 1.23691992262485e-68 | -0.588067118928725 | 0.718 | 0.794 | 1.56606431403532e-64 | 5 | HSP90AA1 |
| **FOS.5** | 2.7196200685356e-68 | -0.585106899795215 | 0.153 | 0.416 | 3.44331096877292e-64 | 5 | FOS |
| **ENSGALG00000014585.4** | 1.43771291843247e-65 | -0.53869008124201 | 0.068 | 0.413 | 1.82028832602735e-61 | 5 | ENSGALG00000014585 |
| **TMSB15B.4** | 1.89571259243315e-65 | -0.539885509238267 | 0.505 | 0.73 | 2.40016171327961e-61 | 5 | TMSB15B |
| **HMOX1.4** | 2.23303577257215e-65 | -0.57500477788128 | 0.089 | 0.383 | 2.82724659165359e-61 | 5 | HMOX1 |
| **COX3.3** | 1.05960471868299e-64 | -0.442692338405364 | 0.959 | 0.992 | 1.34156553432453e-60 | 5 | COX3 |
| **B2M.2** | 1.36530857956254e-63 | -0.398206619414432 | 0.845 | 0.94 | 1.72861719258414e-59 | 5 | B2M |
| **SPIC.3** | 1.83097916330823e-63 | 0.479351610589018 | 0.622 | 0.165 | 2.31820271866455e-59 | 5 | SPIC |
| **MT-CO1.3** | 1.79298670729608e-61 | -0.461367003584956 | 0.93 | 0.981 | 2.27010047010757e-57 | 5 | MT-CO1 |
| **TPP1.3** | 4.5195008985216e-61 | -0.438248592690105 | 0.105 | 0.454 | 5.7221400876182e-57 | 5 | TPP1 |
| **C1QA.4** | 2.39272581646725e-60 | -0.435119094671271 | 0.197 | 0.492 | 3.02943015622918e-56 | 5 | C1QA |
| **ASS1.3** | 5.52793506358684e-60 | -0.472848010769643 | 0.088 | 0.403 | 6.99891858400729e-56 | 5 | ASS1 |
| **PNRC1.5** | 2.41474046462616e-57 | -0.4493745933592 | 0.324 | 0.564 | 3.05730290226318e-53 | 5 | PNRC1 |
| **HSPA5.4** | 2.78541516913037e-57 | -0.495846847583423 | 0.166 | 0.423 | 3.52661414563597e-53 | 5 | HSPA5 |
| **JUN.4** | 5.22427125252382e-57 | -0.502934182967041 | 0.068 | 0.341 | 6.61444983282041e-53 | 5 | JUN |
| **TXN.3** | 1.48309057986221e-56 | -0.447715345657727 | 0.259 | 0.515 | 1.87774098316355e-52 | 5 | TXN |
| **CTSB.3** | 5.82030108875029e-56 | -0.451412245893436 | 0.056 | 0.409 | 7.36908320846674e-52 | 5 | CTSB |
| **CST7.1** | 1.5255058284933e-55 | 0.30555806299754 | 0.416 | 0.156 | 1.93144292945537e-51 | 5 | CST7 |
| **GNG10** | 3.2025452297152e-54 | 0.269662481290389 | 0.363 | 0.183 | 4.05474251534241e-50 | 5 | GNG10 |
| **HSPA8.4** | 7.57496431466079e-54 | -0.481617612224483 | 0.491 | 0.635 | 9.59066231879203e-50 | 5 | HSPA8 |
| **CTSZ.3** | 5.34935591115121e-52 | -0.390154638410691 | 0.106 | 0.406 | 6.77281951910855e-48 | 5 | CTSZ |
| **CTSA.3** | 1.47440945263948e-51 | -0.39264658717825 | 0.084 | 0.389 | 1.86674980798685e-47 | 5 | CTSA |
| **TGM4.5** | 6.38596444217504e-49 | 0.820851098179951 | 0.922 | 0.281 | 8.08526958023782e-45 | 5 | TGM4 |
| **SH3BGRL3.4** | 2.26355731430867e-48 | 0.334654676476361 | 0.635 | 0.368 | 2.86588991564621e-44 | 5 | SH3BGRL3 |
| **ZEB2.5** | 4.98227021483577e-47 | -0.391885137762018 | 0.065 | 0.333 | 6.30805231900357e-43 | 5 | ZEB2 |
| **PSAP.3** | 9.22048038425668e-47 | -0.354076889723553 | 0.116 | 0.397 | 1.16740502145074e-42 | 5 | PSAP |
| **ATP6.3** | 1.76456471803248e-46 | -0.391314640065117 | 0.931 | 0.974 | 2.23411538950092e-42 | 5 | ATP6 |
| **CD47.4** | 5.70883914616542e-46 | -0.352317215249011 | 0.108 | 0.385 | 7.22796124296004e-42 | 5 | CD47 |
| **LAMP1.2** | 4.29676516869842e-45 | -0.341912767292658 | 0.07 | 0.367 | 5.44013438008907e-41 | 5 | LAMP1 |
| **CTSC.3** | 4.78089918828596e-45 | -0.340842703099079 | 0.081 | 0.367 | 6.05309646228885e-41 | 5 | CTSC |
| **CST3.3** | 1.8752124641735e-42 | -0.363697721481371 | 0.483 | 0.733 | 2.37420650089007e-38 | 5 | CST3 |
| **RSFR.3** | 2.6303399640225e-42 | -0.372185264042408 | 0.04 | 0.326 | 3.33027342844888e-38 | 5 | RSFR |
| **GSTA3.2** | 5.54648070867235e-42 | -0.357828243709174 | 0.042 | 0.33 | 7.02239922525007e-38 | 5 | GSTA3 |
| **LY86.5** | 4.22669119699226e-41 | -0.418085495563944 | 0.086 | 0.259 | 5.3514137245119e-37 | 5 | LY86 |
| **COII.3** | 1.90500849676316e-40 | -0.343058267784546 | 0.955 | 0.988 | 2.41193125775183e-36 | 5 | COII |
| **BTG2.4** | 8.68955429514138e-40 | -0.36054270503991 | 0.131 | 0.333 | 1.10018446930785e-35 | 5 | BTG2 |
| **TSC22D4.5** | 1.92447483116922e-39 | -0.33634213126257 | 0.088 | 0.3 | 2.43657758374335e-35 | 5 | TSC22D4 |

| | | | | | | | |
|---|---|---|---|---|---|---|---|
| **CTSV.3** | 3.01877816596958e-38 | -0.349023366400565 | 0.032 | 0.309 | 3.82207503593409e-34 | 5 | CTSV |
| **ENSGALG00000045115.3** | 1.67326717908665e-36 | -0.299939998567748 | 0.06 | 0.287 | 2.11852357544161e-32 | 5 | ENSGALG00000045115 |
| **CD81.3** | 4.84033753723648e-36 | -0.301613646825603 | 0.068 | 0.28 | 6.12835135589511e-32 | 5 | CD81 |
| **NFKBIA.4** | 4.74216772447257e-35 | -0.312827243604131 | 0.158 | 0.353 | 6.00405855595473e-31 | 5 | NFKBIA |
| **LSP1P1.3** | 6.09669798062985e-35 | -0.316693981374202 | 0.189 | 0.387 | 7.71902931327545e-31 | 5 | LSP1P1 |
| **GNS 0.20** | 1.21740561184183e-34 | -0.270094055511737 | 0.134 | 0.362 | 1.54135724515294e-30 | 5 | GNS |
| **HSP90B1.4** | 4.04936201936899e-34 | -0.320761643199266 | 0.259 | 0.457 | 5.12689725272307e-30 | 5 | HSP90B1 |
| **DDX5.4** | 1.38920125120606e-33 | -0.334866877509294 | 0.382 | 0.532 | 1.75886770415199e-29 | 5 | DDX5 |
| **ND3.3** | 1.60554753952666e-33 | -0.34919956949966 | 0.717 | 0.833 | 2.03278373979471e-29 | 5 | ND3 |
| **SCPEP1.3** | 1.77481382197251e-33 | -0.270624491266664 | 0.098 | 0.317 | 2.2470917799994e-29 | 5 | SCPEP1 |
| **ACTB.2** | 2.665801727429e-33 | 0.269075060249213 | 0.991 | 0.979 | 3.37517156709785e-29 | 5 | ACTB |
| **S100A10.5** | 1.10361412959997e-32 | -0.482780135019386 | 0.349 | 0.389 | 1.39728584948652e-28 | 5 | S100A10 |
| **PPIB.2** | 1.16043969372359e-32 | -0.2700503072139 | 0.166 | 0.38 | 1.46923269622344e-28 | 5 | PPIB |
| **ANKRD11.3** | 1.21057801563581e-32 | -0.290785599660203 | 0.15 | 0.348 | 1.5327128255965e-28 | 5 | ANKRD11 |
| **CREG1.1** | 2.92976580700122e-32 | -0.259095232491555 | 0.037 | 0.277 | 3.70937648824425e-28 | 5 | CREG1 |
| **TNFAIP2.2** | 4.53619113370219e-32 | 0.287793839073455 | 0.409 | 0.155 | 5.74327159438034e-28 | 5 | TNFAIP2 |
| **FGL2.2** | 3.6175790914054e-31 | -0.257056662665829 | 0.058 | 0.256 | 4.58021688762838e-27 | 5 | FGL2 |
| **NPC2** | 3.01278091818959e-30 | -0.259097596397361 | 0.255 | 0.463 | 3.81448192051984e-26 | 5 | NPC2 |
| **BTG1.4** | 1.37751554589258e-29 | -0.265702097791758 | 0.147 | 0.341 | 1.7440724326546e-25 | 5 | BTG1 |
| **GRP.2** | 4.65318694335606e-29 | 0.296672581756584 | 0.424 | 0.142 | 5.89139998898311e-25 | 5 | GRP |
| **ND5.2** | 1.29215240283171e-26 | -0.287431285918289 | 0.756 | 0.854 | 1.63599415722523e-22 | 5 | ND5 |
| **EEF2.3** | 8.75852627560074e-24 | -0.289566335131997 | 0.42 | 0.532 | 1.10891701175381e-19 | 5 | EEF2 |
| **ND1.2** | 8.58244777420244e-22 | -0.25246486454693 | 0.688 | 0.821 | 1.08662371269177e-17 | 5 | ND1 |
| **ND4.2** | 2.08310817724479e-21 | -0.252014155541629 | 0.649 | 0.783 | 2.63742326320963e-17 | 5 | ND4 |
| **GAPDH.4** | 3.84951977716101e-19 | -0.250453540561921 | 0.433 | 0.533 | 4.87387698986356e-15 | 5 | GAPDH |
| **CXCL13.5** | 1.37659348883761e-16 | 0.649924532868484 | 0.983 | 0.368 | 1.7429050162173e-12 | 5 | CXCL13 |
| **TPT1.5** | 2.69066981895057e-15 | -0.256363751794699 | 0.794 | 0.801 | 3.40665705777331e-11 | 5 | TPT1 |
| **MARCO.5** | 5.22560003927419e-08 | 0.273426914526267 | 0.71 | 0.348 | 0.000661613220972505 | 5 | MARCO |
| **CTSD.5** | 2.31111601703503e-174 | 1.27658963150212 | 0.972 | 0.447 | 2.92610398916805e-170 | 6 | CTSD |
| **CXCL13.6** | 5.51808918039397e-128 | -1.51882337903324 | 0.216 | 0.439 | 6.9864527112968e-124 | 6 | CXCL13 |
| **LGMN** | 1.22635152732654e-115 | 0.485421619922158 | 0.596 | 0.189 | 1.55268366874813e-111 | 6 | LGMN |
| **TMSB4X.4** | 2.96610536129352e-88 | -0.525067011368009 | 0.987 | 0.994 | 3.75538599793373e-84 | 6 | TMSB4X |
| **HMOX1.5** | 2.68026499200526e-87 | 0.818031752818504 | 0.776 | 0.322 | 3.39348350637786e-83 | 6 | HMOX1 |
| **HSP90AA1.5** | 3.19901587413338e-86 | -0.700231014039136 | 0.659 | 0.798 | 4.05027399824028e-82 | 6 | HSP90AA1 |
| **RPS11.5** | 5.32233090453184e-84 | -0.720862756641219 | 0.633 | 0.798 | 6.73860315822776e-80 | 6 | RPS11 |
| **ENSGALG00000005204** | 4.05611142812638e-81 | 0.267959270907431 | 0.317 | 0.073 | 5.13544267915081e-77 | 6 | ENSGALG00000005204 |
| **RPS17.5** | 9.30401181093767e-81 | -0.615217827442166 | 0.674 | 0.838 | 1.17798093538282e-76 | 6 | RPS17 |
| **RPS19.5** | 3.79078687514204e-80 | -0.645044766589763 | 0.76 | 0.857 | 4.79951526261733e-76 | 6 | RPS19 |
| **ENSGALG00000037441.5** | 4.66462853224611e-77 | -0.624693987510435 | 0.595 | 0.794 | 5.9058861846768e-73 | 6 | ENSGALG00000037441 |
| **RPL37.5** | 3.37333503159976e-76 | -0.610257945371734 | 0.499 | 0.729 | 4.27097948350845e-72 | 6 | RPL37 |
| **TGM4.6** | 1.27732882761525e-75 | -0.834553397630704 | 0.075 | 0.359 | 1.61722602864367e-71 | 6 | TGM4 |
| **RPL39L.3** | 4.14201259420803e-73 | -0.527487248983132 | 0.689 | 0.842 | 5.24420214552679e-69 | 6 | RPL39L |
| **ENSGALG00000026970.3** | 4.90748158179911e-73 | -0.582993005997098 | 0.235 | 0.562 | 6.21336243071586e-69 | 6 | ENSGALG00000026970 |
| **RPS14.5** | 1.57672467930686e-72 | -0.600766940366157 | 0.633 | 0.814 | 1.99629111647042e-68 | 6 | RPS14 |
| **RPL21.5** | 2.26384144362171e-72 | -0.566904542897823 | 0.703 | 0.839 | 2.86624965176945e-68 | 6 | RPL21 |
| **EXFABP.6** | 2.63009234706112e-72 | -1.33757942280423 | 0.036 | 0.251 | 3.32995992061409e-68 | 6 | EXFABP |
| **RPL34.5** | 1.26716310225463e-70 | -0.583445620387468 | 0.566 | 0.739 | 1.60435520376458e-66 | 6 | RPL34 |
| **SLC40A1.6** | 7.17943454422328e-70 | 0.545526075366065 | 0.845 | 0.387 | 9.0898820764411e-66 | 6 | SLC40A1 |
| **RPS15A.5** | 3.04749019908344e-69 | -0.567038253924444 | 0.63 | 0.793 | 3.85842734105954e-65 | 6 | RPS15A |
| **RPS3A.5** | 5.1854193245845e-69 | -0.572134771533438 | 0.526 | 0.736 | 6.56525940685643e-65 | 6 | RPS3A |
| **DDX5.5** | 9.23361661652591e-69 | -0.539253305342885 | 0.231 | 0.543 | 1.16906819981835e-64 | 6 | DDX5 |
| **HSPA8.5** | 1.67269354815451e-68 | -0.58098601772098 | 0.41 | 0.641 | 2.11779730131842e-64 | 6 | HSPA8 |
| **TPT1.6** | 4.7394665002776e-68 | -0.599485471299196 | 0.69 | 0.809 | 6.00063853600147e-64 | 6 | TPT1 |
| **CD74.4** | 9.04789739141448e-68 | -0.484676974094882 | 0.99 | 0.995 | 1.14555428872699e-63 | 6 | CD74 |

| | | | | | | | |
|---|---|---|---|---|---|---|---|
| **RPL15.5** | 2.54529936322022e-67 | -0.553781684806574 | 0.636 | 0.783 | 3.22260352377313e-63 | 6 | RPL15 |
| **RPS8.5** | 4.41279007156899e-67 | -0.545605951202094 | 0.709 | 0.831 | 5.5870335096135e-63 | 6 | RPS8 |
| **FTH1.5** | 7.03073200826675e-67 | 0.517877021033159 | 1 | 0.99 | 8.90160979566653e-63 | 6 | FTH1 |
| **RPLP2.5** | 1.31078563230179e-66 | -0.59747915313623 | 0.647 | 0.785 | 1.6595856890573e-62 | 6 | RPLP2 |
| **RPS15.5** | 2.62996012806798e-66 | -0.573699781639674 | 0.633 | 0.782 | 3.32979251814687e-62 | 6 | RPS15 |
| **FOS.6** | 3.46546536504795e-64 | -0.603323467501985 | 0.145 | 0.414 | 4.38762569868721e-60 | 6 | FOS |
| **RPL23.5** | 1.73628698848682e-63 | -0.543350559700778 | 0.656 | 0.809 | 2.19831295612316e-59 | 6 | RPL23 |
| **EEF1A1.4** | 2.40334113994114e-63 | -0.52443946128921 | 0.694 | 0.829 | 3.04287021727947e-59 | 6 | EEF1A1 |
| **RPL17.5** | 8.30847197485215e-63 | -0.52280813555815 | 0.581 | 0.765 | 1.05193563673603e-58 | 6 | RPL17 |
| **RPL35A.5** | 1.45191049067429e-61 | -0.533322352193066 | 0.577 | 0.751 | 1.83826387224272e-57 | 6 | RPL35A |
| **RPLP1.5** | 1.93158240231768e-61 | -0.525498917882085 | 0.857 | 0.91 | 2.44557647957442e-57 | 6 | RPLP1 |
| **S100A6.6** | 3.76132548452829e-61 | -0.776755070737795 | 0.098 | 0.303 | 4.76221419596127e-57 | 6 | S100A6 |
| **RPS16.5** | 7.89633779063211e-61 | -0.524746306805161 | 0.466 | 0.676 | 9.99755327671932e-57 | 6 | RPS16 |
| **RPL8.5** | 5.08696662018726e-60 | -0.520451033954866 | 0.431 | 0.651 | 6.44060843781909e-56 | 6 | RPL8 |
| **RPL27A.5** | 3.33625434405842e-59 | -0.551111114830017 | 0.493 | 0.684 | 4.22403162501237e-55 | 6 | RPL27A |
| **FABP7** | 4.23050716156411e-59 | 0.253258358328442 | 0.346 | 0.103 | 5.35624511725632e-55 | 6 | FABP7 |
| **RPS10.5** | 5.36745053819014e-59 | -0.524229710176901 | 0.458 | 0.658 | 6.79572912640253e-55 | 6 | RPS10 |
| **GSTA3.3** | 6.1231296067285e-59 | 0.367156331783708 | 0.651 | 0.276 | 7.75249439507895e-55 | 6 | GSTA3 |
| **RPS27A.5** | 1.33402666343574e-58 | -0.532639480066047 | 0.563 | 0.715 | 1.68901115857599e-54 | 6 | RPS27A |
| **RPL37A.5** | 1.37982966493911e-58 | -0.521187705135142 | 0.635 | 0.759 | 1.74700233877941e-54 | 6 | RPL37A |
| **RPL26L1.5** | 3.06179298910146e-58 | -0.524815702443008 | 0.459 | 0.655 | 3.87653610350136e-54 | 6 | RPL26L1 |
| **RPL13.5** | 1.12112522996619e-56 | -0.498057730556931 | 0.428 | 0.646 | 1.4194566536602e-52 | 6 | RPL13 |
| **ITGB1BP3.3** | 1.49930211962472e-56 | -0.474753218667402 | 0.285 | 0.556 | 1.89826641365685e-52 | 6 | ITGB1BP3 |
| **RPL7A.5** | 1.05537284205028e-55 | -0.495807836606471 | 0.386 | 0.606 | 1.33620755531987e-51 | 6 | RPL7A |
| **ACTB.3** | 2.70120877632513e-55 | -0.391609736750864 | 0.958 | 0.982 | 3.42000043170525e-51 | 6 | ACTB |
| **GNB2L1.5** | 5.58930706255335e-55 | -0.512732530717147 | 0.471 | 0.643 | 7.07662167189879e-51 | 6 | GNB2L1 |
| **ENSGALG00000011190.6** | 7.02010561309579e-55 | -0.589230329435289 | 0.198 | 0.409 | 8.88815571674058e-51 | 6 | ENSGALG00000011190 |
| **MARCO.6** | 8.86673172752973e-55 | -0.669252163995084 | 0.228 | 0.393 | 1.12261690402254e-50 | 6 | MARCO |
| **RPL24.5** | 5.57153934249066e-54 | -0.496644896055952 | 0.534 | 0.697 | 7.05412596152743e-50 | 6 | RPL24 |
| **S100A10.6** | 5.68536354170436e-53 | -0.630861837010695 | 0.224 | 0.4 | 7.19823878015189e-49 | 6 | S100A10 |
| **RPS23.4** | 1.58894628569234e-52 | -0.490830504414954 | 0.542 | 0.715 | 2.01176489231507e-48 | 6 | RPS23 |
| **RPS24.5** | 2.43086901289945e-51 | -0.472980720561168 | 0.528 | 0.695 | 3.07772325723199e-47 | 6 | RPS24 |
| **RPS20.5** | 5.1879420732399e-51 | -0.46120611389438 | 0.617 | 0.762 | 6.56845345892904e-47 | 6 | RPS20 |
| **RPL35.4** | 2.51890436465763e-50 | -0.449203590020528 | 0.432 | 0.635 | 3.18918481609303e-46 | 6 | RPL35 |
| **RPL32.5** | 4.55257298634593e-50 | -0.456734446815155 | 0.499 | 0.681 | 5.76401265801258e-46 | 6 | RPL32 |
| **RPS6.4** | 9.81011978166492e-50 | -0.452722662584153 | 0.434 | 0.641 | 1.2420592655566e-45 | 6 | RPS6 |
| **RPS4Y1.5** | 6.36931371084315e-49 | -0.445974095441012 | 0.325 | 0.549 | 8.06418808929851e-45 | 6 | RPS4Y1 |
| **JUN.5** | 2.88308111325805e-48 | -0.532557974136531 | 0.04 | 0.341 | 3.65026899749601e-44 | 6 | JUN |
| **RPL19.5** | 3.2524308463733e-47 | -0.451278050011715 | 0.474 | 0.663 | 4.11790269459323e-43 | 6 | RPL19 |
| **RPL31.4** | 1.24210461696942e-46 | -0.435396815022757 | 0.419 | 0.626 | 1.57262865554498e-42 | 6 | RPL31 |
| **TMSB15B.5** | 1.37017317147468e-46 | 0.466193456994972 | 0.952 | 0.69 | 1.7347762524041e-42 | 6 | TMSB15B |
| **EEF2.4** | 2.05210042623443e-46 | -0.439233731172809 | 0.332 | 0.538 | 2.59816434965541e-42 | 6 | EEF2 |
| **RPL30.4** | 3.46826274536577e-46 | -0.435994617259353 | 0.408 | 0.59 | 4.3911674619076e-42 | 6 | RPL30 |
| **RPSAP58.2** | 5.72130472818148e-46 | -0.40505618473139 | 0.361 | 0.602 | 7.24374391635057e-42 | 6 | RPSAP58 |
| **BTG2.5** | 6.50028834284526e-46 | -0.436516689842403 | 0.069 | 0.336 | 8.23001507087638e-42 | 6 | BTG2 |
| **RPL36AL.5** | 7.26831706284876e-46 | -0.428224397253914 | 0.452 | 0.638 | 9.20241623327282e-42 | 6 | RPL36AL |
| **RPL7.4** | 1.29090855320627e-45 | -0.438215306506076 | 0.407 | 0.605 | 1.63441931921446e-41 | 6 | RPL7 |
| **RPL27.3** | 6.68094749013965e-45 | -0.415640323470583 | 0.337 | 0.552 | 8.45874761726581e-41 | 6 | RPL27 |
| **EEF1D.5** | 1.92793973363275e-44 | -0.450676717508246 | 0.283 | 0.476 | 2.44096449675243e-40 | 6 | EEF1D |
| **UBA52.4** | 2.91831064273411e-44 | -0.412703122019673 | 0.373 | 0.588 | 3.69487310476565e-40 | 6 | UBA52 |
| **RPS25.4** | 3.88720468662064e-44 | -0.425264859365915 | 0.478 | 0.646 | 4.92158985373O4e-40 | 6 | RPS25 |
| **RPS7.3** | 4.01904095641686e-44 | -0.413025237313554 | 0.365 | 0.575 | 5.08850775491939e-40 | 6 | RPS7 |
| **CRIP1.5** | 4.86603485233444e-44 | -0.549051312188531 | 0.042 | 0.256 | 6.16088672654064e-40 | 6 | CRIP1 |
| **RPS28.5** | 7.12763849142331e-44 | -0.429616660850978 | 0.444 | 0.617 | 9.02430309399106e-40 | 6 | RPS28 |

| | | | | | | | |
|---|---|---|---|---|---|---|---|
| **RPS26.5** | 1.52747391988918e-43 | -0.453883299015712 | 0.621 | 0.718 | 1.93393472997169e-39 | 6 | RPS26 |
| **CTSB.4** | 5.9212268850688e-43 | 0.338842092340816 | 0.663 | 0.354 | 7.49686535918561e-39 | 6 | CTSB |
| **BLB1.5** | 6.31132614641352e-43 | -0.42469655393676 | 0.815 | 0.865 | 7.99077003397416e-39 | 6 | BLB1 |
| **RPL23A.4** | 7.23816935455602e-43 | -0.426997210894551 | 0.447 | 0.627 | 9.16424621980338e-39 | 6 | RPL23A |
| **RPL29.3** | 8.84021322722104e-43 | -0.401640641977707 | 0.475 | 0.638 | 1.11925939669846e-38 | 6 | RPL29 |
| **RPL11.4** | 2.18699857215323e-42 | -0.408738803497899 | 0.505 | 0.667 | 2.76895889220321e-38 | 6 | RPL11 |
| **ENSGALG00000028749.[illegible]** | 1.54776557209884e-41 | -0.359242915587232 | 0.468 | 0.679 | 1.95962599083434e-37 | 6 | ENSGALG00000028749 |
| **RPL18A.4** | 2.75710861958899e-41 | -0.41350619335147 | 0.423 | 0.578 | 3.49077522326162e-37 | 6 | RPL18A |
| **RPL22.4** | 4.12495142881426e-41 | -0.395307321971818 | 0.428 | 0.611 | 5.22260100402174e-37 | 6 | RPL22 |
| **RPL36.5** | 1.86036650002932e-40 | -0.402254347492573 | 0.382 | 0.548 | 2.35541002568713e-36 | 6 | RPL36 |
| **ANKRD11.4** | 3.50822932755266e-40 | -0.35692427323033 | 0.092 | 0.351 | 4.44176915161443e-36 | 6 | ANKRD11 |
| **CTSS.4** | 3.97068050622808e-40 | 0.390158932131437 | 0.903 | 0.635 | 5.02727858893537e-36 | 6 | CTSS |
| **RPL12.4** | 1.29973290292084e-39 | -0.401877752239264 | 0.441 | 0.613 | 1.64559182838808e-35 | 6 | RPL12 |
| **RPL14.2** | 9.33606718605387e-39 | -0.369582118948022 | 0.368 | 0.576 | 1.18203946642628e-34 | 6 | RPL14 |
| **CSF1R.3** | 1.68395341705937e-38 | -0.331822566764289 | 0.189 | 0.432 | 2.13205342133886e-34 | 6 | CSF1R |
| **RPS29.3** | 2.83627344721428e-38 | -0.376878819205646 | 0.346 | 0.537 | 3.591005811518e-34 | 6 | RPS29 |
| **RPL10A.4** | 3.80842113639619e-38 | -0.370396273407814 | 0.362 | 0.563 | 4.82184200079121e-34 | 6 | RPL10A |
| **RPL9.4** | 3.92932063828388e-38 | -0.379870385019499 | 0.425 | 0.592 | 4.97491286013122e-34 | 6 | RPL9 |
| **SH3BGRL3.5** | 5.32067522457635e-37 | -0.329932098700539 | 0.177 | 0.409 | 6.73650690183612e-33 | 6 | SH3BGRL3 |
| **RPL4.3** | 6.78548292027502e-37 | -0.352471337491444 | 0.185 | 0.397 | 8.5910999253602e-33 | 6 | RPL4 |
| **C1QB.5** | 7.33874168154346e-37 | 0.380935523432671 | 0.718 | 0.413 | 9.29158084300218e-33 | 6 | C1QB |
| **RPS12.5** | 2.73464747130623e-36 | -0.390654509417676 | 0.39 | 0.547 | 3.46233716342082e-32 | 6 | RPS12 |
| **EEF1B2.2** | 2.83410660053845e-36 | -0.340281610056062 | 0.428 | 0.621 | 3.58826236694173e-32 | 6 | EEF1B2 |
| **RPS13.4** | 6.6283062739269e-36 | -0.365182084993769 | 0.461 | 0.622 | 8.39209857341885e-32 | 6 | RPS13 |
| **RPL5.2** | 7.10223002046718e-36 | -0.352993875766534 | 0.307 | 0.502 | 8.99213342891349e-32 | 6 | RPL5 |
| **RPS3.4** | 9.5812704840022e-36 | -0.367263508137651 | 0.353 | 0.532 | 1.21308465597952e-31 | 6 | RPS3 |
| **C1QC.4** | 1.44688719606688e-35 | 0.397225809526924 | 0.852 | 0.521 | 1.83190387894028e-31 | 6 | C1QC |
| **PABPC1.3** | 1.83287639481082e-35 | -0.336796145051232 | 0.396 | 0.603 | 2.32060480346998e-31 | 6 | PABPC1 |
| **ENSGALG00000015183.[illegible]** | 2.75836371549569e-35 | -0.329851165957734 | 0.1 | 0.314 | 3.4923643001891e-31 | 6 | ENSGALG00000015183 |
| **TSC22D4.6** | 2.9139290666732e-35 | -0.33491127904252 | 0.088 | 0.298 | 3.68932559131494e-31 | 6 | TSC22D4 |
| **FAU.4** | 5.14525318960747e-35 | -0.358512801197571 | 0.361 | 0.531 | 6.51440506336202e-31 | 6 | FAU |
| **ENSGALG00000004509.[illegible]** | 2.30164548650986e-34 | -0.351514826600551 | 0.808 | 0.871 | 2.91411335047014e-30 | 6 | ENSGALG00000004509 |
| **NFKBIA.5** | 3.70526679690445e-34 | -0.327539828642717 | 0.148 | 0.352 | 4.69123829156072e-30 | 6 | NFKBIA |
| **LAMP1.3** | 7.92807515259687e-34 | 0.259723479582136 | 0.548 | 0.323 | 1.00377359507029e-29 | 6 | LAMP1 |
| **ZEB2.6** | 9.34354086895836e-34 | -0.321938426608803 | 0.116 | 0.326 | 1.18298570941882e-29 | 6 | ZEB2 |
| **RPS21.4** | 3.10944861710062e-33 | -0.357739036332581 | 0.399 | 0.545 | 3.93687289411109e-29 | 6 | RPS21 |
| **RPL6.3** | 3.42712932111459e-32 | -0.333450180790449 | 0.414 | 0.578 | 4.33908843346319e-28 | 6 | RPL6 |
| **PNRC1.6** | 6.16509135672628e-32 | -0.343754828877452 | 0.404 | 0.555 | 7.80562216675114e-28 | 6 | PNRC1 |
| **NME2.3** | 1.03200035256299e-31 | -0.321150010758187 | 0.304 | 0.497 | 1.30661564638e-27 | 6 | NME2 |
| **RPL38.2** | 1.70042895082634e-31 | -0.314935783768845 | 0.374 | 0.55 | 2.15291309464123e-27 | 6 | RPL38 |
| **ENSGALG00000045199.[illegible]** | 7.68998337616295e-31 | -0.327420607313362 | 0.028 | 0.283 | 9.73628795255991e-27 | 6 | ENSGALG00000045199 |
| **HSPA5.5** | 4.78000519555069e-30 | -0.372524031596221 | 0.288 | 0.41 | 6.05196457808673e-26 | 6 | HSPA5 |
| **RPS2.4** | 3.62412823967754e-29 | -0.318561352506596 | 0.487 | 0.628 | 4.58850876425574e-25 | 6 | RPS2 |
| **FAM49A.3** | 6.32002063413429e-29 | -0.273277070554821 | 0.218 | 0.421 | 8.00177812487742e-25 | 6 | FAM49A |
| **DDX3X.2** | 9.0418461633288e-29 | -0.267455206197252 | 0.154 | 0.355 | 1.14478814273906e-24 | 6 | DDX3X |
| **ENSGALG00000047532.[illegible]** | 2.31073042591754e-28 | -0.260466174359095 | 0.171 | 0.378 | 2.9256157922542e-24 | 6 | ENSGALG00000047532 |
| **MYO9B.1** | 4.15709587426395e-28 | -0.256057025722966 | 0.155 | 0.374 | 5.26329908640559e-24 | 6 | MYO9B |
| **PTPRC.3** | 5.94512961471893e-28 | -0.266726497985742 | 0.197 | 0.394 | 7.52712860519564e-24 | 6 | PTPRC |
| **ENO1.2** | 1.42483415166779e-27 | -0.264718931126754 | 0.235 | 0.424 | 1.80398251942659e-23 | 6 | ENO1 |
| **MT-CO1.4** | 7.05105472064481e-27 | -0.315517799826184 | 0.982 | 0.976 | 8.9273403818084e-23 | 6 | MT-CO1 |
| **BTG1.5** | 1.55238888686158e-26 | -0.264601127914773 | 0.173 | 0.337 | 1.96547956965545e-22 | 6 | BTG1 |
| **DSTN.2** | 1.920299735899e-26 | -0.280133509426791 | 0.537 | 0.657 | 2.43129149562173e-22 | 6 | DSTN |
| **PPDPF.6** | 6.10822219416136e-26 | -0.470731534988421 | 0.374 | 0.36 | 7.73362012002769e-22 | 6 | PPDPF |
| **RPL3.4** | 6.99343178249382e-26 | -0.288050724713969 | 0.377 | 0.515 | 8.85438397981543e-22 | 6 | RPL3 |

| | | | | | | | |
|---|---|---|---|---|---|---|---|
| **CORO1C.2** | 1.96980742998158e-25 | -0.267204185855503 | 0.304 | 0.467 | 2.49397318709968e-21 | 6 | CORO1C |
| **AIF1L.1** | 3.34390863361367e-25 | -0.250185094363902 | 0.268 | 0.463 | 4.23372272101827e-21 | 6 | AIF1L |
| **H3F3B.2** | 9.84180659707644e-25 | -0.274623578123292 | 0.495 | 0.627 | 1.24607113325585e-20 | 6 | H3F3B |
| **ENSGALG00000014585.5** | 5.4165151646392e-24 | 0.331281367102289 | 0.674 | 0.358 | 6.85784984994969e-20 | 6 | ENSGALG00000014585 |
| **BLB2.5** | 2.07715041653528e-21 | -0.267577122954435 | 0.855 | 0.864 | 2.62988014237532e-17 | 6 | BLB2 |
| **COX3.4** | 8.43443412374728e-21 | -0.257021296749308 | 0.994 | 0.989 | 1.06788370440764e-16 | 6 | COX3 |
| **ATP6.4** | 2.9543552474947e-18 | -0.250536422659051 | 0.967 | 0.971 | 3.74050917885304e-14 | 6 | ATP6 |
| **TPT1.7** | 5.77504464882505e-290 | 1.10668936747795 | 0.994 | 0.784 | 7.3117840298774e-286 | 7 | TPT1 |
| **RPS11.6** | 7.77002117281973e-222 | 1.11151264065136 | 0.995 | 0.768 | 9.83762380690706e-218 | 7 | RPS11 |
| **RPS27A.6** | 4.26376421644502e-201 | 0.909596348466133 | 0.982 | 0.68 | 5.39835187444104e-197 | 7 | RPS27A |
| **EEF2.5** | 5.5275204121656e-201 | 0.846159770668087 | 0.951 | 0.486 | 6.99839359384287e-197 | 7 | EEF2 |
| **RPS3A.6** | 1.803674304694e-197 | 0.877003138465023 | 0.986 | 0.697 | 2.28363203717307e-193 | 7 | RPS3A |
| **RPL19.6** | 2.44063012529531e-196 | 0.839588523008013 | 0.971 | 0.622 | 3.09008180163639e-192 | 7 | RPL19 |
| **RPS16.6** | 2.32173795771877e-194 | 0.850635036739419 | 0.959 | 0.635 | 2.93955242826773e-190 | 7 | RPS16 |
| **ENSGALG00000026970.4** | 2.38559786352477e-194 | 0.854787735012081 | 0.951 | 0.503 | 3.02040545500871e-190 | 7 | ENSGALG00000026970 |
| **RPS23.5** | 2.99771443973395e-192 | 0.861917214053525 | 0.982 | 0.679 | 3.79540625214715e-188 | 7 | RPS23 |
| **EEF1A1.5** | 8.78398086551444e-191 | 0.815014149520975 | 0.992 | 0.804 | 1.11213981738278e-186 | 7 | EEF1A1 |
| **RPS24.6** | 1.05951892790208e-189 | 0.82565003676647 | 0.977 | 0.657 | 1.34145691461682e-185 | 7 | RPS24 |
| **RPL23.6** | 4.45685957432345e-188 | 0.852186308715313 | 0.991 | 0.781 | 5.64282990705092e-184 | 7 | RPL23 |
| **RPS14.6** | 1.63923813526895e-187 | 0.889918130852202 | 0.989 | 0.784 | 2.07543940306401e-183 | 7 | RPS14 |
| **RPL15.6** | 2.65435829893832e-186 | 0.830962281985055 | 0.983 | 0.754 | 3.36068304228581e-182 | 7 | RPL15 |
| **RPS15A.6** | 6.85471945521696e-185 | 0.839828264383105 | 0.992 | 0.763 | 8.67876030225019e-181 | 7 | RPS15A |
| **RPS19.6** | 5.72883293742827e-183 | 0.907443972712466 | 0.998 | 0.837 | 7.25327538207793e-179 | 7 | RPS19 |
| **RPS17.6** | 1.19634347117579e-182 | 0.836597635663393 | 0.997 | 0.811 | 1.51469046885567e-178 | 7 | RPS17 |
| **RPL35A.6** | 4.14828760733737e-182 | 0.837567978747884 | 0.976 | 0.717 | 5.25214693964984e-178 | 7 | RPL35A |
| **RPLP2.6** | 8.65718076737476e-182 | 0.931594068487523 | 0.989 | 0.757 | 1.09608565695732e-177 | 7 | RPLP2 |
| **RPL26L1.6** | 1.85234241588568e-180 | 0.862835207303259 | 0.974 | 0.612 | 2.34525073275286e-176 | 7 | RPL26L1 |
| **RPL24.6** | 6.22580021802062e-180 | 0.835775949709075 | 0.977 | 0.66 | 7.88248565603591e-176 | 7 | RPL24 |
| **RPS8.6** | 9.24445243611208e-178 | 0.802314064720326 | 0.991 | 0.808 | 1.17044012293615e-173 | 7 | RPS8 |
| **RPS13.5** | 5.14439410077779e-176 | 0.730124317788277 | 0.942 | 0.582 | 6.51331737099476e-172 | 7 | RPS13 |
| **ENSGALG00000037441.6** | 7.13137022960757e-175 | 0.872339547931024 | 0.986 | 0.761 | 9.02902784770614e-171 | 7 | ENSGALG00000037441 |
| **RPS10.6** | 9.93815494498568e-175 | 0.839162963040431 | 0.97 | 0.615 | 1.25826979758464e-170 | 7 | RPS10 |
| **RPL32.6** | 1.50002195483646e-173 | 0.770427350075142 | 0.967 | 0.643 | 1.89917779701844e-169 | 7 | RPL32 |
| **ITM2A.5** | 7.32041498752226e-173 | 0.765065758350818 | 0.947 | 0.508 | 9.26837741570194e-169 | 7 | ITM2A |
| **RPS7.4** | 2.09299335807612e-172 | 0.744330974022737 | 0.925 | 0.529 | 2.64993889066018e-168 | 7 | RPS7 |
| **RPL37.6** | 5.29702617417763e-172 | 0.842492219957849 | 0.983 | 0.688 | 6.7065648391263e-168 | 7 | RPL37 |
| **RPL27.4** | 1.20706527473318e-166 | 0.734009212619289 | 0.918 | 0.504 | 1.52826534433968e-162 | 7 | RPL27 |
| **RPL18A.5** | 2.35665628779077e-166 | 0.780366396669821 | 0.956 | 0.534 | 2.98376252597189e-162 | 7 | RPL18A |
| **RPS26.6** | 5.79989585105525e-165 | 0.836564958169822 | 0.983 | 0.688 | 7.34324813702105e-161 | 7 | RPS26 |
| **RPL21.6** | 4.42316556955122e-164 | 0.772702952084939 | 0.991 | 0.815 | 5.6001699276088e-160 | 7 | RPL21 |
| **RPL7A.6** | 3.80252771618094e-163 | 0.792183112874898 | 0.933 | 0.56 | 4.81438034145669e-159 | 7 | RPL7A |
| **RPL17.6** | 2.83090526944287e-162 | 0.758987976500127 | 0.971 | 0.732 | 3.58420916164162e-158 | 7 | RPL17 |
| **RPS4Y1.6** | 5.17507146222078e-162 | 0.756650419855122 | 0.927 | 0.499 | 6.55215797831773e-158 | 7 | RPS4Y1 |
| **GNB2L1.6** | 6.02869755081228e-162 | 0.839340136184073 | 0.965 | 0.602 | 7.63293396908343e-158 | 7 | GNB2L1 |
| **RPS20.6** | 1.34053730232602e-159 | 0.740050606284724 | 0.979 | 0.732 | 1.69725427847497e-155 | 7 | RPS20 |
| **RPL34.6** | 2.47043508077924e-159 | 0.811790582795045 | 0.989 | 0.704 | 3.1278178557746e-155 | 7 | RPL34 |
| **RPL23A.5** | 3.67459266470251e-159 | 0.772925891675137 | 0.96 | 0.584 | 4.65240177277984e-155 | 7 | RPL23A |
| **FAU.5** | 5.59220989400284e-159 | 0.714527207139517 | 0.91 | 0.485 | 7.08029694679699e-155 | 7 | FAU |
| **RPL27A.6** | 1.21526204658374e-158 | 0.867097392063034 | 0.979 | 0.644 | 1.53864327717967e-154 | 7 | RPL27A |
| **PABPC1.4** | 2.25294113194867e-158 | 0.613179987454173 | 0.927 | 0.559 | 2.85244876716021e-154 | 7 | PABPC1 |
| **EEF1D.6** | 2.9136055531379e-158 | 0.868452617242258 | 0.921 | 0.423 | 3.68891599082789e-154 | 7 | EEF1D |
| **RPL36AL.6** | 3.7918146581993e-158 | 0.724332308561666 | 0.95 | 0.597 | 4.80081653874613e-154 | 7 | RPL36AL |
| **RPL11.5** | 2.61091542823619e-156 | 0.714475294744556 | 0.947 | 0.631 | 3.30568002368984e-152 | 7 | RPL11 |
| **RPL35.5** | 2.29050975460805e-155 | 0.715043151108444 | 0.947 | 0.593 | 2.90001440030926e-151 | 7 | RPL35 |

| | | | | | | | |
|---|---|---|---|---|---|---|---|
| **RPS6.5** | 4.05522222503471e-153 | 0.729018345012622 | 0.954 | 0.598 | 5.13431685911644e-149 | 7 | RPS6 |
| **RPL31.5** | 6.37739958454514e-153 | 0.725940087133243 | 0.944 | 0.582 | 8.0744256139926e-149 | 7 | RPL31 |
| **DUSP1.2** | 8.6494039288392e-153 | 0.744832848602115 | 0.741 | 0.15 | 1.09510103143033e-148 | 7 | DUSP1 |
| **RPL37A.6** | 1.03252970663678e-150 | 0.777822143038132 | 0.989 | 0.73 | 1.30728586157282e-146 | 7 | RPL37A |
| **UBA52.5** | 9.62447586045477e-150 | 0.695992958734158 | 0.941 | 0.541 | 1.21855488869218e-145 | 7 | UBA52 |
| **RPL8.6** | 1.32597873353378e-149 | 0.765931316176662 | 0.96 | 0.607 | 1.67882167452712e-145 | 7 | RPL8 |
| **RPS21.5** | 7.13778851667666e-149 | 0.722437439231352 | 0.915 | 0.502 | 9.03715404096432e-145 | 7 | RPS21 |
| **CG-1B.1** | 1.16131220085011e-147 | 0.770269242925392 | 0.72 | 0.132 | 1.47033737749632e-143 | 7 | CG-1B |
| **RPS3.5** | 1.31232624920823e-147 | 0.709202887339694 | 0.921 | 0.485 | 1.66153626412254e-143 | 7 | RPS3 |
| **RPS25.5** | 4.71611804391388e-147 | 0.719314224245142 | 0.945 | 0.607 | 5.97107705539937e-143 | 7 | RPS25 |
| **RPL13.6** | 4.72677492284103e-146 | 0.742727972577856 | 0.962 | 0.602 | 5.98456972980902e-142 | 7 | RPL13 |
| **RPL7.5** | 2.31463368752377e-145 | 0.738797272836094 | 0.944 | 0.56 | 2.93055771177385e-141 | 7 | RPL7 |
| **RPL29.4** | 3.9889946323294e-144 | 0.666329200427521 | 0.939 | 0.6 | 5.05046610399226e-140 | 7 | RPL29 |
| **RPL30.5** | 1.38071670801114e-143 | 0.721883519521944 | 0.928 | 0.547 | 1.74812542401291e-139 | 7 | RPL30 |
| **RPL5.3** | 7.87297202695606e-143 | 0.657131194107006 | 0.875 | 0.454 | 9.96796988332907e-139 | 7 | RPL5 |
| **RPS15.6** | 1.38416471174123e-139 | 0.786848703771118 | 0.995 | 0.752 | 1.75249094153558e-135 | 7 | RPS15 |
| **CST3.4** | 1.92117535865902e-138 | 0.611927095175779 | 0.96 | 0.69 | 2.43240012159818e-134 | 7 | CST3 |
| **ASS1.4** | 2.29510720984303e-136 | 0.697527925859425 | 0.846 | 0.337 | 2.90583523838226e-132 | 7 | ASS1 |
| **RPL9.5** | 1.46355614807983e-135 | 0.66790015311615 | 0.919 | 0.551 | 1.85300843908388e-131 | 7 | RPL9 |
| **RPL3.5** | 2.49801570127041e-134 | 0.61492819950823 | 0.857 | 0.475 | 3.16273767937847e-130 | 7 | RPL3 |
| **LGALS3** | 3.93181040559422e-132 | 0.348925821146811 | 0.423 | 0.09 | 4.97806515452284e-128 | 7 | LGALS3 |
| **RPL22.5** | 1.46710071721087e-131 | 0.653077284623033 | 0.936 | 0.569 | 1.85749621806068e-127 | 7 | RPL22 |
| **RPS28.6** | 1.79703290494628e-131 | 0.70589270100275 | 0.954 | 0.575 | 2.27522336095248e-127 | 7 | RPS28 |
| **CD81.4** | 4.59806610654416e-131 | 0.600391191597414 | 0.74 | 0.222 | 5.82161149749556e-127 | 7 | CD81 |
| **RPL12.5** | 5.89998653874158e-131 | 0.692869900863263 | 0.95 | 0.571 | 7.46997295670072e-127 | 7 | RPL12 |
| **MT-CO1.5** | 4.08283956030408e-130 | 0.677237857466653 | 1 | 0.975 | 5.16928316730099e-126 | 7 | MT-CO1 |
| **H3F3C.1** | 4.59974406843423e-129 | 0.521423282649836 | 0.839 | 0.48 | 5.82373596504457e-125 | 7 | H3F3C |
| **RPS12.6** | 1.96052462155651e-128 | 0.724787658366092 | 0.916 | 0.504 | 2.4822202233527e-124 | 7 | RPS12 |
| **B2M.3** | 3.95936302708799e-128 | 0.51656382212397 | 0.997 | 0.926 | 5.01294952859611e-124 | 7 | B2M |
| **LDHA.2** | 1.98866464600932e-127 | 0.95977119341745 | 0.88 | 0.162 | 2.5178483083124e-123 | 7 | LDHA |
| **RPL36.6** | 3.18751524967432e-127 | 0.686566718823734 | 0.935 | 0.503 | 4.03571305761265e-123 | 7 | RPL36 |
| **COX3.5** | 5.24571164971741e-127 | 0.626953547390015 | 1 | 0.988 | 6.64159551970721e-123 | 7 | COX3 |
| **RPL39L.4** | 5.3496444669682e-127 | 0.610371379384166 | 0.989 | 0.817 | 6.77318485962844e-123 | 7 | RPL39L |
| **SLC25A6.3** | 7.82276013242445e-127 | 0.552844613776494 | 0.753 | 0.302 | 9.9043966036626e-123 | 7 | SLC25A6 |
| **CTSZ.4** | 1.90273511565507e-126 | 0.551013555307882 | 0.773 | 0.347 | 2.40905292993088e-122 | 7 | CTSZ |
| **FOS.7** | 2.30106109853267e-124 | 0.931760926028405 | 0.915 | 0.35 | 2.91337345685222e-120 | 7 | FOS |
| **LSP1P1.4** | 4.92721024564111e-124 | 0.626493246227862 | 0.814 | 0.333 | 6.2383408920062e-120 | 7 | LSP1P1 |
| **ENSGALG00000008518.1** | 6.00284936174988e-123 | 0.489836881769489 | 0.568 | 0.128 | 7.60020757691152e-119 | 7 | ENSGALG00000008518 |
| **RPL10A.5** | 3.13228242919796e-122 | 0.615562296024568 | 0.893 | 0.519 | 3.96578278360754e-118 | 7 | RPL10A |
| **ATP6V0C.6** | 1.49916750201039e-121 | 0.666610623297896 | 0.918 | 0.526 | 1.89809597429535e-117 | 7 | ATP6V0C |
| **MGST1.3** | 2.24551099143108e-119 | 0.853378221842444 | 0.863 | 0.177 | 2.84304146625089e-115 | 7 | MGST1 |
| **ATP6.5** | 1.38700356339487e-118 | 0.630769270880473 | 0.998 | 0.968 | 1.75608521161424e-114 | 7 | ATP6 |
| **CD99.1** | 2.17229866182535e-117 | 0.563484812257952 | 0.565 | 0.078 | 2.75034733573708e-113 | 7 | CD99 |
| **PPDPF.7** | 6.56742814445481e-114 | 1.20335019625763 | 0.977 | 0.311 | 8.31502077369423e-110 | 7 | PPDPF |
| **RPL4.4** | 2.02765388263614e-113 | 0.617995375307915 | 0.83 | 0.344 | 2.56721258080561e-109 | 7 | RPL4 |
| **NME2.4** | 1.68854883335429e-112 | 0.569433501644133 | 0.855 | 0.452 | 2.13787167790987e-108 | 7 | NME2 |
| **RPL6.4** | 2.88566772216069e-111 | 0.577194458185314 | 0.878 | 0.539 | 3.65354390302765e-107 | 7 | RPL6 |
| **TXN.4** | 2.32852671299377e-110 | 0.63047847747794 | 0.861 | 0.462 | 2.94814767132141e-106 | 7 | TXN |
| **RPL14.3** | 2.64175102241898e-110 | 0.574184185532494 | 0.884 | 0.533 | 3.34472096948467e-106 | 7 | RPL14 |
| **NFE2L2.1** | 1.12734716763775e-109 | 0.37375060463281 | 0.478 | 0.133 | 1.42733424894616e-105 | 7 | NFE2L2 |
| **SPTBN1.1** | 2.72402660945695e-108 | 0.532554991669387 | 0.549 | 0.087 | 3.44889009023344e-104 | 7 | SPTBN1 |
| **ZNF706.1** | 1.02931574940661e-107 | 0.405124452223031 | 0.638 | 0.29 | 1.30321667032371e-103 | 7 | ZNF706 |
| **PRCP.1** | 1.6352631911255e-105 | 0.386683439045525 | 0.466 | 0.112 | 2.07040672628399e-101 | 7 | PRCP |
| **ITGB1BP3.4** | 1.79053137991129e-105 | 0.602543621652827 | 0.916 | 0.503 | 2.26699178010569e-101 | 7 | ITGB1BP3 |

| | | | | | | | |
|---|---|---|---|---|---|---|---|
| **CXCL13.7** | 4.44231242239362e-105 | -1.41412722553092 | 0.307 | 0.431 | 5.62441175799257e-101 | 7 | CXCL13 |
| **ANXA5.1** | 1.44544934714419e-103 | 0.369119432889719 | 0.478 | 0.116 | 1.83008341841926e-99 | 7 | ANXA5 |
| **RPS2.5** | 5.39916428971978e-103 | 0.560640794739883 | 0.915 | 0.592 | 6.83588190721421e-99 | 7 | RPS2 |
| **CDK5RAP1.1** | 7.11096921243541e-102 | 0.544198296277185 | 0.654 | 0.162 | 9.00319811986447e-98 | 7 | CDK5RAP1 |
| **HNRNPA1.1** | 1.01836236383703e-101 | 0.420025928203083 | 0.61 | 0.229 | 1.28934858885406e-97 | 7 | HNRNPA1 |
| **PRNP.1** | 1.12453031064415e-101 | 0.368877509500838 | 0.463 | 0.119 | 1.42376782630656e-97 | 7 | PRNP |
| **BTG1.6** | 1.2589185211372e-101 | 0.515365277146524 | 0.728 | 0.291 | 1.59391673961181e-97 | 7 | BTG1 |
| **CORO1C.3** | 1.30052600203547e-101 | 0.496059665654553 | 0.798 | 0.426 | 1.64659597117711e-97 | 7 | CORO1C |
| **RPSAP58.3** | 1.74096223707e-101 | 0.544754629475531 | 0.89 | 0.558 | 2.20423228835432e-97 | 7 | RPSAP58 |
| **BZW2.1** | 2.21066854846132e-98 | 0.331620408367596 | 0.419 | 0.108 | 2.79892744920688e-94 | 7 | BZW2 |
| **TLR7.1** | 1.70574349027604e-97 | 0.315894719749729 | 0.394 | 0.079 | 2.1596418330385e-93 | 7 | TLR7 |
| **RABGAP1.1** | 1.79715918629001e-96 | 0.296285420094172 | 0.362 | 0.07 | 2.27538324576179e-92 | 7 | RABGAP1 |
| **FTH1.6** | 4.35528222722107e-96 | 0.623157179520208 | 1 | 0.99 | 5.5142228278846e-92 | 7 | FTH1 |
| **RPLP1.6** | 1.99378773817869e-95 | 0.623964854476788 | 0.995 | 0.898 | 2.52433465530804e-91 | 7 | RPLP1 |
| **FGL2.3** | 4.84734376151729e-95 | 0.45558732680217 | 0.629 | 0.207 | 6.13722193645704e-91 | 7 | FGL2 |
| **GNAI3.1** | 1.07451542254804e-94 | 0.393723939850481 | 0.553 | 0.181 | 1.36044397648807e-90 | 7 | GNAI3 |
| **HSPA8.6** | 1.07506695795304e-94 | 0.66503907568511 | 0.942 | 0.597 | 1.36114227546434e-90 | 7 | HSPA8 |
| **RASSF2.2** | 4.81067760148536e-94 | 0.443663576241346 | 0.686 | 0.295 | 6.09079891124061e-90 | 7 | RASSF2 |
| **COX4I1.1** | 1.19328750372163e-93 | 0.427056592856165 | 0.691 | 0.356 | 1.51082130846196e-89 | 7 | COX4I1 |
| **EEF1B2.3** | 2.49140125249689e-93 | 0.486872299408153 | 0.858 | 0.585 | 3.15436312578631e-89 | 7 | EEF1B2 |
| **AHR** | 4.98682910662884e-93 | 0.307332408012291 | 0.387 | 0.09 | 6.31382433190278e-89 | 7 | AHR |
| **CALM2.2** | 4.34925480048664e-92 | 0.460815907004585 | 0.752 | 0.396 | 5.50659150289613e-88 | 7 | CALM2 |
| **COII.4** | 4.46195641148314e-92 | 0.530791960151759 | 1 | 0.984 | 5.6492830125788e-88 | 7 | COII |
| **H3F3B.3** | 1.09062396790994e-91 | 0.484293335887547 | 0.898 | 0.593 | 1.38083900577077e-87 | 7 | H3F3B |
| **LTA4H.1** | 6.423261255763e-91 | 0.431118408984649 | 0.46 | 0.092 | 8.13249107592154e-87 | 7 | LTA4H |
| **CD74.5** | 2.25647510971283e-90 | 0.543606065479576 | 1 | 0.994 | 2.85692313640741e-86 | 7 | CD74 |
| **GNAI2.2** | 4.10890152132509e-89 | 0.411766867711927 | 0.636 | 0.249 | 5.2022802161497e-85 | 7 | GNAI2 |
| **RAMP2.1** | 4.77435926725651e-89 | 0.364765137995161 | 0.393 | 0.053 | 6.04481626827347e-85 | 7 | RAMP2 |
| **VDAC1** | 5.73472548729524e-89 | 0.263788779742961 | 0.333 | 0.065 | 7.2607359394645e-85 | 7 | VDAC1 |
| **S100A11.1** | 7.61698637852939e-89 | 0.475619073266979 | 0.578 | 0.159 | 9.64386645385606e-85 | 7 | S100A11 |
| **GAPDH.5** | 1.47800238525137e-88 | 0.577397983391329 | 0.837 | 0.499 | 1.87129881996676e-84 | 7 | GAPDH |
| **DSTN.3** | 2.79258277902035e-88 | 0.460956111159232 | 0.898 | 0.627 | 3.53568905651766e-84 | 7 | DSTN |
| **ARF6.1** | 2.24650567579372e-87 | 0.376172428189729 | 0.56 | 0.233 | 2.84430083612243e-83 | 7 | ARF6 |
| **EIF3M.1** | 5.02279047789732e-87 | 0.354902614462489 | 0.511 | 0.184 | 6.35935502406579e-83 | 7 | EIF3M |
| **VIM.2** | 5.27606078849165e-87 | 0.906914795447042 | 0.839 | 0.161 | 6.68002056430928e-83 | 7 | VIM |
| **HMGB1.2** | 5.94295480099826e-87 | 0.488770564861315 | 0.715 | 0.309 | 7.5243750735439e-83 | 7 | HMGB1 |
| **S100A10.7** | 6.42299888750138e-87 | 1.01193393723188 | 0.954 | 0.339 | 8.13215889146549e-83 | 7 | S100A10 |
| **RPL38.3** | 8.85182454293513e-86 | 0.479833815096106 | 0.846 | 0.511 | 1.12072950538102e-81 | 7 | RPL38 |
| **CSF3R.1** | 1.83930162279128e-85 | 0.330737150193338 | 0.399 | 0.063 | 2.32873978461605e-81 | 7 | CSF3R |
| **SHFM1.1** | 5.93113296532867e-85 | 0.345280458432726 | 0.53 | 0.194 | 7.50940744740263e-81 | 7 | SHFM1 |
| **CRIP1.6** | 6.46071576615385e-85 | 1.03708003079292 | 0.871 | 0.188 | 8.17991223152739e-81 | 7 | CRIP1 |
| **GATD3AL1.1** | 2.4295071086427e-84 | 0.366322099237246 | 0.534 | 0.171 | 3.07599895025252e-80 | 7 | GATD3AL1 |
| **MECR.1** | 2.67864058067521e-84 | 0.348556109091409 | 0.444 | 0.129 | 3.39142683919288e-80 | 7 | MECR |
| **APLP2.1** | 5.95291330138459e-84 | 0.372757650297278 | 0.546 | 0.209 | 7.53698353088303e-80 | 7 | APLP2 |
| **GNG5.2** | 6.92342010203571e-84 | 0.425079477636395 | 0.788 | 0.468 | 8.76574219118741e-80 | 7 | GNG5 |
| **EIF3F.1** | 9.0909663386727e-84 | 0.391160424302375 | 0.518 | 0.151 | 1.15100724813935e-79 | 7 | EIF3F |
| **RPS29.4** | 1.39487010359175e-83 | 0.539606585164428 | 0.851 | 0.495 | 1.76604503815752e-79 | 7 | RPS29 |
| **CALM1.1** | 4.93072979011233e-83 | 0.412905545959527 | 0.784 | 0.502 | 6.24279698726122e-79 | 7 | CALM1 |
| **BLB2.6** | 5.83440250531183e-83 | 0.504976675048069 | 0.995 | 0.853 | 7.3869370119753e-79 | 7 | BLB2 |
| **ND3.4** | 3.19789673022914e-82 | 0.554140217512975 | 0.995 | 0.808 | 4.04885705014312e-78 | 7 | ND3 |
| **BSG** | 5.60102539534957e-82 | 0.259956516843399 | 0.347 | 0.085 | 7.09145825305209e-78 | 7 | BSG |
| **RPLP0.1** | 4.1920601335448e-81 | 0.434105883118385 | 0.679 | 0.235 | 5.30756733508107e-77 | 7 | RPLP0 |
| **INIP** | 5.30121380539571e-81 | 0.262251460601959 | 0.344 | 0.085 | 6.7118667990115e-77 | 7 | INIP |
| **COX7B.1** | 1.72694002313185e-80 | 0.363881640837661 | 0.632 | 0.331 | 2.18647876328723e-76 | 7 | COX7B |

| | | | | | | | |
|---|---|---|---|---|---|---|---|
| **EMP1.1** | 2.78812718388535e-80 | 0.471153436510139 | 0.49 | 0.069 | 3.53004782751724e-76 | 7 | EMP1 |
| **KLF2.1** | 4.12873695756901e-80 | 0.332387413881935 | 0.37 | 0.068 | 5.22739386197812e-76 | 7 | KLF2 |
| **ANKRD11.5** | 1.89373938244575e-79 | 0.488235533019911 | 0.72 | 0.299 | 2.39766343211457e-75 | 7 | ANKRD11 |
| **MXD1** | 2.83563345457286e-79 | 0.266640224239517 | 0.336 | 0.069 | 3.5901955168347e-75 | 7 | MXD1 |
| **EIF3E.1** | 2.97820346677681e-79 | 0.349095610875279 | 0.546 | 0.242 | 3.77070340928612e-75 | 7 | EIF3E |
| **PRDX6.1** | 1.19520574429423e-78 | 0.300843765800479 | 0.405 | 0.126 | 1.51324999285092e-74 | 7 | PRDX6 |
| **LY86.6** | 1.84432800341866e-78 | 0.833249140838794 | 0.839 | 0.195 | 2.33510368512837e-74 | 7 | LY86 |
| **MARCO.7** | 1.22353636462618e-77 | -0.817698324298116 | 0.093 | 0.404 | 1.5491193912532e-73 | 7 | MARCO |
| **YWHAQ.1** | 1.86219219221614e-77 | 0.348944627701648 | 0.536 | 0.24 | 2.35772153456486e-73 | 7 | YWHAQ |
| **TPM3.1** | 5.78840297316043e-77 | 0.398180000915493 | 0.583 | 0.217 | 7.32869700431842e-73 | 7 | TPM3 |
| **GPX1.1** | 1.01245517562131e-76 | 0.475563286003338 | 0.606 | 0.146 | 1.28186949785414e-72 | 7 | GPX1 |
| **NDUFS8.1** | 1.64386988503932e-76 | 0.295891054988895 | 0.449 | 0.147 | 2.08130366144828e-72 | 7 | NDUFS8 |
| **ZEB2.7** | 2.77520991491964e-76 | 0.510806720269945 | 0.699 | 0.277 | 3.51369327327976e-72 | 7 | ZEB2 |
| **CTSD.6** | 8.18542976643662e-76 | -0.771093698986105 | 0.245 | 0.508 | 1.03635726272854e-71 | 7 | CTSD |
| **TGFBI.1** | 1.22446539595242e-74 | 0.422418178109519 | 0.633 | 0.293 | 1.55029563781536e-70 | 7 | TGFBI |
| **SLC1A6.1** | 2.05268099832777e-74 | 0.293629140163819 | 0.329 | 0.049 | 2.59889941198279e-70 | 7 | SLC1A6 |
| **FABP3.1** | 4.8211872313145e-74 | 0.736569473258457 | 0.699 | 0.1 | 6.10410515356729e-70 | 7 | FABP3 |
| **PGD.1** | 6.49670009881798e-74 | 0.277192899034687 | 0.368 | 0.089 | 8.22547199511344e-70 | 7 | PGD |
| **ANXA2.1** | 8.46838164775127e-74 | 0.696841167427455 | 0.747 | 0.111 | 1.07218180042179e-69 | 7 | ANXA2 |
| **TSC22D4.7** | 8.50324949767001e-74 | 0.525775055261471 | 0.729 | 0.245 | 1.0765964189e-69 | 7 | TSC22D4 |
| **RHOA.1** | 3.16226749119975e-73 | 0.38568457394557 | 0.752 | 0.464 | 4.003746870608e-69 | 7 | RHOA |
| **CDKN1B.1** | 4.40404351777046e-73 | 0.34508231369256 | 0.473 | 0.133 | 5.57595949784918e-69 | 7 | CDKN1B |
| **MMR1L4.1** | 1.09967770757218e-72 | 0.291371126005361 | 0.355 | 0.092 | 1.39230194555713e-68 | 7 | MMR1L4 |
| **SLC38A2.1** | 2.92967919016377e-72 | 0.284157403773933 | 0.377 | 0.09 | 3.70926682266635e-68 | 7 | SLC38A2 |
| **TGM4.7** | 3.2178579412246e-72 | -0.811711911862368 | 0.099 | 0.356 | 4.07412993938447e-68 | 7 | TGM4 |
| **SPEN.1** | 2.19025301288186e-71 | 0.360510956840952 | 0.524 | 0.213 | 2.77307933960972e-67 | 7 | SPEN |
| **COTL1.1** | 2.846262352364e-71 | 0.355048461004487 | 0.571 | 0.256 | 3.60365276432806e-67 | 7 | COTL1 |
| **IFI30.5** | 7.61368987123924e-71 | 0.602094709625814 | 0.998 | 0.811 | 9.63969274597601e-67 | 7 | IFI30 |
| **TLR1B.1** | 8.02709654367315e-71 | 0.289758528686814 | 0.371 | 0.101 | 1.01631069339446e-66 | 7 | TLR1B |
| **CD44** | 1.68352167333368e-70 | 0.283865965280241 | 0.388 | 0.1 | 2.13150679060777e-66 | 7 | CD44 |
| **SMAP2.2** | 2.00658334159871e-70 | 0.361471424301538 | 0.665 | 0.336 | 2.54053516879813e-66 | 7 | SMAP2 |
| **FN1.2** | 4.70356814567138e-70 | 1.35304583827233 | 0.979 | 0.142 | 5.95518762923454e-66 | 7 | FN1 |
| **S100A6.7** | 9.62400213660721e-70 | 1.20791818594112 | 0.974 | 0.231 | 1.21849491051584e-65 | 7 | S100A6 |
| **GSTA3.4** | 2.10372553184544e-69 | 0.400651170584763 | 0.667 | 0.275 | 2.66352689586951e-65 | 7 | GSTA3 |
| **RAB1B.2** | 4.89551996370228e-69 | 0.349488656921307 | 0.63 | 0.31 | 6.19821782604346e-65 | 7 | RAB1B |
| **AP1S2.1** | 5.32680619881605e-69 | 0.330342677777304 | 0.495 | 0.196 | 6.744269328321e-65 | 7 | AP1S2 |
| **URM1.1** | 7.11118184565123e-69 | 0.377498405767143 | 0.566 | 0.212 | 9.00346733477902e-65 | 7 | URM1 |
| **ZNFX1** | 8.15519681093323e-69 | 0.359683283335021 | 0.496 | 0.196 | 1.03252946823226e-64 | 7 | ZNFX1 |
| **RCSD1.1** | 9.73219982142066e-69 | 0.335451610065854 | 0.451 | 0.151 | 1.23219381939007e-64 | 7 | RCSD1 |
| **EIF3H.1** | 1.24851631896636e-68 | 0.319536088719944 | 0.498 | 0.21 | 1.58074651144331e-64 | 7 | EIF3H |
| **HSP90AA1.6** | 1.69019436502035e-68 | 0.614140952245632 | 0.98 | 0.771 | 2.13995508555226e-64 | 7 | HSP90AA1 |
| **MSN.1** | 1.97420253397917e-68 | 0.36587549008318 | 0.612 | 0.308 | 2.49953782827103e-64 | 7 | MSN |
| **SPI1.2** | 2.95944864497057e-68 | 0.351787762986112 | 0.613 | 0.288 | 3.74695792939724e-64 | 7 | SPI1 |
| **DDX5.6** | 4.33894067792992e-68 | 0.49529156704357 | 0.863 | 0.491 | 5.49353279232707e-64 | 7 | DDX5 |
| **VCAN.1** | 3.07805097082338e-67 | 0.957553036943459 | 0.851 | 0.122 | 3.89712033415947e-63 | 7 | VCAN |
| **DBI.1** | 3.96683590249218e-67 | 0.313571526507814 | 0.478 | 0.189 | 5.02241093614534e-63 | 7 | DBI |
| **ZNF593.1** | 1.54340890036625e-66 | 0.347139430757931 | 0.609 | 0.292 | 1.95411000875371e-62 | 7 | ZNF593 |
| **BRI3.3** | 3.37583538290982e-66 | 0.359333973867687 | 0.688 | 0.35 | 4.27414517830213e-62 | 7 | BRI3 |
| **C15orf48.1** | 1.41651426747285e-65 | 0.303940215331925 | 0.379 | 0.11 | 1.79344871404737e-61 | 7 | C15orf48 |
| **AIF1L.2** | 2.47626018260132e-65 | 0.36287364832491 | 0.706 | 0.426 | 3.13519301719153e-61 | 7 | AIF1L |
| **SLC40A1.7** | 4.63139666846865e-65 | -0.585612127711487 | 0.076 | 0.451 | 5.86381132194816e-61 | 7 | SLC40A1 |
| **CTSA.4** | 1.52014916921065e-64 | 0.4047627449349 | 0.691 | 0.336 | 1.9246608631376e-60 | 7 | CTSA |
| **PNRC1.7** | 1.89644563284479e-64 | 0.477157568222676 | 0.884 | 0.515 | 2.40108981574479e-60 | 7 | PNRC1 |
| **TOP2B.1** | 2.36183147454737e-64 | 0.285109069826656 | 0.44 | 0.159 | 2.99031482992443e-60 | 7 | TOP2B |

| | | | | | | | |
|---|---|---|---|---|---|---|---|
| **ENSGALG00000011190.1** | 2.46006311924146e-64 | -0.648583137691789 | 0.143 | 0.414 | 3.11468591527161e-60 | 7 | ENSGALG00000011190 |
| **AVD.1** | 3.03677502024691e-64 | 0.699048817037063 | 0.385 | 0.062 | 3.84486085313461e-60 | 7 | AVD |
| **EXFABP.7** | 3.82051572436883e-63 | 1.86748391001521 | 0.979 | 0.173 | 4.83715495862338e-59 | 7 | EXFABP |
| **ENSGALG00000054657.1** | 4.40647763503956e-63 | 0.28443318390763 | 0.358 | 0.057 | 5.57904133372358e-59 | 7 | ENSGALG00000054657 |
| **HSPA5.6** | 9.15225453446675e-62 | 0.606324932963007 | 0.791 | 0.369 | 1.15876694660883e-57 | 7 | HSPA5 |
| **EMB.1** | 1.51906442500616e-61 | 0.260266763008478 | 0.307 | 0.052 | 1.9232874685003e-57 | 7 | EMB |
| **SH3BGRL.1** | 3.05259113404228e-61 | 0.328251413561078 | 0.458 | 0.157 | 3.86488563481093e-57 | 7 | SH3BGRL |
| **BTG2.6** | 5.20749078005567e-61 | 0.532219905491949 | 0.732 | 0.281 | 6.59320407662848e-57 | 7 | BTG2 |
| **ANXA1.2** | 5.51277755041252e-61 | 1.16361212239924 | 0.941 | 0.142 | 6.9797276565773e-57 | 7 | ANXA1 |
| **BNIP3.1** | 7.25124188237856e-61 | 0.251558604767648 | 0.339 | 0.091 | 9.1807973472795e-57 | 7 | BNIP3 |
| **CYBA.1** | 2.28659641005743e-60 | 0.356649883161113 | 0.624 | 0.325 | 2.89505971477371e-56 | 7 | CYBA |
| **SPCS3.1** | 1.36463501052764e-59 | 0.289593098556674 | 0.479 | 0.209 | 1.72776438682904e-55 | 7 | SPCS3 |
| **TPI1.1** | 1.73534209130773e-59 | 0.276308128516598 | 0.434 | 0.174 | 2.19711662180471e-55 | 7 | TPI1 |
| **PLXNC1.1** | 1.86886519281607e-59 | 0.311013591392069 | 0.504 | 0.212 | 2.36617022062443e-55 | 7 | PLXNC1 |
| **EIF3G.1** | 2.61483897283412e-59 | 0.258537902137384 | 0.393 | 0.144 | 3.31064762350527e-55 | 7 | EIF3G |
| **CDV3.1** | 3.12092089754211e-59 | 0.280468082142343 | 0.432 | 0.161 | 3.95139794837807e-55 | 7 | CDV3 |
| **EFHD1** | 3.96275358713634e-59 | 0.290917535904768 | 0.49 | 0.222 | 5.01724231667332e-55 | 7 | EFHD1 |
| **JUN.6** | 4.88199313826267e-59 | 0.605957146946823 | 0.804 | 0.277 | 6.18109151235436e-55 | 7 | JUN |
| **ALCAM.1** | 6.92764775947377e-59 | 0.258499472258698 | 0.33 | 0.086 | 8.77109482826974e-55 | 7 | ALCAM |
| **TMEM167A.1** | 4.86849068163003e-58 | 0.260950970605366 | 0.364 | 0.111 | 6.16399605201178e-54 | 7 | TMEM167A |
| **ND5.3** | 1.84769800516964e-57 | 0.426924561246962 | 0.982 | 0.835 | 2.33937044434528e-53 | 7 | ND5 |
| **LAPTM5.2** | 2.12487250083112e-57 | 0.337787716677601 | 0.732 | 0.467 | 2.69030107330228e-53 | 7 | LAPTM5 |
| **MYH9.2** | 2.98709840007919e-57 | 0.368706905833106 | 0.721 | 0.43 | 3.78196528434027e-53 | 7 | MYH9 |
| **RNASET2.1** | 2.3555073450288e-55 | 0.305499516112997 | 0.521 | 0.248 | 2.98230784954096e-51 | 7 | RNASET2 |
| **ITGB1.1** | 3.30998803350893e-55 | 0.295745583614724 | 0.466 | 0.178 | 4.19077584922566e-51 | 7 | ITGB1 |
| **EIF1.1** | 1.35356523394391e-54 | 0.304992539863244 | 0.578 | 0.336 | 1.71374894269638e-50 | 7 | EIF1 |
| **YWHAZ.1** | 1.22337752799419e-53 | 0.256757407288464 | 0.411 | 0.173 | 1.54891828819344e-49 | 7 | YWHAZ |
| **CCL4.1** | 4.637312707142e-53 | 0.606180693750481 | 0.566 | 0.127 | 5.87130161851248e-49 | 7 | CCL4 |
| **SEC61B** | 5.26641994704943e-53 | 0.265350199454591 | 0.435 | 0.187 | 6.66781429495929e-49 | 7 | SEC61B |
| **EIF3D.1** | 6.91416749794841e-53 | 0.250611341479374 | 0.394 | 0.153 | 8.75402746915248e-49 | 7 | EIF3D |
| **MYO1F.1** | 2.90369005730953e-52 | 0.275176266374704 | 0.473 | 0.207 | 3.6763619815596e-48 | 7 | MYO1F |
| **ENSGALG00000003348.1** | 3.06751610784477e-52 | 0.293783989516084 | 0.542 | 0.242 | 3.88378214414226e-48 | 7 | ENSGALG00000003348 |
| **KLF6.1** | 3.20035747599076e-52 | 0.375092353031704 | 0.495 | 0.147 | 4.0519726003519e-48 | 7 | KLF6 |
| **PPIA.2** | 4.25079982294752e-52 | 0.353794076618489 | 0.822 | 0.585 | 5.38193765583385e-48 | 7 | PPIA |
| **MCL1.2** | 1.03546268906125e-51 | 0.313884372296213 | 0.524 | 0.217 | 1.31099931062044e-47 | 7 | MCL1 |
| **NDFIP1** | 6.34384283083906e-51 | 0.276812432989904 | 0.542 | 0.303 | 8.03193940812533e-47 | 7 | NDFIP1 |
| **SGK1.2** | 5.10604862461147e-50 | 0.325541591165908 | 0.435 | 0.159 | 6.46476816362058e-46 | 7 | SGK1 |
| **ENSGALG00000045199.3** | 6.54967573719312e-50 | 0.353489430766933 | 0.56 | 0.239 | 8.29254445086021e-46 | 7 | ENSGALG00000045199 |
| **TMSB15B.6** | 1.19046508579304e-49 | -0.498358909048894 | 0.568 | 0.722 | 1.50724784512256e-45 | 7 | TMSB15B |
| **NDUFB2.1** | 7.43614677639672e-49 | 0.279792230483674 | 0.574 | 0.317 | 9.41490543359588e-45 | 7 | NDUFB2 |
| **ENO1.3** | 1.43655586084608e-48 | 0.3242173435595 | 0.668 | 0.388 | 1.81882337541723e-44 | 7 | ENO1 |
| **COX6A1.1** | 2.4956004448788e-48 | 0.317385183779858 | 0.673 | 0.413 | 3.15967972326105e-44 | 7 | COX6A1 |
| **ALDOC.1** | 3.95259768350534e-48 | 0.262704463393804 | 0.397 | 0.14 | 5.00438392708611e-44 | 7 | ALDOC |
| **HNRNPH2.1** | 7.74857304157419e-48 | 0.293485856897333 | 0.591 | 0.336 | 9.81046832793709e-44 | 7 | HNRNPH2 |
| **NPM1.1** | 1.53971499021666e-47 | 0.296184046404946 | 0.549 | 0.289 | 1.94943314911331e-43 | 7 | NPM1 |
| **HNRNPA3.1** | 2.60314452930332e-47 | 0.289989114282356 | 0.554 | 0.299 | 3.29584128855093e-43 | 7 | HNRNPA3 |
| **CTSC.4** | 4.45820087220639e-47 | 0.311956956643079 | 0.61 | 0.32 | 5.64452812430051e-43 | 7 | CTSC |
| **TMED10.1** | 1.18704459234221e-46 | 0.265827835744387 | 0.49 | 0.227 | 1.50291715836447e-42 | 7 | TMED10 |
| **SEPT7.1** | 5.09151085524461e-46 | 0.28201208595232 | 0.551 | 0.278 | 6.4463618938252e-42 | 7 | SEPT7 |
| **ENSGALG00000043329.1** | 9.51952977929197e-46 | 0.330485938847884 | 0.702 | 0.448 | 1.20526766535616e-41 | 7 | ENSGALG00000043329 |
| **ATP5F1D.1** | 1.26043202716434e-43 | 0.289386623819878 | 0.551 | 0.296 | 1.59583298959277e-39 | 7 | ATP5F1D |
| **CSF1R.4** | 3.81470956959222e-41 | 0.310614238520395 | 0.667 | 0.392 | 4.82980378606071e-37 | 7 | CSF1R |
| **SERBP1.1** | 1.5869250856092e-40 | 0.256266456396006 | 0.539 | 0.297 | 2.00920585088981e-36 | 7 | SERBP1 |
| **EIF4A2.1** | 3.03654448161421e-40 | 0.286515865984854 | 0.743 | 0.542 | 3.84456896817175e-36 | 7 | EIF4A2 |

| | | | | | | | |
|---|---|---|---|---|---|---|---|
| **CTSV.4** | 6.12891103335915e-40 | 0.317504613123196 | 0.566 | 0.262 | 7.75981425933602e-36 | 7 | CTSV |
| **TLN1.1** | 4.58630150808448e-39 | 0.251118842084587 | 0.469 | 0.239 | 5.80671633938576e-35 | 7 | TLN1 |
| **PPIB.3** | 1.12564468400338e-38 | 0.287534968684 | 0.606 | 0.341 | 1.42517873441668e-34 | 7 | PPIB |
| **ENSGALG00000005257.3** | 3.28760989813288e-38 | -0.350872204971815 | 0.085 | 0.339 | 4.16244289202604e-34 | 7 | ENSGALG00000005257 |
| **HSP90B1.5** | 3.82947367718099e-38 | 0.358633970472815 | 0.679 | 0.42 | 4.84849662267886e-34 | 7 | HSP90B1 |
| **GNS 0.30** | 7.37198124822692e-38 | 0.263485720876925 | 0.569 | 0.324 | 9.3336654583801e-34 | 7 | GNS |
| **ND4.3** | 2.81233331452172e-37 | 0.338189309363889 | 0.927 | 0.758 | 3.56069520951595e-33 | 7 | ND4 |
| **CD47.5** | 7.93846752041206e-37 | 0.306328552739409 | 0.647 | 0.337 | 1.00508937275937e-32 | 7 | CD47 |
| **ND1.3** | 7.99742979522486e-37 | 0.333233060626334 | 0.954 | 0.797 | 1.01255458637342e-32 | 7 | ND1 |
| **ENSGALG00000028749.3** | 3.32561253916713e-36 | 0.29758993757354 | 0.845 | 0.648 | 4.2105580358395e-32 | 7 | ENSGALG00000028749 |
| **ARPC4.2** | 3.74051834821863e-36 | 0.274491512523532 | 0.703 | 0.481 | 4.73587028067961e-32 | 7 | ARPC4 |
| **ENSGALG00000052354.2** | 1.85249492821272e-35 | 0.250705368781179 | 0.438 | 0.219 | 2.34544382861013e-31 | 7 | ENSGALG00000052354 |
| **DDX3X.3** | 3.78876523814282e-33 | 0.273435478623301 | 0.581 | 0.32 | 4.79695566801263e-29 | 7 | DDX3X |
| **PSAP.4** | 8.11767637839496e-33 | 0.282866699901984 | 0.609 | 0.353 | 1.02777900626859e-28 | 7 | PSAP |
| **ENSGALG00000047532.4** | 3.80432190635751e-32 | 0.254551146646405 | 0.584 | 0.343 | 4.81665196563924e-28 | 7 | ENSGALG00000047532 |
| **ENSGALG00000014585.6** | 6.03012374489746e-32 | -0.359574346533062 | 0.222 | 0.396 | 7.63473967341468e-28 | 7 | ENSGALG00000014585 |
| **CTSB.5** | 7.95054880008559e-32 | 0.294034004266606 | 0.621 | 0.358 | 1.00661898357884e-27 | 7 | CTSB |
| **PRDX1.3** | 2.00116049698616e-31 | 0.260452211364916 | 0.613 | 0.377 | 2.53366930523417e-27 | 7 | PRDX1 |
| **BLB1.6** | 4.57116699558544e-30 | 0.342359521345479 | 0.986 | 0.85 | 5.78755453311073e-26 | 7 | BLB1 |
| **PTPRC.4** | 1.52796727104606e-29 | 0.258182511206703 | 0.588 | 0.362 | 1.93455936187142e-25 | 7 | PTPRC |
| **NFKBIA.6** | 1.60894077918556e-28 | 0.310642508861756 | 0.606 | 0.313 | 2.03707992052683e-24 | 7 | NFKBIA |
| **ENSGALG00000052026.1** | 1.70695648983457e-27 | 0.44446095991345 | 0.443 | 0.082 | 2.16117761177955e-23 | 7 | ENSGALG00000052026 |
| **C1QA.5** | 4.05681443848333e-26 | -0.283155748351113 | 0.317 | 0.479 | 5.13633276056375e-22 | 7 | C1QA |
| **ENSGALG00000004509.6** | 1.12991136434943e-25 | 0.288111106102698 | 0.963 | 0.858 | 1.43058077840281e-21 | 7 | ENSGALG00000004509 |
| **S100A4.2** | 4.70397299075215e-24 | 0.847696320773881 | 0.763 | 0.153 | 5.95570020359129e-20 | 7 | S100A4 |
| **IFIT5.1** | 5.3752118319567e-24 | 0.317346145142538 | 0.42 | 0.203 | 6.80555570044037e-20 | 7 | IFIT5 |
| **HMOX1.6** | 2.25000905726825e-20 | -0.334083218067078 | 0.286 | 0.363 | 2.84873646740734e-16 | 7 | HMOX1 |
| **GRN.5** | 1.79635163124052e-19 | 0.251615652887753 | 0.843 | 0.611 | 2.27436080031362e-15 | 7 | GRN |
| **C1QB.6** | 1.59809851489816e-18 | -0.260473200825695 | 0.341 | 0.444 | 2.02335252971256e-14 | 7 | C1QB |
